# Infection-related images reduce cortical edge metastability: a specification-curve analysis of a block-design fMRI study

**DOI:** 10.64898/2026.10.27.762178

**Authors:** Jeyoon Choi, Tomas Berjaga-Buisan, Elvira Garcia-Guzman, Motoaki Sugiura, Gustavo Deco

## Abstract

Cues of infection evoke disgust, attentional capture, withdrawal and shifts in social orientation, a set of responses attributed to the behavioral immune system. Responses this varied have to be coordinated, as the effects of inflammation are, so they should be visible in whole-cortex coupling. We predicted that infection-related images hold cortical dynamics in a narrower repertoire, reducing cortical edge metastability, the temporal variability of mean cofluctuation across parcel pairs. In two fMRI runs, 44 adults viewed alternating 18-s blocks of infection-related and neutral images; the primary test was specified before we looked at the data. That test gave Cohen’s *d* = −0.31 (permutation *p* = .042), and 263 of 480 specifications were significant, all negative (joint *p < .*001). With global signal regression, the synchrony-based reduction survived removal of the stimulus-locked response, detrending, and motion, white-matter and cerebrospinal-fluid regression (*d* = −0.47), recurred in the second run (*d* = −0.44), and was smaller on strictly alternating blocks (*d* = −0.36). Without global signal regression, the pre-specified effect did not survive white-matter and cerebrospinal-fluid regression. Individual-difference predictions were not supported, and participants’ contrasts did not replicate across runs. Infection-related images reduce the variability of residual global cofluctuation at the group level; whether this is specific to pathogen cues remains open.

**Author Summary:** Cues of infection evoke disgust, attention, withdrawal and changes in social behavior, and responses this varied have to be coordinated, as the effects of inflammation are. We asked whether that coordination can be seen in whole-cortex coupling, predicting that infection-related images hold cortical dynamics in a narrower repertoire and so reduce edge metastability, the moment-to-moment variability of that coupling. The prediction held under the analysis we fixed in advance, and every significant result among 480 ways of computing it had the predicted sign. After removal of the global signal, the reduction survived controls for task-evoked responses, head motion and white-matter and cerebrospinal-fluid signals, and recurred in a second run; without that removal it did not. Individual differences in the effect did not replicate across runs.

## Introduction

Infectious disease has shaped two lines of defense. The physiological immune system acts once pathogens have entered the body, and its response is diverse but coordinated: inflammation produces sickness behavior, an adaptive withdrawal marked by fatigue, reduced appetite and social withdrawal ^1,2^, and prostaglandins made from arachidonic acid act on separate circuits to produce fever, anorexia and hyperalgesia as one syndrome ^3^. The behavioral immune system is proposed as a coarser first line that acts before contact: it detects pathogen cues, generates disgust, and biases cognition and behavior toward avoidance ^4–7^; the COVID-19 pandemic made its cues part of daily life worldwide ^8^. Pathogen cues raise disgust ^9–12^, whose expression narrows the visual field and nasal inspiration that fear expands ^13^. They also hold attention ^14^, lower self-rated extraversion^15^, and shift social orientation, toward collectivism where pathogens are prevalent^16,17^ and toward horizontal individualism after induced infection threat^18^. Responses this varied must be coordinated, as the effects of inflammation are, and from this we draw a working hypothesis: cues of infection, like inflammation itself, shift the organism as a whole toward withdrawal and a narrowed focus. We call this a defensive mode; engagement here means the breadth of the organism’s interaction with its environment, whereas attention to the cue itself should increase.

Neuroimaging has mapped components of this response, the anterior insula in disgust ^19^ and regions responding to pathogen-related acts and sick conspecifics^20,21^, but a coordinated response with many components is a property of the cortex as a whole, and a regional response shows where a cue is processed without showing whether the cortex has changed its mode. Cortical regions form and dissolve coalitions, and metastability, the temporal variability of global synchrony, indexes how widely the system moves through this repertoire ^22,23^; after traumatic brain injury it falls with structural damage and lost cognitive flexibility ^24^. Coordination implies less variability, since a cortex that settles into one configuration for a time varies less in its coupling, and threat sits at this low end. Acute stress increases coupling within the salience network ^25^, imminent threat narrows behavior to a few defensive responses ^26–28^, and an attention-demanding task raised global synchrony and lowered metastability ^29^, although task effects have differed in direction between studies^30^. If cues of infection coordinate a withdrawal as sickness does, cortical dynamics should move through less of their repertoire while the cues are present. This is our primary prediction: infection-related images reduce cortical edge metastability. Two accounts without defensive withdrawal make the same prediction: a salient stimulus lowers metastability ^29^, and global signal amplitude, which dominates the measure, falls as vigilance rises^31^. The design tests the direction of the effect, not its specificity.

Edge time series make this measurable. The product of two parcels’ z-scored signals at each time point is their instantaneous cofluctuation^32–34^, and the mean over all pairs, the global cofluctuation *R*(*t*), tracks how synchronized the cortex is. Sliding-window connectivity averages over tens of seconds and depends on window length and shape^35^; edge time series resolve cofluctuation at every volume, so its variability can be measured within 18-s blocks. Edge metastability, the temporal standard deviation of *R*(*t*), is the amplitude counterpart of the variability of the squared order parameter in phase-based models. We use the term as a name for this quantity, computed from amplitude cofluctuation over short task windows, without assuming that it indexes the same property as phase-based metastability; a reduction is compatible with a narrowed repertoire but does not demonstrate one.

The behavioral immune system framework also predicts who should show the shift most; these secondary and exploratory predictions were specified after the primary result was known. Pathogen disgust is proposed to be stronger in women, for whom infection carries greater reproductive costs ^36^, and neural responses to disgusting images differ by sex ^37^. The secondary predictions are a larger reduction in women and a smaller reduction, habituation ^38^, in a second, identical run. Because infection threat increased horizontal individualism in our behavioral work^18^, trait horizontal individualism is an exploratory moderator without a directional prediction, and the pre–post change in individualism and the in-scanner ratings are candidate state correlates. All of these assume that a participant’s contrast is stable, which two consecutive runs let us test.

In short task blocks the measure depends on window placement, definition, normalization and global signal regression. Such flexibility is a recognized source of unstable results in task fMRI ^39,40^, and the remedy is to report how results vary over the reasonable choices ^41,42^. We therefore fixed the primary test before looking at the data, evaluated it over 480 specifications, compared it with covariance-matched surrogates, tested whether it survived removal of task-evoked responses and nuisance signals and recurred in the second run, and then tested the secondary and exploratory predictions.

## Results

### The stimulus manipulation worked as intended

In run 1, infection-related images were rated higher than neutral images on disgust (*M* = 2.94 versus 1.17 on a 4-point scale, *t*(43) = 22.6, *d* = 3.40), fear (2.72 versus 1.14, *t*(43) = 16.8, *d* = 2.53) and perceived infection risk (2.81 versus 1.33, *t*(43) = 15.1, *d* = 2.28; all *p < .*001). The neutral images scored low on all three dimensions. Because the design had no negatively valenced non-pathogen condition, the contrast separates infection-related images from neutral ones but cannot separate pathogen specificity from negative arousal (Discussion). Figure 1 shows the task and the computation of edge metastability for one participant.

**Figure 1:**
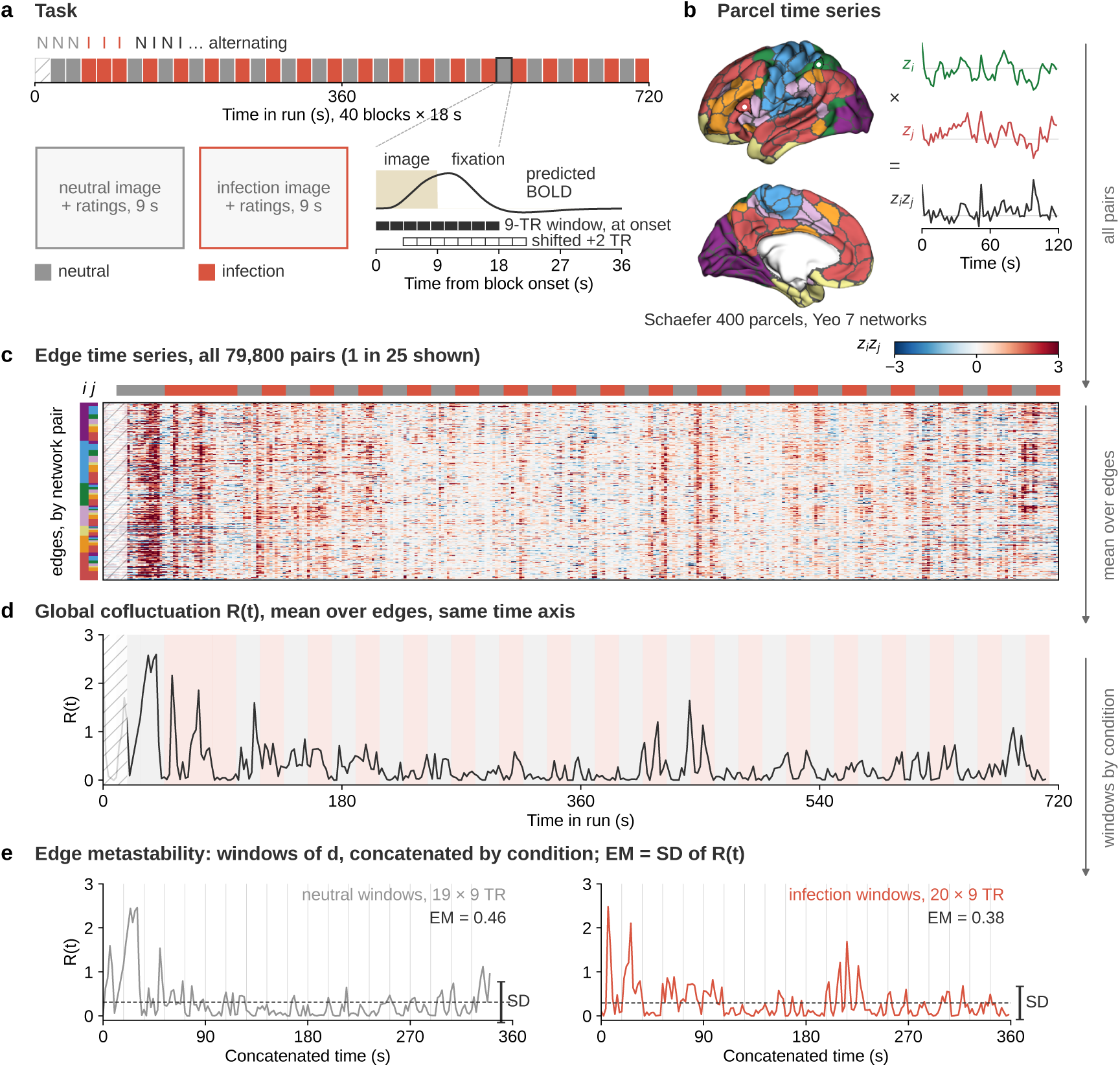
From stimulus blocks to edge metastability, illustrated with one participant’s run. Terms: TR, repetition time (2 s); BOLD, blood-oxygen-level-dependent signal. (a) Task. Top: the 40 blocks of one run (18 s each; orange infection, gray neutral), one image per block. The order is fixed: three neutral, three infection, then alternating. The first block (hatched here and in c, d) is excluded because its window overlaps the discarded volumes, leaving 19 neutral and 20 infection blocks. Bottom left: gray placeholders mark where one infection-related and one neutral image appeared; the images themselves are not reproduced because their licenses do not permit publication under CC BY. Bottom right: one block (outlined above), one image shown for 9 s while it is rated, followed by 9 s of fixation, with the predicted hemodynamic response and the 9-TR analysis window placed at block onset (filled, pre-specified) or shifted by 2 TR (open). (b) BOLD is parcellated (Schaefer 400, left hemisphere, lateral and medial views, parcel boundaries in gray, colored by Yeo network as in Figure 5; atlas sources in Methods) and z-scored; the element-wise product of two parcel time series, a dorsal-attention parcel *i* and a default-mode parcel *j* (marked on the surface), is one edge time series (first 120 s shown). (c) All 79,800 edge time series of the run, every 25th shown, sorted by network pair (strips at left give the networks of *i* and *j*); vertical bands are moments of brainwide cofluctuation. (d) Global cofluctuation *R*(*t*), the mean over edges, on the same time axis, with block windows shaded. (e) Edge metastability: the windows of each condition are concatenated and z-scored, *R*(*t*) recomputed (thin vertical lines separate windows), and edge metastability (EM) is its standard deviation (dashed line: mean; bar at right: mean *±* SD). The two values are this participant’s (participant 1, run 1) entries in the data release.

### The pre-specified test supports the predicted reduction

The pre-specified test (synchrony-based edge metastability, the standard deviation over time of the global cofluctuation *R*(*t*) over the concatenated blocks of a condition; 9-TR windows at block onset, where TR is the repetition time of 2 s; normalization within condition; Schaefer 400; no global signal regression, GSR; run 1; two-tailed) gave lower edge metastability (EM) during infection than during neutral blocks in 27 of 44 participants, with a mean infection minus neutral difference ΔEM = −0.059, Cohen’s *d* = −0.31 (95% CI −0.61 to −0.01), *t*(43) = −2.06, *p* = .045, and a two-tailed sign-flip permutation *p* = .042 (10,000 draws; Figure 2a–c). A block-level linear mixed model, whose dependent variable is the metastability of each 9-TR window on its own (a quantity related to the within-block component; Methods; Supplementary Table S5), gave a descriptive estimate in the same direction (*b* = −0.021, *z* = −3.02, *p* = .003; with block-wise framewise displacement (FD) as a covariate, *b* = −0.019, *p* = .008). Block-level metastability also rose over the run (*b* = 0.013 per ten blocks, *z* = 4.28, *p < .*001), so the model includes block order as a covariate; the fixed block order is taken up below and in the Discussion.

**Figure 2:**
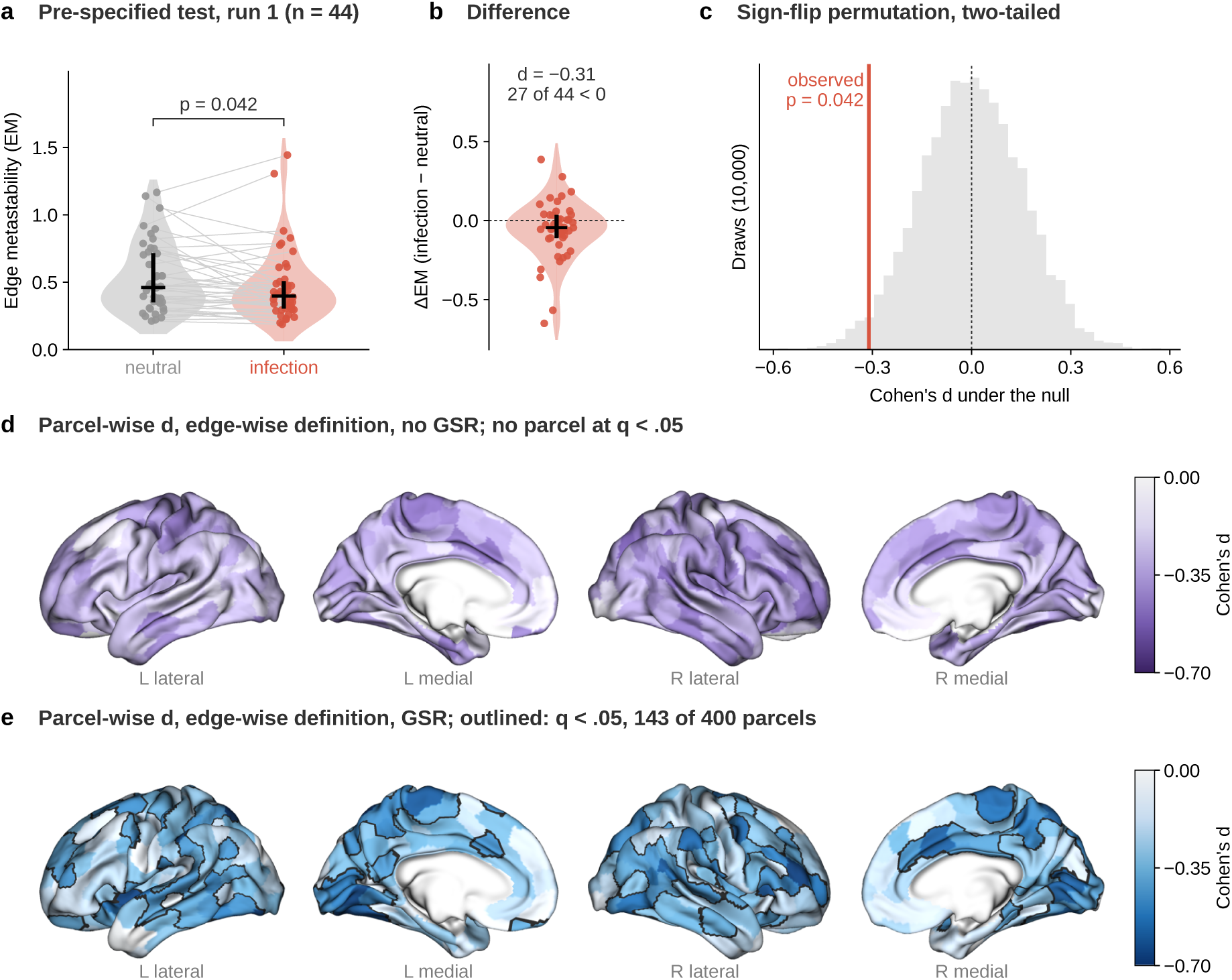
Infection-related images reduce edge metastability. (a) Synchrony-based edge metastability for neutral and infection blocks in run 1 under the pre-specified analysis (*n* = 44); violins show the distribution, points are participants, lines connect the two conditions within a participant, bars mark the median and interquartile range, and the bracket gives the two-tailed permutation *p*. (b) The within-participant difference ΔEM (infection minus neutral) for each participant; values below zero mean lower edge metastability during infection blocks, and the text gives Cohen’s *d* and the number of participants with a reduction. (c) Sign-flip permutation null distribution of Cohen’s *d* (the 10,000 draws used for the test) with the observed *d* and its two-tailed *p*. (d) Parcel-wise Cohen’s *d* for the edge-wise definition without global signal regression (GSR; both hemispheres, lateral and medial views); no parcel reaches *q < .*05 (Benjamini–Hochberg false discovery rate across 400 parcels), so none is outlined. (e) The same after GSR; 143 outlined parcels at *q < .*05, all reductions. In d and e and in Figure 5, purple shades mark specifications without GSR and blue shades specifications with GSR; the two scales are matched in lightness, so equal *d* has equal shade. Color is clipped to [−0.7, 0]; the three parcels in d and the one in e with a small positive *d* (at most 0.06) appear white. The edge-wise effects did not survive detrending or nuisance regression (Supplementary Table S17), so both maps are descriptive.

### The reduction holds across 480 specifications and is largest after global signal regression

We then evaluated the same contrast across 480 specifications (160 in the plan as written; Methods): window shift of 0 to 3 TR after onset; four definitions of edge metastability (synchrony-based; its within-block and between-block components, the mean within-window variability of *R*(*t*) and the variability of the window means; and edge-wise, the mean over parcel pairs of each edge’s own temporal standard deviation); normalization within condition or within run; Schaefer parcellations at 100, 200, 400, 600, 800 and 1000 parcels; GSR absent or present; and all blocks or a common set of 19 blocks per condition. Figure 3a shows the sorted effect sizes (all values in Supplementary Table S2). Across specifications the median *d* was −0.34 (range −0.83 to +0.07). Of the 480 specifications, 263 were significant at two-tailed *p < .*05, all in the negative direction. The specifications are strongly dependent, since most share the data and differ in one choice, so the count is descriptive; the inference comes from joint sign-flip permutation with flips shared across specifications, which rejected the null for the median effect, for the number of significant negative specifications and for the sum of significant effects (all *p < .*001; Supplementary Table S3).

**Figure 3:**
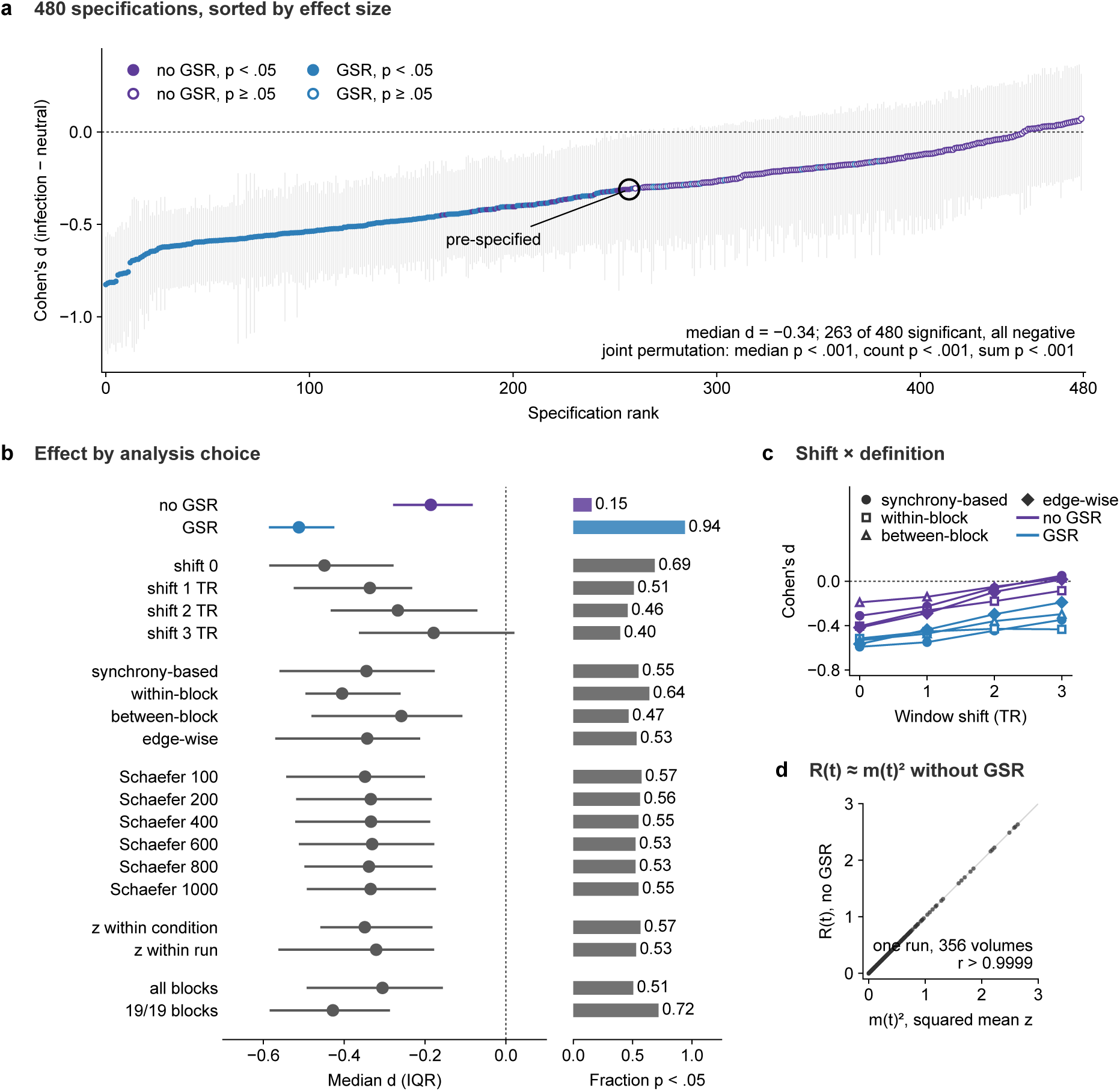
Specification curve for the infection minus neutral contrast in run 1. (a) Cohen’s *d* for 480 specifications sorted by effect size, with bootstrap 95% intervals (2,000 resamples); colors as in Figure 2 (blue, with GSR; purple, without), filled markers are two-tailed permutation *p < .*05, and the black ring marks the pre-specified analysis. (b) For each level of each dimension, the median and interquartile range of *d* across the specifications that use it (left) and the fraction of those specifications with *p < .*05 (right). (c) Effect size as a function of window shift for the four definitions, without (purple) and with (blue) GSR (Schaefer 400, normalization within condition); shift 0 corresponds to the decomposition reported in the text. (d) In one run without GSR (participant 1, 356 volumes) the global cofluctuation *R*(*t*) is approximately the squared mean of the z-scored parcel signals, *m*(*t*)^2^, so its temporal variability is the variability of the power of the parcel-mean signal.

Summarized by dimension (Figure 3b; the full curve with its indicator matrix is Supplementary Figure S1), one choice, GSR, organizes most of the curve. With GSR, 226 of 240 specifications were significant (median *d* = −0.51); without GSR, 37 of 240 (median *d* = −0.19). Parcellation resolution made no difference; the median *d* lay between −0.33 and −0.35, and 42 to 46 of 80 specifications were significant, at every resolution from 100 to 1000 parcels, so the six resolutions add little independent information. Window shift mattered (shift 0: 132 of 192, median −0.45; shift 3: 38 of 96, median −0.18), and normalization within condition or within run made little difference (136 versus 127 of 240). Across the four definitions, the median *d* was −0.35 for the synchrony-based measure, −0.41 for its within-block component, −0.26 for its between-block component and −0.34 for the edge-wise measure. The block set made little difference except at shift 0 for the run-normalized edge-wise GSR specifications, where dropping the final block, an infection block whose window ends at the last retained volume, moved *d* from −0.62 to −0.71 (all blocks) to −0.81 to −0.83 (19 per condition); the largest values in the curve therefore depend on one block.

Without GSR the effect on the pre-specified measure weakened with window shift (Figure 3c), from *d* = −0.31 at onset to −0.23 at a 1-TR shift, −0.06 at 2 TR and +0.05 at 3 TR. It was present only when the window included the block transition and the response to the preceding block, and absent at the shifts that align the window with the block’s own hemodynamic response. With GSR and normalization within condition, the synchrony-based measure declined from −0.59 at onset to −0.35 at 3 TR (all *p ≤ .*022), whereas the edge-wise measure fell from −0.56 to −0.19 (*p* = .22 at 3 TR) and at 2 TR did not survive motion adjustment (below). With normalization within run, the edge-wise GSR measure stayed between −0.57 and −0.77 (Schaefer 400) and between −0.55 and −0.76 (Schaefer 200) at every shift with all blocks (all *p < .*001; Supplementary Table S2). Part of this effect is a lower signal variance over the concatenated windows, and these run-normalized specifications were not tested against detrending or nuisance regression (below). After GSR, part of the effect persists at windows aligned with the hemodynamic response; once nuisance signals are removed, it does so only for the synchrony-based measure (below).

### What the contrast contains

*R*(*t*) depends only on the mean *m*(*t*) and the mean square *q*(*t*) of the z-scored parcel signals (Methods, Equation 1), and its first term dominates whenever *N* Var*_t_*[*m*(*t*)], with *N* the number of parcels, is large compared with Var*_t_*[*q*(*t*)]. In run 1 at Schaefer 400 it was: *N* Var*_t_*[*m*(*t*)] = 122 without GSR and 45 after GSR, against Var*_t_*[*q*(*t*)] = 0.45 and 0.23 (means over participants, averaged over the two conditions; per-condition values in Supplementary Table S16). *R*(*t*) is therefore close to *m*(*t*)^2^, the instantaneous power of the cortical parcel-mean signal, with and without GSR (Figure 3d), and the synchrony-based measure is its temporal variability. *R*(*t*) depends only on *m*(*t*) and *q*(*t*) and carries no information about which pairs cofluctuate; only the edge-wise definition does.

For jointly Gaussian signals every definition used here is a function of the static correlation matrix of the concatenated windows, so a condition difference in edge metastability could be a difference in static connectivity. We tested this against covariance-matched surrogates: for each participant and condition, Gaussian time series with the same static covariance as the concatenated windows but no temporal structure. Without GSR, the mean pairwise correlation, which equals the mean of *R*(*t*) over the concatenated windows up to a factor (*T −* 1)*/T*, did not differ between conditions (0.299 versus 0.305; *d* = −0.08, *p* = .60), the surrogates reproduced none of the effect (surrogate ΔEM *d* = −0.02, *p* = .88), and the observed contrast exceeded the surrogate contrast (*d* = −0.42, *p* = .006). With GSR, the residual mean correlation was lower during infection (*d* = −0.54, *p* = .001), and the surrogates reproduced about half of the synchrony-based effect (surrogate *d* = −0.51; observed minus surrogate *d* = −0.50, *p < .*001) but none of the edge-wise effect (surrogate *d* = −0.17, *p* = .26; observed minus surrogate *d* = −0.55, *p < .*001). Without GSR, then, the reduction is not a difference in static connectivity, and with GSR about half of it is not. The difference lies in the temporal structure. The standard deviation over time of *q*(*t*), the instantaneous global power, was lower during infection with and without GSR (*d* = −0.46, *p* = .003 without GSR; *d* = −0.60, *p < .*001 with GSR), which means that infection blocks contained fewer or smaller brain-wide bursts of cofluctuation ^33^. The mean within-block variance of the run-normalized parcel signals did not differ between conditions (*d* = −0.04, *p* = .77 without GSR; *d* = +0.05, *p* = .75 with GSR), so a uniform change in signal amplitude does not explain the effect; after GSR, however, the variance of the run-normalized signals over the concatenated windows, which includes block-to-block level shifts, was lower during infection (*d* = −0.58, *p < .*001), and this contributes to the run-normalized GSR specifications that gave the largest effects. The dispersion of parcel variances across parcels was higher during infection after GSR (*d* = +0.41, *p* = .010) and unchanged without GSR (*d* = +0.07, *p* = .65), so the GSR residuals were less uniform in amplitude during infection blocks (Supplementary Table S16).

To see how the pre-specified measure distributes over time scales, we decomposed its variance into the mean within-block variability of *R*(*t*) and the variability of the block means (shift 0 in Figure 3c; Supplementary Table S6). The within-block component was reduced (*d* = −0.40, *p* = .007) whereas the between-block component was not (*d* = −0.19, *p* = .22), and the two components ranked participants differently (*r* = −0.19 across participants between the two ΔEM values; Supplementary Figure S2). The edge-wise definition gave *d* = −0.42 (*p* = .007) under the same conditions. Participant rankings also differed between specifications with and without GSR: the ΔEM values of the pre-specified analysis correlated *r* = 0.24 with the synchrony-based GSR measure and *r* = −0.21 with the edge-wise GSR measure (Supplementary Figure S2). So although the specifications agree at the group level, they rank individual participants almost independently of one another.

Because the block order was fixed, with three neutral and three infection blocks before strict alternation began, we recomputed the contrast on the strictly alternating segment (blocks 7 to 40, 17 blocks per condition). The effects were smaller: without GSR, synchrony-based *d* = −0.27 (*p* = .087) and edge-wise *d* = −0.31 (*p* = .042); with GSR, synchrony-based *d* = −0.30 (*p* = .049) and edge-wise *d* = −0.32 (*p* = .039) (Supplementary Table S16). Part of this loss reflects the smaller number of blocks (17 rather than 19 or 20 per condition). The rest may come from the opening, non-alternating blocks, but the design cannot separate that contribution from sampling variability.

FD was slightly lower during infection than during neutral blocks (difference = −0.007 mm, *t*(43) = −2.51, *p* = .016), and the condition difference in FD correlated with ΔEM (*r* = −0.34, *p* = .023) in a direction that works against the effect between participants, since those who moved less during infection than during neutral blocks showed a smaller reduction. Within participants, motion favors the effect, because block-wise FD predicts higher block-level metastability (*b* = 0.38 per mm, *z* = 7.09) and FD was lower in infection blocks; the condition estimate changed little when FD entered the model (−0.021 to −0.019), because the FD difference is small. Excluding the quartile with highest mean FD left the pre-specified effect intact (*d* = −0.43, *p* = .016, *n* = 33). Under GSR, the condition difference in FD correlated positively with the edge-wise ΔEM (*r* = 0.33 at Schaefer 400, *r* = 0.31 at Schaefer 200), so we re-estimated those effects at zero FD difference by regression. The onset-aligned GSR effects remained significant (adjusted *p* = .002 to .007), as did the 2-TR-shifted synchrony-based GSR effects (*p* = .028 and .026); the 2-TR-shifted edge-wise GSR effects did not (*p* = .27 and .26 at Schaefer 400 and 200; Supplementary Table S4). In contrast, regression of 24 motion parameters, a stricter control, removed the onset-aligned edge-wise GSR effect (*d* = −0.25, *p* = .11) but not the synchrony-based one (next section).

### After GSR the synchrony-based reduction survives control for evoked responses and nuisance signals and recurs in run 2

The specification curve varied analytic choices but did not remove task-evoked responses, which inflate connectivity estimates in task fMRI ^43^, or nuisance signals from motion and physiology, which bias them ^44^. We therefore re-estimated the four central specifications (synchrony-based and edge-wise, with and without GSR; Schaefer 400, normalization within condition, all blocks, run 1) at window shifts of 0 and 2 TR under six variants (Methods; Supplementary Table S17). The first subtracts the condition-mean response at each time point of the window, which is nearly identical to finite impulse response regression and removes the stimulus-locked component^43^. Three remove nuisance signals one at a time: linear and quadratic trends, 24 head-motion parameters^45^, and the mean white-matter and cerebrospinal-fluid (CSF) signals. The last two apply the three nuisance steps together, which we call nuisance regression, without and with the first operation.

Removing the stimulus-locked response changed the effects little (without GSR, synchrony-based *d* = −0.27, *p* = .083, and edge-wise *d* = −0.38, *p* = .015; with GSR, *d* = −0.54, *p < .*001, and *d* = −0.50, *p* = .001), so the reduction is not the mean evoked response. The synchrony-based reduction after GSR survived every nuisance step: detrending (*d* = −0.55, *p < .*001), motion regression (*d* = −0.42, *p < .*001), white-matter and CSF regression (*d* = −0.35, *p* = .014) and nuisance regression (*d* = −0.47, *p < .*001; *d* = −0.44, *p* = .002 with the evoked response also removed), and it remained significant after Holm–Bonferroni correction across the six variants at each shift. At a 2-TR shift, which aligns the window with the block’s own hemodynamic response, it was *d* = −0.57 after nuisance regression (*p < .*001). Without GSR, motion regression left the prespecified effect unchanged (*d* = −0.33, *p* = .033) and detrending weakened it slightly (*d* = −0.28, *p* = .067), but white-matter and CSF regression removed it (*d* = −0.10, *p* = .51). The edgewise reduction did not survive white-matter and CSF regression (*d* = −0.18, *p* = .25, with GSR; *d* = −0.10, *p* = .52, without) or detrending (*d* = −0.19, *p* = .22, with GSR; *d* = −0.27, *p* = .075, without), and with GSR it did not survive motion regression (*d* = −0.25, *p* = .11). The edge-wise GSR effect therefore depends on slow drift and motion-related variance (the run-normalized edgewise specifications that gave the largest effects in the curve were not tested). Of the four forms of the effect, only the synchrony-based reduction after GSR withstands these controls, and it is also the one present at hemodynamically aligned windows. The pre-specified effect does not survive removal of white-matter and CSF signals.

Because this form became the main result only after the controls had been run, we tested it in run 2, which played no part in selecting it, and against two alternative explanations (Figure 4; Supplementary Table S21). In run 2 (*n* = 43; Methods), the synchrony-based reduction after GSR and nuisance regression was *d* = −0.44 (*p* = .005) at onset and *d* = −0.29 (*p* = .063) at a 2-TR shift; with the contrast averaged over the two runs it was *d* = −0.65 and −0.60 (both *p < .*001). Without GSR, nuisance regression left no effect in run 2 either (*d* = −0.12), and the edge-wise measure after nuisance regression was null in both runs (*|d| ≤* 0.17). A single global-signal coefficient per parcel could turn a condition difference in global signal variance into a difference in the residuals; fitting a separate coefficient for samples in infection windows, in neutral windows and elsewhere left the regressed effect unchanged (*d* = −0.47 and −0.56 at shifts of 0 and 2 TR in run 1, *d* = −0.42 and −0.35 in run 2; all *p ≤ .*028), whereas it halved the onset-aligned edge-wise GSR effect before nuisance regression (*d* = −0.56 to −0.28, *p* = .072), so part of that effect does arise from the shared fit. Restricted to the strictly alternating blocks 7 to 40, it was smaller: *d* = −0.36 (*p* = .021) at onset and *d* = −0.27 (*p* = .080) at 2 TR in run 1, *d* = −0.26 (*p* = .095) at onset in run 2, and *d* = −0.38 (*p* = .018) at onset with the runs averaged. The regressed reduction therefore recurs across runs and does not depend on how the global signal was fitted. Part of it, however, comes from the opening, non-alternating blocks, and once those are removed it is no longer established at hemodynamically aligned windows.

**Figure 4:**
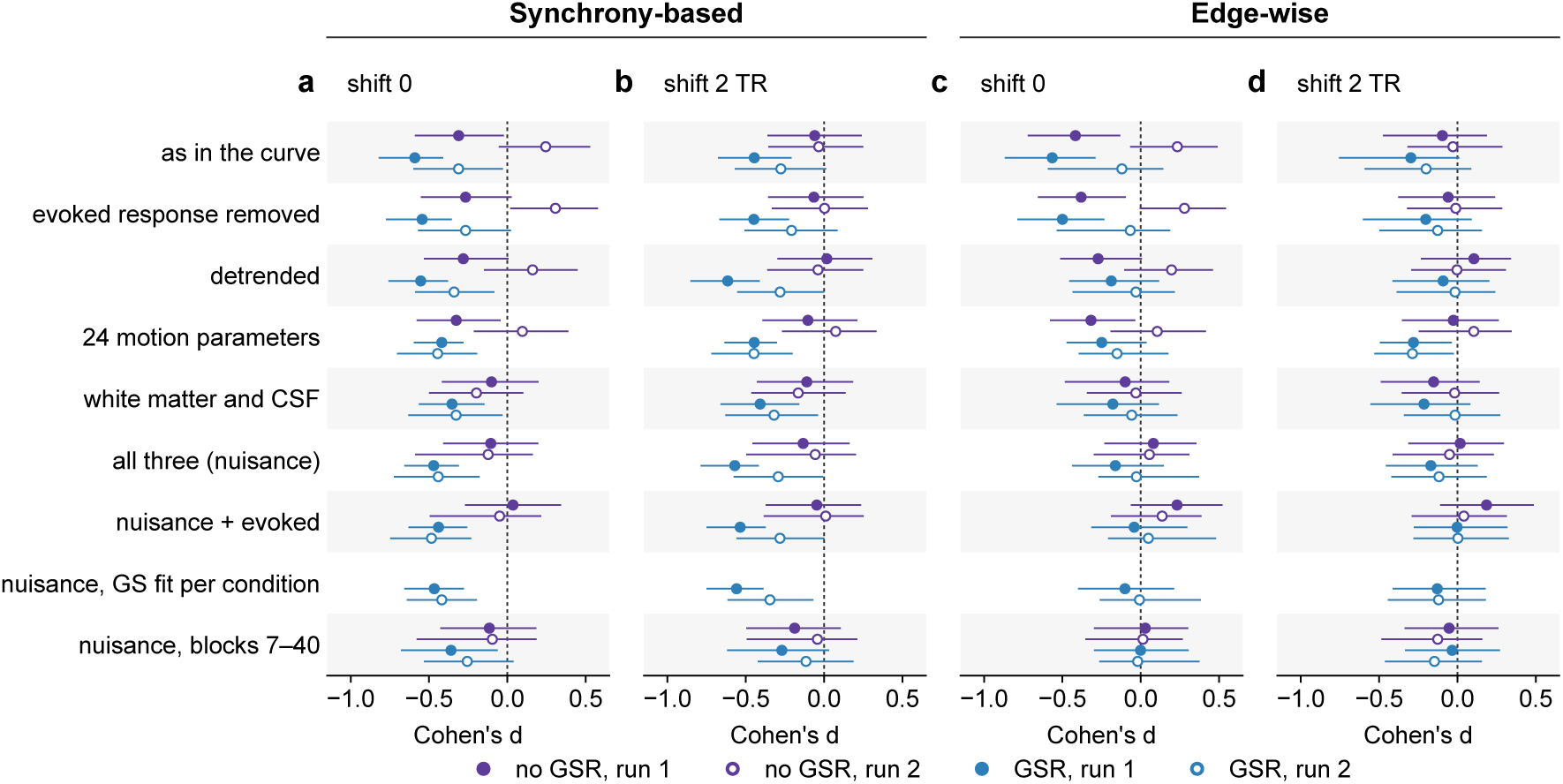
Control analyses and replication in run 2. Cohen’s *d* of ΔEM (infection minus neutral) with bootstrap 95% intervals (5,000 resamples) for the synchrony-based (a, b) and edge-wise (c, d) definitions at window shifts of 0 (a, c) and 2 TR (b, d), Schaefer 400, normalization within condition, without (purple) and with (blue) global signal regression (GSR), in run 1 (filled, *n* = 44) and run 2 (open, *n* = 43). Rows: the contrast as in the specification curve; after subtracting the condition-mean stimulus-locked response; after removing linear and quadratic trends; after regressing 24 motion parameters; after regressing the mean white-matter and cerebrospinal-fluid (CSF) signals; after all three nuisance steps (nuisance regression); after nuisance regression and evoked-response removal; after nuisance regression with the global signal fitted separately for infection windows, neutral windows and the remaining samples (GSR only); and after nuisance regression on the strictly alternating blocks 7 to 40. Values in Supplementary Tables S17 and S21.

### After GSR the reduction extends beyond visual cortex

Figure 2d,e maps the change in edge-wise metastability parcel by parcel (mean over each parcel’s edges). Without GSR, 397 of 400 parcels had a negative mean difference (median *d* = −0.31) but none reached *q < .*05 (Benjamini–Hochberg false discovery rate, BH-FDR, across parcels; smallest *q* = .055). The map is uniform and weak, which is what one expects of a measure dominated by the global signal. With GSR, 143 of 400 parcels showed a reduction at *q < .*05 and none an increase; because the edge-wise effects did not survive detrending or nuisance regression (above), both maps are descriptive. The 143 parcels were spread over every network and nearly absent from the limbic network (visual 29 of 61 parcels, ventral attention 24 of 47, dorsal attention 23 of 46, default mode 27 of 91, somatomotor 20 of 77, control 18 of 52, limbic 2 of 26), and the largest effects (*d* = −0.60 to −0.70) were in early visual cortex, posterior dorsal-attention parcels and the frontal operculum, with single parcels in the control, default-mode and somatomotor networks. At the network level (Figure 5), the pre-specified analysis gave a reduction at *q < .*05 only in the somatomotor network (*d* = −0.51, *q* = .011), with visual (*d* = −0.36, *q* = .072) and ventral attention (*d* = −0.32, *q* = .096) next and limbic smallest (*d* = −0.04). After GSR six of seven networks were reduced at *q < .*05 (visual *d* = −0.67; ventral attention −0.51; control −0.42; default mode −0.39; dorsal attention −0.39; somatomotor −0.35), with limbic again the exception (*d* = −0.05, *q* = .74; Supplementary Table S14). Because the effect after GSR was largest in visual cortex and the two image sets differed on several low-level properties (Supplementary Table S1), we recomputed the GSR contrasts with the 61 visual parcels excluded: synchrony-based *d* = −0.56 (*p < .*001) and edge-wise *d* = −0.53 (*p < .*001), close to the full-cortex values (Supplementary Table S16). Before nuisance regression, then, the reduction was distributed across sensorimotor and association cortex and did not depend on the visual parcels, although a stimulus-driven contribution within visual cortex cannot be excluded. After nuisance regression it was narrower (below).

**Figure 5:**
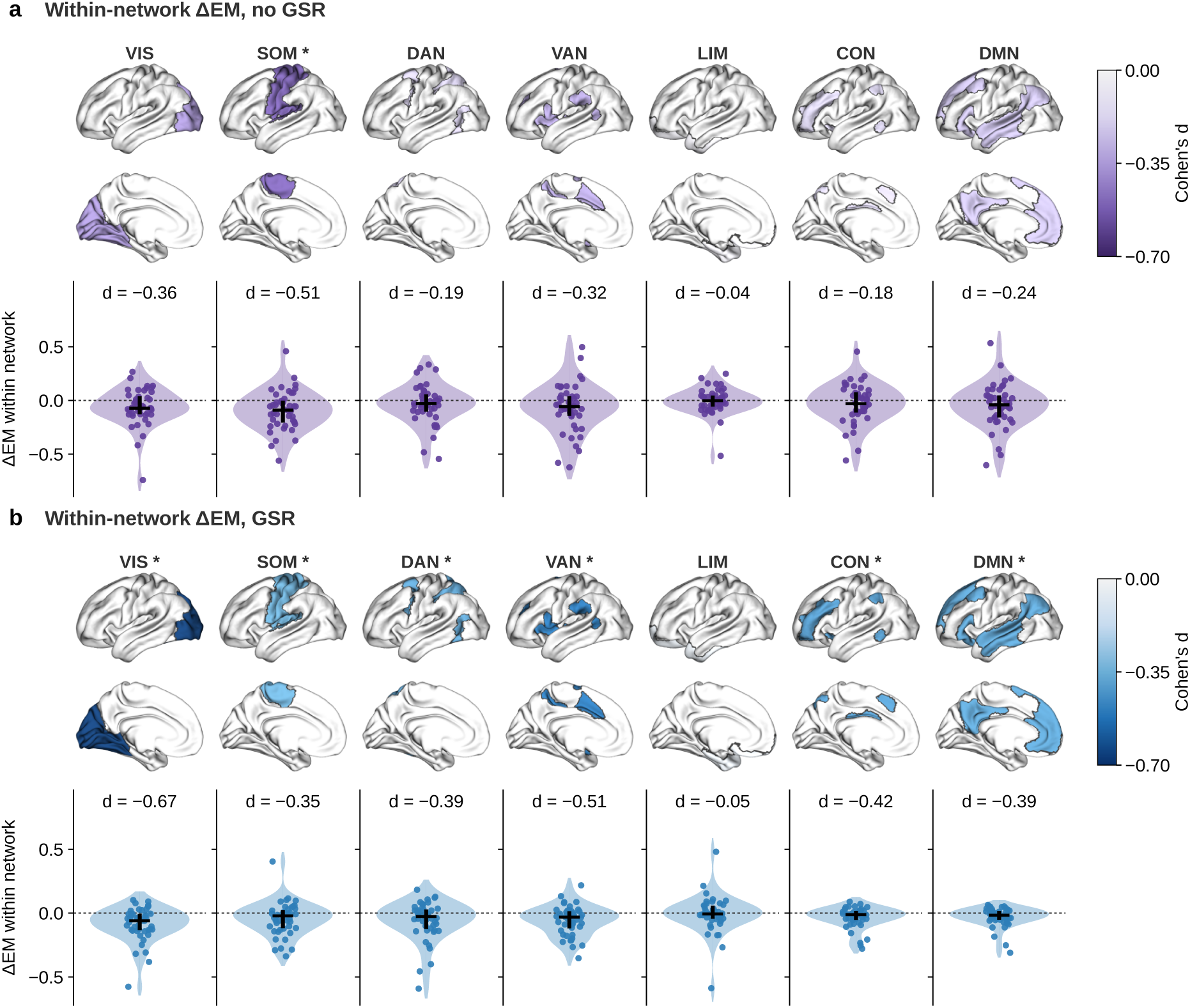
Network-level reduction with and without global signal regression. Change in within-network synchrony-based edge metastability (ΔEM, infection minus neutral) for the seven Yeo networks (a) without and (b) with global signal regression (GSR). Surfaces (left hemisphere, lateral and medial views) show each network outlined and filled by its Cohen’s *d*, colors as in Figure 2; asterisks mark *q < .*05 (Benjamini–Hochberg false discovery rate, BH-FDR, across seven networks). Violins show per-participant ΔEM with median and interquartile range; *d* is given in each panel. Values after nuisance regression are in Supplementary Table S19.

Participants responded with the right hand and rated infection-related images higher (mean rating 2.82 versus 1.21), so the somatomotor reduction might reflect finger use. Participants answered slightly fewer prompts during infection blocks in run 1, the direction a motor contribution would take (2.94 versus 2.98 of three per block, about one response in 80 prompts; *t*(43) = −3.47, *p* = .001) but not in run 2 (*p* = .12; Supplementary Table S18). Without GSR the somatomotor reduction was bilateral (left hemisphere *d* = −0.43, right −0.51; difference *p* = .59), whereas a contribution of right-hand responses should be strongest on the left. With GSR it was larger on the left (−0.47 versus −0.22), the direction a motor contribution would take, but the difference was not significant (*p* = .21; Supplementary Table S19). Like the whole-cortex effect without GSR, the somatomotor reduction disappeared after nuisance regression (*d* = −0.07 without and −0.14 with GSR). After nuisance regression without GSR, only the visual network retained a reduction (*d* = −0.44, *q* = .038); with GSR, the visual (*d* = −0.55), ventral attention (−0.50) and dorsal attention (−0.41) networks did (all *q < .*05), and the other four did not. The remaining visual reduction was larger in the left than in the right hemisphere (difference *p* = .004 without and *p* = .024 with GSR; Supplementary Table S19).

### The individual-difference predictions depend on the specification

The sex prediction was nominally supported only under specifications dominated by the global signal. Under the pre-specified analysis in run 1, women showed a reduction (*d* = −0.61, *n* = 21) and men none (*d* = 0.00, *n* = 23); the between-sex comparison of ΔEM, the first of two secondary tests corrected with the Holm–Bonferroni procedure, gave *t*(42) = −2.28, *p* = .027, Holm *p* = .055, and did not survive correction. The plan specified this test as a participant-level mixed model, which gave *z* = −2.34, *p* = .019 (Holm *p* = .039; Supplementary Table S7); with two values per participant that model is the same comparison, and its Wald *z* ignores the loss of degrees of freedom and is anticonservative at *n* = 44, so we report the *t* test. For that model, the Bayes factor approximated from the Bayesian information criterion (BIC; Methods) gave only weak support for the interaction (BF_01_ = 0.71, that is, BF_10_ = 1.4), and the block-level model showed no condition by sex interaction (*b* = 0.005, *p* = .70; Supplementary Table S5). The condition by run interaction, the direct test of habituation, was not significant (*b* = +0.102, *z* = 1.84, *p* = .066; BF_01_ = 2.5), and the run 2 contrast alone was not significant under the pre-specified analysis (*d* = +0.25, *p* = .12, *n* = 43).

Across six specifications (Supplementary Figure S3), the sex difference was nominally present without GSR (synchrony-based *p* = .027, edge-wise *p* = .047, uncorrected). After GSR the reduction appeared in both sexes (synchrony-based: women *d* = −0.68, men *d* = −0.57, sex difference *p* = .46). Under the edge-wise GSR specification it was nominally larger in men (women −0.30, men −0.78, uncorrected *p* = .023), a difference of the opposite sign, and the between- and within-block components of the pre-specified measure also carried sex differences of opposite sign, the within-block one not significant (Supplementary Table S6). In run 2 the synchrony-based GSR specifications gave negative contrasts of about half the run 1 size (Schaefer 400: *d* = −0.31, *p* = .048; Schaefer 200: *d* = −0.32, *p* = .043), and the edge-wise GSR specifications small nonsignificant values (*d* = −0.12 and −0.11; Supplementary Table S8). Over all 480 specifications, run 2 gave a median *d* = −0.17 (joint permutation *p* = .019 for the median; *p* = .10 for both the number of significant negative specifications and their sum; Supplementary Table S3). Whether the effect habituated across runs therefore depends on the specification, so we make no claim of habituation.

The individualism analyses were exploratory, and the pre–post change in individualism turned out to be untestable. The eight individualism items had Cronbach’s *α* = .51 (pre) and .59 (post) and a pre–post correlation of .83; because that correlation exceeds both internal-consistency estimates, the classical difference-score reliability ^46^ estimated from these values is at or below zero for every subscale, and whatever change occurred cannot be separated from measurement error (Supplementary Table S9). For completeness, the change score correlated *r* = −0.06 with ΔEM. Among 12 trait predictors (total, horizontal and vertical individualism and collectivism, each at pre and post), none survived max-statistic permutation across the family. The largest correlations were with horizontal individualism (HI), the dimension our behavioral work had linked to pathogen cues^18^: *r* = 0.35 at pre and *r* = 0.35 at post (uncorrected *p* = .020 and .021; family-wise *p* = .13 and .14; Supplementary Table S10); higher trait HI was associated with a smaller reduction. Because HI was the dimension of prior interest within this exploratory family, we describe the association further, without treating it as evidence (Supplementary Table S11). The mean of the two HI scores correlated *r* = 0.36 with ΔEM (Spearman *ρ* = 0.30, *p* = .048). The partial correlations controlling for motion and for sex were 0.34 and 0.37, and the between-block component carried the association (*r* = 0.37; within-block *r* = 0.02). It vanished at a 2-TR shift (*r* = −0.18; within-block component *r* = −0.40), was absent in run 2 (*r* = −0.05), and reversed sign under the edge-wise GSR specification at 2 TR (*r* = −0.35; Supplementary Figure S3c–e). Leave-one-out correlations ranged from 0.19 to 0.42 and the bootstrap 95% interval was [−0.05, 0.65].

### The condition contrast does not replicate across runs

Any correlation between ΔEM and a trait presupposes that ΔEM is a stable property of a participant, and the two consecutive runs allow a direct check. Under the pre-specified analysis, ΔEM in run 1 correlated *r* = −0.32 (*p* = .037, *n* = 43) with ΔEM in run 2; under the edge-wise definition *r* = −0.43 (*p* = .004); under the GSR specifications *r* lay between −0.11 and +0.07 (Supplementary Figure S4a,b; Supplementary Table S12). The edge-metastability levels themselves were more stable across runs (*r* = 0.33 to 0.66 for the neutral and infection values without GSR and for the synchrony-based GSR values; 0.15 to 0.37 for the edge-wise GSR values), so the instability is specific to the condition contrast. The across-run covariance of ΔEM is a signed sum of four covariances between condition levels, two of which enter negatively, and the correlations between levels show where the negative sign comes from. Without GSR, run 1 infection levels correlated more strongly with run 2 neutral levels (*r* = 0.65) than with run 2 infection levels (*r* = 0.48), whereas run 1 neutral levels correlated similarly with both (0.46 and 0.43; Supplementary Table S20). Regression to the mean does not produce this pattern. It concerns change relative to an extreme baseline, and two noisy estimates of a contrast with no stable component would correlate near zero rather than negatively. A negative correlation means that participants with the largest reductions in run 1 tended to show increases in run 2. We have no established account of this reversal, which was confined to the specifications without GSR; it may reflect participant-specific habituation or sensitization, or an interaction of the fixed block order with within-run drift. In both cases the contrast fails to behave as a stable property of the participant. A state-level check with the in-scanner ratings gave the same answer. In block-level mixed models with condition and block order as covariates, the rating of a block, centered within participant and condition, did not predict that block’s metastability in either run (run 1: *b* = 0.010, *p* = .18 for the mean rating; run 2, *n* = 42 because one further participant lacked a run 2 rating log: *p* = .93; Supplementary Figure S4c, Supplementary Table S13), and the participant’s rating difference between conditions did not predict ΔEM (*r* = 0.16 and 0.02). Whether the behavioral measure is a trait or a state, then, the contrast is not reliable enough to support an individual-difference claim.

## Discussion

### The predicted reduction holds at the group level, but the design does not identify its cause

Infection-related images reduced cortical edge metastability. The pre-specified test alone gave a small effect at the significance threshold. The specification curve shows that its sign does not depend on analytic choices, since every significant specification points the same way and the joint permutation tests reject the null at *p < .*001, whereas its size depended on GSR and window shift. The control analyses then separate the forms of the effect. The synchrony-based reduction after GSR survived removal of the stimulus-locked response, detrending, and motion, white-matter and CSF regression, did not depend on how the global signal was fitted, was present at windows aligned with the block’s own hemodynamic response, and recurred in run 2, which played no part in selecting it. The GSR specifications were planned before the curve was computed, but neither they nor the nuisance-regressed form were the pre-specified test. The pre-specified value, without GSR and at onset, amounts to a reduction in the variability of global signal power. It vanished at hemodynamically aligned windows and after white-matter and CSF regression, which points to a global component shared with white matter and CSF rather than to cortical coupling. Without GSR the reduction is not a difference in static connectivity, and with GSR about half of it is not. In both cases the remainder is temporal: infection blocks contained fewer or smaller brain-wide bursts of cofluctuation.

The reduction has the sign predicted by the defensive-mode account, but so does a generic effect of stimulus salience, so the design confirms what the two accounts share without testing what separates them. Three readings remain open. The first motivated the study: a defensive mode in which the systems that produce disgust, attention and withdrawal are coordinated and coupling settles into a narrower repertoire. It fits the increased salience-network coupling that acute stress produces ^25^ and the withdrawal that pathogen cues are proposed to induce^4,5^, although lower metastability can also arise when regions fluctuate more independently. The second is attentional capture, which lowers metastability without any defensive withdrawal ^29^. The third is neuromodulation, a possible mechanism for either of the first two: higher neural gain tightens the clustering of functional connectivity ^47^, raising catecholamine levels reduces interregional correlations at rest ^48^, and global signal amplitude falls as vigilance rises ^31^, which predicts the pre-specified result directly. Separating these readings requires a test of specificity with a negatively valenced non-pathogen control. After nuisance regression with GSR the reduction was significant only in the visual, ventral attention and dorsal attention networks. The ventral attention network includes the anterior insula, which is active when disgust is experienced and observed ^19^, but this localization is post hoc, fits the defensive-mode and attentional readings equally well, and does not rule out a stimulus-driven change in visual cortex.

### The individual-difference predictions cannot be tested with this contrast

The sex difference was nominally present under the pre-specified analysis but did not survive correction. It was absent under the synchrony-based GSR specification and nominally reversed under the edge-wise GSR specification, as expected when specifications rank participants almost independently. Under the pre-specified analysis, individual differences in ΔEM tracked the static covariance of the windows (*r* = 0.74 with the surrogate contrast; Supplementary Table S16), so if the uncorrected sex difference is real it concerns the variability of global signal power rather than of coupling. The trait HI correlation did not survive correction and did not reappear in run 2. The across-run analysis shows why: ΔEM showed no positive correlation across runs above *r* = 0.09 under any of eight specifications, whereas the levels did. Difference scores from 19 or 20 blocks per condition have low reliability whatever the size of the group effect ^49^, as task-fMRI contrasts generally do ^50^, and the negative correlation without GSR also points to a systematic reversal between runs that we cannot explain. The change score has a different problem. Modest internal consistency and a high pre–post correlation give an estimated difference-score reliability at or below zero^46^, so the shift that our behavioral studies detected with 100 to 300 participants per condition^18^ cannot be resolved at the individual level here. None of this refutes the predictions^36,37,51^. They concern stable differences between people, and a contrast that does not replicate across runs cannot test them in either direction, whereas the group-level result requires only that the average participant shifts.

### What the measure contains

*R*(*t*) is a function of the parcel mean and the mean squared parcel amplitude alone, so the synchrony-based measure is a global-amplitude quantity with and without GSR. Without GSR it is the temporal variability of the power of the cortical parcel-mean signal; after GSR, which leaves a residual parcel mean whose power still dominates *R*(*t*), it is the variability of residual global power. The global signal carries respiratory, cardiac, arousal and motion contributions as well as neural coordination ^31,52,53^, and white-matter and CSF signals are regressed because they carry such non-neuronal fluctuations^44,45^. Removing them abolished the pre-specified effect and left the synchrony-based GSR effect in place, which makes the latter the better candidate for a change in cortical cofluctuation. Only the edge-wise definition depends on which pairs cofluctuate; its reduction survived the surrogate test but not detrending or white-matter and CSF regression. For Gaussian signals every definition used here follows from the static correlation structure^54,55^, and the surrogates place the condition difference outside that dependence, in the clustering of cofluctuation into bursts ^33^.

Because the GSR and no-GSR specifications rank participants almost independently, they measure different quantities, both lower during infection blocks at the group level, rather than noisy and clean versions of one. The regressed effect did not change when the global signal was fitted separately for each condition, which argues against the fit itself producing it, but the recentering of correlations that GSR introduces by construction^53^ remains and was the reason GSR was not the primary analysis. We therefore recommend reporting results with and without GSR as different measures. Published task effects on metastability have differed in direction across studies that differed in definition, task and sample ^29,30^. In principle the edge-wise measure is the right estimator for a task effect that changes the dispersion of cofluctuation across pairs without changing their mean, but here none of its effects survived white-matter and CSF regression, and its GSR variants were the least stable across runs and the most sensitive to motion and slow drift.

### Limitations

The design has three limitations that no analysis can remove. The block order was fixed and identical for all participants, so condition is confounded with position in the run; block-level metastability drifted upward, and on the strictly alternating segment the effects were smaller, the regressed effect among them, which there was not established at hemodynamically aligned windows. The control condition was neutral rather than negatively valenced, so the contrast cannot attribute the effect to pathogen relevance rather than to negative affect in general; in one of our two behavioral studies, non-pathogen threats produced many of the same shifts in social orientation^18^. With *n* = 44, the study had 80% power for *d* = 0.43 in a paired design at two-tailed *α* = .05 and about 50% for *d* = 0.30, which would need about 90 participants (Supplementary Table S15).

The measure brings further caveats. No respiratory or cardiac signal was recorded. The expression of disgust reduces nasal inspiration^13^, respiration is a principal source of global signal variance ^52^, and a change in breathing during infection blocks could produce the pre-specified result without a change in cortical coupling. The loss of that result after white-matter and CSF regression is consistent with such a source; the synchrony-based GSR effect survived that regression, but only physiological recording can exclude a respiratory contribution to it. Windows of 9 TRs are short for estimating a standard deviation, and without GSR the effect was present only when the window included the block transition. Detrending and nuisance regression were applied to the central specifications, not to the whole curve. Finally, run 2 was acquired in the same visit with the same images in the same order, so it replicates the measurement rather than the study.

### Outlook

For task-evoked metastability in block designs, we suggest fixing the primary specification before inspecting the data, reporting a specification curve over alignment, definition, normalization, parcellation, GSR and nuisance regression, removing the stimulus-locked response and nuisance signals before reading a contrast as a change in coupling, comparing the contrast with covariance-matched surrogates, and estimating its stability across runs before relating it to individual differences. A replication with randomized block order, a negatively valenced control, physiological recording, the regressed synchrony-based measure at hemodynamically aligned windows specified in advance, and *n ≥* 90 would address the questions left open here.

### Conclusion

Infection-related images reduced the moment-to-moment variability of global cortical cofluctuation in the predicted direction. One form of the reduction, the synchrony-based reduction after global signal regression, survived controls for task-evoked responses, head motion and white-matter and CSF signals, did not depend on how the global signal was fitted, recurred in a second run, and was strongest in visual and attention networks. It was smaller on strictly alternating blocks, and the pre-specified form, without global signal regression, did not survive removal of white-matter and CSF signals. The reduction is compatible with the defensive mode that the behavioral immune system is proposed to induce but does not demonstrate it, and as measured here it cannot serve as a marker of individual differences. This sequence, a pre-specified test followed by a specification curve, surrogates, controls for evoked and nuisance signals, and checks across runs, separated an effect that withstood the controls from variants that did not; we suggest it for any estimate of metastability from short task blocks.

## Methods

### Terms

Technical terms are defined after the references. The pre-specified analysis is the combination of analytic choices fixed before data inspection, named as such wherever it is applied (run 2, one network, one sex); the pre-specified test is its application to the whole cortex in run 1. Normalization within run is also called run-normalized. The two consecutive runs are called run 1 and run 2 here; the data release labels them 1 and 2 in its session column and S1 and S2 in file names.

### Participants

Forty-four healthy Japanese adults (21 women, 23 men; age *M* = 20.8 years, *SD* = 1.6, range 18– 24) were recruited from Tohoku University; the sample size was not determined by a power analysis. No participant was excluded from run 1. For one participant (participant 1 in the data release), the stored run 2 image file duplicated run 1 (identical image data and acquisition time), so the run 2 imaging data were unavailable; that run was excluded, and all run 2 and across-run analyses use *n* = 43. The study was approved by the MRI Research Ethics Committee of the Smart-Aging Research Center, Tohoku University (approval no. 2303-01, 21 March 2023), and was conducted in accordance with the Declaration of Helsinki. All participants gave written informed consent. Sex and age were taken from the participant registration record.

### Task and stimuli

Participants viewed alternating blocks of infection-related and neutral images (Figure 1a) presented with PsychoPy ^56^. The infection set (20 images) showed morphological and behavioral cues of communicable disease, which we selected from 120 candidates rated by 180 independent Japanese raters. Neutral images (20) came from the Disgust-Related Images (DIRTI) database^57^. Each run contained 40 blocks of 18 s in a fixed order, the same for all participants: three neutral blocks, three infection blocks, then strict alternation (Figure 1a). In each block one image was shown for 9 s, during which participants rated it for disgust, fear and perceived infection risk on 4-point scales (one prompt every 3 s, button-box response), followed by 9 s of fixation. Two runs with identical parameters were acquired consecutively within one scanning visit. Individualism and collectivism were measured with the 16-item horizontal–vertical individualism–collectivism scale^58^, translated into Japanese for this study, once before and once after the scanning visit; perceived vulnerability to disease was measured at the same times ^59^ and is not analyzed here.

### MRI acquisition and preprocessing

Imaging used a 3.0-T Philips Achieva dStream scanner. Blood-oxygen-level-dependent (BOLD) runs (repetition time (TR) 2,000 ms; 3.75 *×* 3.75 *×* 4 mm^3^; 360 volumes) were preprocessed with fMRIPrep 24.1.1^60^, including MCFLIRT motion correction ^61^, coregistration and normalization to MNI152NLin2009cAsym space at 2 mm. FD^62^, the brain-mask global signal and, for the control analyses, the motion parameters and the mean white-matter and CSF signals were taken from the fMRIPrep confound file. The first four volumes of each run were discarded. Apart from the optional GSR, no temporal filtering, detrending or nuisance regression was applied in the main analyses. Parcel time series were the mean over voxels of the Schaefer parcellations ^63^ at 100, 200, 400, 600, 800 and 1000 parcels in the 2-mm space of the fMRIPrep output (the 100- and 200-parcel versions were released after the original publication; all six are available from TemplateFlow^64^, https://www.templateflow.org, and from https://github.com/ThomasYeoLab/CBIG), and parcels were assigned to the seven Yeo networks ^65^ by the labels distributed with the atlas. When GSR was applied, the brain-mask global signal was regressed from each parcel time series within run, with one coefficient per parcel, before any further step in the main analyses.

### Block extraction

Block windows of 9 TRs (18 s) were placed at each block onset, shifted by 0, 1, 2 or 3 TR according to the specification. We fixed the onset-aligned window before data inspection because it covers the whole block as defined by the stimulus schedule (9 s of image and rating, 9 s of fixation). With a hemodynamic delay of about 4 to 6 s, its first two or three volumes still carry the response to the preceding block, and the end of the block’s own response falls into the next window; window shift was therefore one dimension of the specification curve. Because the first block began at the start of the run and its window overlapped the discarded volumes, it was excluded, leaving 20 infection and 19 neutral blocks per run. In the “common” block set the final block was also dropped so that all shifts used 19 blocks per condition; at shifts of one TR or more the final block falls outside the run and the two block sets coincide. The predicted hemodynamic response in Figure 1a is a canonical double-gamma function convolved with the 9-s image period.

### Edge metastability

Let *z_i_*(*t*) be the z-scored time series of parcel *i* (Figure 1b–e), normalized either within the run (before window extraction) or within the concatenated windows of a condition (after extraction). The global cofluctuation is

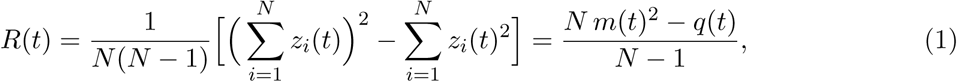

where *m*(*t*) is the mean of *z_i_*(*t*) over parcels and *q*(*t*) the mean of *z_i_*(*t*)^2^. *R*(*t*) is the mean over all parcel pairs of the edge time series *z_i_*(*t*)*z_j_*(*t*) and depends only on *m*(*t*) and *q*(*t*). The first term dominates when *N* Var*_t_*[*m*(*t*)] is large compared with Var*_t_*[*q*(*t*)], which held in these data with and without GSR (Supplementary Table S16), so *R*(*t*) *≈ m*(*t*)^2^. *R*(*t*) is the amplitude counterpart of mean pairwise phase coherence, and *m*(*t*)^2^ plays the role of the squared Kuramoto order parameter; edge metastability, the temporal standard deviation of *R*(*t*), is therefore the counterpart of the temporal variability of the squared order parameter, whereas the metastability index of models is the variability of the order parameter itself ^23^. The four definitions of edge metastability were: *synchrony-based* (the pre-specified definition), the standard deviation of *R*(*t*) over the concatenated windows of a condition; its *within-block* component, the mean over windows of the within-window standard deviation of *R*(*t*); its *between-block* component, the standard deviation over windows of the window means of *R*(*t*); and *edge-wise*, the mean over all parcel pairs of the temporal standard deviation of the edge time series *z_i_*(*t*)*z_j_*(*t*)^32^ over the concatenated windows. ΔEM is the infection minus neutral value for each participant. Network-level values were computed with the sums in the equation restricted to the parcels of one network. Parcel-wise edge-wise metastability was the mean over the 399 edges of a parcel of the temporal standard deviation of their cofluctuation; parcel-wise condition differences were tested with paired *t*-tests and BH-FDR across 400 parcels, as a descriptive map. Block-level metastability, used in the mixed models, is the standard deviation of *R*(*t*) within one 9-TR window after z-scoring within that window; it is related to, but not identical with, the within-block component, which uses z-scores from the concatenated windows.

### Statistical analysis

We wrote down the primary contrast before inspecting the data; the study was not preregistered on an external registry. The primary prediction was directional and the test was specified one-tailed; we report two-tailed *p* values throughout, the more conservative choice. We wrote the rest of the plan (secondary tests, specification curve, exploratory families and their corrections) after running the primary test and before computing the specification curve; it is summarized in the Supplementary Information and archived with the code. Later additions are the four extra parcellation resolutions, noted where they apply, and the composition, surrogate, control and replication analyses (Supplementary Tables S16 to S21), all computed after the specification curve. Permutation *p* values are the proportion of draws at least as extreme as the observed statistic (*b/B*; the (*b* + 1)*/*(*B* + 1) estimator differs by at most .0001 here), so *p < .*001 denotes fewer than 10 of 10,000 draws. All analyses used Python 3.10 with NumPy 2.2, SciPy 1.15, pandas 2.3 and statsmodels 0.15; every resampling procedure started from a NumPy generator seeded with 20260911, so a test applied to the same data returns the same *p* value wherever it is reported.

#### Primary test

Synchrony-based edge metastability, window at onset, normalization within condition, Schaefer 400, no GSR, all blocks, run 1; tested with a 10,000-draw sign-flip permutation on Cohen’s *d* of ΔEM (the mean of ΔEM divided by its standard deviation, *d_z_*; all within-participant effect sizes are of this form), with a 95% confidence interval from the noncentral *t* distribution. No correction is applied to the single primary test.

#### Secondary tests

A condition by sex difference in run 1 and a condition by run difference across runs, corrected with the Holm–Bonferroni procedure across the two (two-tailed, as for the primary test). With one ΔEM value per participant, the sex test is a Student’s *t* test of ΔEM between sexes (42 degrees of freedom); the participant-level mixed model specified in the plan is the same comparison without the loss of degrees of freedom (Results). The run test uses a mixed model over both runs (random intercepts, maximum likelihood, statsmodels MixedLM; 44 participants, 43 in run 2). Bayes factors for the interaction terms (BF_01_, evidence for the model without the term) were obtained from the Bayesian information criterion (BIC) difference between maximum-likelihood fits with and without the term, with the number of observations as *n* ^66^. Sex-stratified effects are descriptive.

#### Specification curve

Resting-state metastability is usually estimated from runs lasting many minutes ^23^; estimates of time-varying coupling depend on window length, shape and placement^35^ and on GSR, which itself changes correlation estimates^31,53^. Each choice is defensible and changes the estimate, so we report the whole space. The full crossing of window shift (0–3 TR), definition (four), normalization (two), parcellation (six), GSR (two) and block set (two), minus the block-set duplicates at non-zero shifts, gives 480 specifications. The plan as first written listed two parcellations (400 and 200 parcels; 160 specifications); we added the four further resolutions after computing the first curve, and the 160-specification subset gives the same conclusions (median *d* = −0.33, 89 significant). Each specification was tested with a two-tailed sign-flip permutation on *d*. For joint inference, adapted from Simonsohn et al. ^42^, the same 10,000 sign-flip draws were applied to all specifications to preserve their dependence, and the median *d*, the number of significant negative specifications and the sum of significant *d* values were compared with their null distributions; within draws, and for the observed count, significance was the parametric paired *t*-test at *p < .*05 (263 observed, as with the permutation criterion).

#### Static-connectivity surrogates

For jointly Gaussian signals, both measures are determined by the static correlation matrix of the concatenated windows. For each participant and condition we generated 30 Gaussian series with that covariance and no temporal structure (Schaefer 400, window at onset, normalization within condition, all blocks, run 1, with and without GSR), computed both measures, and tested the observed minus surrogate contrast with the sign-flip permutation (details in the Supplementary Information; Supplementary Table S16).

#### Control for evoked responses and nuisance signals

The four central specifications were the synchrony-based and edge-wise definitions with and without GSR (Schaefer 400, normalization within condition, all blocks, run 1). At shifts of 0 and 2 TR we applied six variants. Variant (i) subtracted, for each parcel, condition and time point within the window, the mean over that condition’s blocks before concatenation^43^. Variants (ii) to (iv) regressed linear and quadratic trends; 24 motion parameters, the six rigid-body parameters, their temporal derivatives and the squares of both ^45^; and the mean white-matter and CSF signals. Variant (v) applied (ii) to (iv) together, alone and followed by (i). Regressions were fitted within run on the post-dummy volumes; unlike the main analyses, the global signal entered the same model when GSR was applied. Nuisance regression, step (v) without (i), was also applied to network-level values at shift 0, and the somatomotor and visual networks were split by hemisphere. To test the form of the effect that survived these steps, all variants were repeated in run 2 (Figure 4), and the base and nuisance-regressed variants were repeated with the global signal fitted with separate coefficients for samples in infection windows, in neutral windows and elsewhere, and on the strictly alternating blocks 7 to 40; the contrast averaged over the two runs was tested in the 43 participants with both (Supplementary Table S21). Responses per block were counted from the in-scanner rating logs. These analyses re-estimate the same contrast under alternative preprocessing rather than test new hypotheses, so we report uncorrected *p* values and interpret the pattern across steps; for the synchrony-based GSR effect we also state whether it survives Holm–Bonferroni correction across the six variants. Network-level *q* values apply BH-FDR across the seven networks to paired *t*-test *p* values, as in the main analysis (Supplementary Table S14); all other *p* values, including the descriptive hemisphere comparisons, are from the sign-flip permutation.

#### Exploratory analyses

Sex-stratified effects, the within- and between-block decomposition, run 2, network-level effects (BH-FDR across seven networks), the amplitude checks (paired *t* tests), the strictly alternating segment (blocks 7 to 40) and the GSR contrasts without the 61 visual-network parcels are reported descriptively; their definitions are given in the Supplementary Information. The 12 trait predictors (total, horizontal and vertical individualism and collectivism at pre and post) were tested against ΔEM with Pearson correlation and corrected by max-statistic permutation over the family (20,000 draws). Sensitivity analyses for HI, the dimension of prior interest, are listed in Supplementary Table S11. Change-score reliability was computed with the classical difference-score formula ^46^, with the two internal consistencies standing in for the reliabilities of the pre and post scores. Stability of ΔEM and of the condition levels across the two runs was the Pearson correlation between runs (*n* = 43) under eight specifications (synchrony-based and edge-wise definitions, with and without GSR, shifts of 0 and 2 TR); for the onset-aligned specifications we decomposed the covariance of ΔEM across runs into the four covariances between condition levels. The in-scanner ratings, centered within participant and condition, entered block-level mixed models with condition and block order (Supplementary Information).

#### Motion

Block-wise FD entered the block-level mixed model as a covariate; the condition difference in FD was tested and correlated with ΔEM; effects were re-estimated after excluding the highest-FD quartile and, for GSR specifications, at zero FD difference by regression.

#### Power

With *n* = 44 the minimum detectable paired effect at 80% power and two-tailed *α* = .05 is *d_z_* = 0.43; *n* = 90 would be needed for *d_z_*= 0.30^67^.

## Data availability

Per-participant edge metastability under every specification, questionnaire item responses and scale scores, in-scanner ratings, motion summaries and all tables are available at https://github.com/choijeyoon/bis-edge-metastability (release v2.1.0; ^68^). The preprocessed imaging derivatives are available from the corresponding author under an institutional data-sharing agreement.

## Code availability

https://github.com/choijeyoon/bis-edge-metastability also holds the code for the parcellation, the specification curve, the decomposition, the surrogate and control analyses and all tables. The released scripts regenerate every table and figure from the per-participant data.

## Author contributions

Jeyoon Choi: Conceptualization; Data curation; Formal analysis; Investigation; Methodology; Software; Visualization; Writing – original draft; Writing – review & editing. Tomas Berjaga-Buisan: Formal analysis; Investigation; Methodology; Software; Validation; Visualization; Writing – review & editing. Elvira Garcia-Guzman: Formal analysis; Validation; Writing – review & editing. Motoaki Sugiura: Conceptualization; Funding acquisition; Resources; Supervision; Writing – review & editing. Gustavo Deco: Methodology; Supervision; Writing – review & editing.

## Use of generative AI

Language models (Anthropic Claude) were used to assist with editing and with drafting analysis code; all analyses were verified and all scientific content is the responsibility of the authors.

## Acknowledgments

J.C. was supported by the Japan Science and Technology Agency SPRING program (JPMJSP2114). M.S. was supported by the Mitsubishi Foundation. G.D. is supported by the project NEurological MEchanismS of Injury, and Sleep-like cellular dynamics (NEMESIS; ref. 101071900) funded by the EU ERC Synergy Horizon Europe. G.D. also acknowledges support from grant PID2022-136216NB-I00 funded by MICIU/AEI/10.13039/501100011033 and by “ERDF A way of making Europe”, ERDF, EU; and from the AGAUR research support grant (2021 SGR 00917) funded by the Department of Research and Universities of the Generalitat of Catalunya.

## Technical Terms

**Behavioral immune system:** Proposed psychological mechanisms that detect cues of infection and promote disgust and avoidance before contact.

**Edge time series:** Element-wise product of two z-scored regional time series; its value at each time point is their cofluctuation.

**Cofluctuation:** Product of two z-scored regional signals at one time point; its mean over time is their correlation.

**Global cofluctuation,** *R*(*t*): Mean of all edge time series at one time point; approximately the squared mean of the regional signals.

**Metastability:** Temporal variability of global synchrony, taken in models to index how widely dynamics move through possible states.

**Edge metastability:** Temporal standard deviation of global cofluctuation (synchrony-based) or mean temporal standard deviation of single edges (edge-wise).

**Global signal regression (GSR):** Removal of the whole-brain mean signal from every regional time series by linear regression.

**Nuisance regression:** Removal of slow trends and of signals attributed to head motion, white matter and cerebrospinal fluid.

**Specification:** One combination of analytic choices, such as window shift, definition, normalization, parcellation and global signal regression.

**Specification curve:** Effect estimates across all reasonable specifications, sorted by size, with joint inference across them.

**Sign-flip permutation:** Null distribution built by randomly flipping the sign of each participant’s condition difference.

**Covariance-matched surrogate:** Gaussian time series with the observed static covariance but no temporal structure.

## References

[1] Benjamin L. Hart. Biological basis of the behavior of sick animals. Neuroscience & Biobehavioral Reviews, 12(2):123–137, 1988. doi: 10.1016/s0149-7634(88)80004-6.

[2] Robert Dantzer, Jason C. O’Connor, Gregory G. Freund, Rodney W. Johnson, and Keith W. Kelley. From inflammation to sickness and depression: when the immune system subjugates the brain. Nature Reviews Neuroscience, 9(1):46–56, 2008. doi: 10.1038/nrn2297.

[3] Clifford B. Saper, Andrej A. Romanovsky, and Thomas E. Scammell. Neural circuitry engaged by prostaglandins during the sickness syndrome. Nature Neuroscience, 15(8):1088–1095, 2012. doi: 10.1038/nn.3159.

[4] Mark Schaller and Justin H. Park. The behavioral immune system (and why it matters). Current Directions in Psychological Science, 20(2):99–103, 2011. doi: 10.1177/0963721411402596.

[5] Mark Schaller. The behavioural immune system and the psychology of human sociality. Philosophical Transactions of the Royal Society B: Biological Sciences, 366(1583):3418–3426, 2011. doi: 10.1098/rstb.2011.0029.

[6] Damian R. Murray and Mark Schaller. The behavioral immune system: Implications for social cognition, social interaction, and social influence. In James M. Olson and Mark P. Zanna, editors, Advances in Experimental Social Psychology, volume 53, pages 75–129. Academic Press, 2016. doi: 10.1016/bs.aesp.2015.09.002.

[7] Joshua M. Ackerman, Sarah E. Hill, and Damian R. Murray. The behavioral immune system: Current concerns and future directions. Social and Personality Psychology Compass, 12(2): e12371, 2018. doi: 10.1111/spc3.12371.

[8] Joshua M. Ackerman, Joshua M. Tybur, and Aaron D. Blackwell. What role does pathogen-avoidance psychology play in pandemics? Trends in Cognitive Sciences, 25(3):177–186, 2021. doi: 10.1016/j.tics.2020.11.008.

[9] Megan Oaten, Richard J. Stevenson, and Trevor I. Case. Disgust as a disease-avoidance mechanism. Psychological Bulletin, 135(2):303–321, 2009. doi: 10.1037/a0014823.

[10] Joshua M. Tybur, Debra Lieberman, and Vladas Griskevicius. Microbes, mating, and morality: Individual differences in three functional domains of disgust. Journal of Personality and Social Psychology, 97(1):103–122, 2009. doi: 10.1037/a0015474.

[11] Valerie Curtis, Michéal de Barra, and Robert Aunger. Disgust as an adaptive system for disease avoidance behaviour. Philosophical Transactions of the Royal Society B, 366(1563):389–401, 2011. doi: 10.1098/rstb.2010.0117.

[12] Joshua M. Tybur, Debra Lieberman, Robert Kurzban, and Peter DeScioli. Disgust: Evolved function and structure. Psychological Review, 120(1):65–84, 2013. doi: 10.1037/a0030778.

[13] Joshua M Susskind, Daniel H Lee, Andrée Cusi, Roman Feiman, Wojtek Grabski, and Adam K Anderson. Expressing fear enhances sensory acquisition. Nature Neuroscience, 11(7):843–850, 2008. doi: 10.1038/nn.2138.

[14] Joshua M. Ackerman, D. Vaughn Becker, Chad R. Mortensen, Takao Sasaki, Steven L. Neuberg, and Douglas T. Kenrick. A pox on the mind: Disjunction of attention and memory in the processing of physical disfigurement. Journal of Experimental Social Psychology, 45(3):478– 485, 2009. doi: 10.1016/j.jesp.2008.12.008.

[15] Chad R. Mortensen, D. Vaughn Becker, Joshua M. Ackerman, Steven L. Neuberg, and Douglas T. Kenrick. Infection breeds reticence: The effects of disease salience on self-perceptions of personality and behavioral avoidance tendencies. Psychological Science, 21(3):440–447, 2010. doi: 10.1177/0956797610361706.

[16] Corey L. Fincher, Randy Thornhill, Damian R. Murray, and Mark Schaller. Pathogen prevalence predicts human cross-cultural variability in individualism/collectivism. Proceedings of the Royal Society B, 275(1640):1279–1285, 2008. doi: 10.1098/rspb.2008.0094.

[17] Randy Thornhill, Corey L. Fincher, and Devaraj Aran. Parasites, democratization, and the liberalization of values across contemporary countries. Biological Reviews, 84(1):113–131, 2009. doi: 10.1111/j.1469-185X.2008.00062.x.

[18] Jeyoon Choi and Motoaki Sugiura. Infection changes social orientations: The effect of disease salience on shifts in individualism and collectivism. Personality and Individual Differences, 250:113524, 2026. doi: 10.1016/j.paid.2025.113524.

[19] Bruno Wicker, Christian Keysers, Jane Plailly, Jean-Pierre Royet, Vittorio Gallese, and Giacomo Rizzolatti. Both of us disgusted in my insula: The common neural basis of seeing and feeling disgust. Neuron, 40(3):655–664, 2003. doi: 10.1016/S0896-6273(03)00679-2.

[20] Jana Schaich Borg, Debra Lieberman, and Kent A. Kiehl. Infection, incest, and iniquity: Investigating the neural correlates of disgust and morality. Journal of Cognitive Neuroscience, 20(9):1529–1546, 2008. doi: 10.1162/jocn.2008.20109.

[21] Christina Regenbogen, John Axelsson, Julie Lasselin, Danja K. Porada, Tina Sundelin, Moa G. Peter, et al. Behavioral and neural correlates to multisensory detection of sick humans. Proceedings of the National Academy of Sciences, 114(24):6400–6405, 2017. doi: 10.1073/pnas.1617357114.

[22] Emmanuelle Tognoli and J. A. Scott Kelso. The metastable brain. Neuron, 81(1):35–48, 2014. doi: 10.1016/j.neuron.2013.12.022.

[23] Gustavo Deco, Morten L. Kringelbach, Viktor K. Jirsa, and Petra Ritter. The dynamics of resting fluctuations in the brain: metastability and its dynamical cortical core. Scientific Reports, 7:3095, 2017. doi: 10.1038/s41598-017-03073-5.

[24] P. J. Hellyer, G. Scott, M. Shanahan, D. J. Sharp, and R. Leech. Cognitive flexibility through metastable neural dynamics is disrupted by damage to the structural connectome. Journal of Neuroscience, 35(24):9050–9063, 2015. doi: 10.1523/JNEUROSCI.4648-14.2015.

[25] Erno J. Hermans, Hein J. F. van Marle, Lindsey Ossewaarde, Marloes J. A. G. Henckens, Shaozheng Qin, Marlieke T. R. van Kesteren, et al. Stress-related noradrenergic activity prompts large-scale neural network reconfiguration. Science, 334(6059):1151–1153, 2011. doi: 10.1126/science.1209603.

[26] Michael S. Fanselow. Neural organization of the defensive behavior system responsible for fear. Psychonomic Bulletin & Review, 1(4):429–438, 1994. doi: 10.3758/BF03210947.

[27] Dean Mobbs, Cindy C. Hagan, Tim Dalgleish, Brian Silston, and Charlotte Prévost. The ecology of human fear: Survival optimization and the nervous system. Frontiers in Neuroscience, 9:55, 2015. doi: 10.3389/fnins.2015.00055.

[28] Karin Roelofs. Freeze for action: neurobiological mechanisms in animal and human freezing. Philosophical Transactions of the Royal Society B: Biological Sciences, 372(1718):20160206, 2017. doi: 10.1098/rstb.2016.0206.

[29] Peter J. Hellyer, Murray Shanahan, Gregory Scott, Richard J. S. Wise, David J. Sharp, and Robert Leech. The control of global brain dynamics: Opposing actions of frontoparietal control and default mode networks on attention. The Journal of Neuroscience, 34(2):451–461, 2014. doi: 10.1523/JNEUROSCI.1853-13.2014.

[30] Thomas H. Alderson, Arun L. W. Bokde, J. A. Scott Kelso, Liam Maguire, and Damien Coyle. Metastable neural dynamics underlies cognitive performance across multiple behavioural paradigms. Human Brain Mapping, 41(12):3212–3234, 2020. doi: 10.1002/hbm.25009.

[31] Thomas T Liu, Alican Nalci, and Maryam Falahpour. The global signal in fMRI: Nuisance or information? NeuroImage, 150:213–229, 2017. doi: 10.1016/j.neuroimage.2017.02.036.

[32] Joshua Faskowitz, Farnaz Zamani Esfahlani, Youngheun Jo, Olaf Sporns, and Richard F. Betzel. Edge-centric functional network representations of human cerebral cortex reveal overlapping system-level architecture. Nature Neuroscience, 23(12):1644–1654, 2020. doi: 10.1038/s41593-020-00719-y.

[33] Farnaz Zamani Esfahlani, Youngheun Jo, Joshua Faskowitz, Lisa Byrge, Daniel P. Kennedy, Olaf Sporns, et al. High-amplitude cofluctuations in cortical activity drive functional connectivity. Proceedings of the National Academy of Sciences, 117(45):28393–28401, 2020. doi: 10.1073/pnas.2005531117.

[34] Richard F. Betzel, Joshua Faskowitz, and Olaf Sporns. Living on the edge: network neuroscience beyond nodes. Trends in Cognitive Sciences, 27(11):1068–1084, 2023. doi: 10.1016/j.tics.2023.08.009.

[35] Daniel J. Lurie, Daniel Kessler, Danielle S. Bassett, Richard F. Betzel, Michael Breakspear, Shella Keilholz, et al. Questions and controversies in the study of time-varying functional connectivity in resting fmri. Network Neuroscience, 4(1):30–69, 2020. doi: 10.1162/netn_a_00116.

[36] Laith Al-Shawaf, David M. G. Lewis, and David M. Buss. Sex differences in disgust: Why are women more easily disgusted than men? Emotion Review, 10(2):149–160, 2018. doi: 10.1177/1754073917709940.

[37] Xavier Caseras, David Mataix-Cols, Suk Kyoon An, Natalia S. Lawrence, Anne Speckens, Vincent Giampietro, et al. Sex differences in neural responses to disgusting visual stimuli: Implications for disgust-related psychiatric disorders. Biological Psychiatry, 62(5):464–471, 2007. doi: 10.1016/j.biopsych.2006.10.030.

[38] Catharine H. Rankin, Thomas Abrams, Robert J. Barry, Seema Bhatnagar, David F. Clayton, John Colombo, et al. Habituation revisited: An updated and revised description of the behavioral characteristics of habituation. Neurobiology of Learning and Memory, 92(2):135–138, 2009. doi: 10.1016/j.nlm.2008.09.012.

[39] Joshua Carp. On the plurality of (methodological) worlds: estimating the analytic flexibility of fMRI experiments. Frontiers in Neuroscience, 6:149, 2012. doi: 10.3389/fnins.2012.00149.

[40] Rotem Botvinik-Nezer, Felix Holzmeister, Colin F Camerer, Anna Dreber, Juergen Huber, Magnus Johannesson, et al. Variability in the analysis of a single neuroimaging dataset by many teams. Nature, 582(7810):84–88, 2020. doi: 10.1038/s41586-020-2314-9.

[41] Sara Steegen, Francis Tuerlinckx, Andrew Gelman, and Wolf Vanpaemel. Increasing transparency through a multiverse analysis. Perspectives on Psychological Science, 11(5):702–712, 2016. doi: 10.1177/1745691616658637.

[42] Uri Simonsohn, Joseph P Simmons, and Leif D Nelson. Specification curve analysis. Nature Human Behaviour, 4(11):1208–1214, 2020. doi: 10.1038/s41562-020-0912-z.

[43] Michael W. Cole, Takuya Ito, Douglas Schultz, Ravi Mill, Richard Chen, and Carrisa Cocuzza. Task activations produce spurious but systematic inflation of task functional connectivity estimates. NeuroImage, 189:1–18, 2019. doi: 10.1016/j.neuroimage.2018.12.054.

[44] Rastko Ciric, Daniel H. Wolf, Jonathan D. Power, David R. Roalf, Graham L. Baum, Kosha Ruparel, et al. Benchmarking of participant-level confound regression strategies for the control of motion artifact in studies of functional connectivity. NeuroImage, 154:174–187, 2017. doi: 10.1016/j.neuroimage.2017.03.020.

[45] Theodore D. Satterthwaite, Mark A. Elliott, Raphael T. Gerraty, Kosha Ruparel, James Loughead, Monica E. Calkins, et al. An improved framework for confound regression and filtering for control of motion artifact in the preprocessing of resting-state functional connectivity data. NeuroImage, 64:240–256, 2013. doi: 10.1016/j.neuroimage.2012.08.052.

[46] Lee J Cronbach and Lita Furby. How we should measure “change”: Or should we? Psychological Bulletin, 74(1):68–80, 1970. doi: 10.1037/h0029382.

[47] Eran Eldar, Jonathan D. Cohen, and Yael Niv. The effects of neural gain on attention and learning. Nature Neuroscience, 16(8):1146–1153, 2013. doi: 10.1038/nn.3428.

[48] Ruud L. van den Brink, Thomas Pfeffer, Christopher M. Warren, Peter R. Murphy, Klodiana-Daphne Tona, Nic J. A. van der Wee, et al. Catecholaminergic neuromodulation shapes intrinsic MRI functional connectivity in the human brain. The Journal of Neuroscience, 36(30):7865–7876, 2016. doi: 10.1523/JNEUROSCI.0744-16.2016.

[49] Craig Hedge, Georgina Powell, and Petroc Sumner. The reliability paradox: Why robust cognitive tasks do not produce reliable individual differences. Behavior Research Methods, 50 (3):1166–1186, 2018. doi: 10.3758/s13428-017-0935-1.

[50] Maxwell L. Elliott, Annchen R. Knodt, David Ireland, Meriwether L. Morris, Richie Poulton, Sandhya Ramrakha, et al. What is the test-retest reliability of common task-functional MRI measures? New empirical evidence and a meta-analysis. Psychological Science, 31(7):792–806, 2020. doi: 10.1177/0956797620916786.

[51] Diana S. Fleischman and Daniel M. T. Fessler. Progesterone’s effects on the psychology of disease avoidance: Support for the compensatory behavioral prophylaxis hypothesis. Hormones and Behavior, 59(2):271–275, 2011. doi: 10.1016/j.yhbeh.2010.11.014.

[52] Jonathan D Power, Mark Plitt, Timothy O Laumann, and Alex Martin. Sources and implications of whole-brain fMRI signals in humans. NeuroImage, 146:609–625, 2017. doi: 10.1016/j.neuroimage.2016.09.038.

[53] Kevin Murphy and Michael D Fox. Towards a consensus regarding global signal regression for resting state functional connectivity MRI. NeuroImage, 154:169–173, 2017. doi: 10.1016/j.neuroimage.2016.11.052.

[54] Leonardo Novelli and Adeel Razi. A mathematical perspective on edge-centric brain functional connectivity. Nature Communications, 13(1):2693, 2022. doi: 10.1038/s41467-022-29775-7.

[55] Zach Ladwig, Benjamin A. Seitzman, Ally Dworetsky, Yuhua Yu, Babatunde Adeyemo, Derek M. Smith, et al. BOLD cofluctuation ‘events’ are predicted from static functional connectivity. NeuroImage, 260:119476, 2022. doi: 10.1016/j.neuroimage.2022.119476.

[56] Jonathan Peirce, Jeremy R. Gray, Sol Simpson, Michael MacAskill, Richard Höchenberger, Hiroyuki Sogo, et al. PsychoPy2: Experiments in behavior made easy. Behavior Research Methods, 51(1):195–203, 2019. doi: 10.3758/s13428-018-01193-y.

[57] Anke Haberkamp, Julia A. Glombiewski, Frank Schmidt, and Antonia Barke. The DIsgust-RelaTed-Images (DIRTI) database: Validation of a novel standardized set of disgust pictures. Behaviour Research and Therapy, 89:86–94, 2017. doi: 10.1016/j.brat.2016.11.010.

[58] Harry C Triandis and Michele J Gelfand. Converging measurement of horizontal and vertical individualism and collectivism. Journal of Personality and Social Psychology, 74(1):118–128, 1998. doi: 10.1037/0022-3514.74.1.118.

[59] Lesley A Duncan, Mark Schaller, and Justin H Park. Perceived vulnerability to disease: Development and validation of a 15-item self-report instrument. Personality and Individual Differences, 47(6):541–546, 2009. doi: 10.1016/j.paid.2009.05.001.

[60] Oscar Esteban, Christopher J. Markiewicz, Ross W. Blair, Craig A. Moodie, A. Ilkay Isik, Asier Erramuzpe, et al. fMRIPrep: a robust preprocessing pipeline for functional MRI. Nature Methods, 16(1):111–116, 2019. doi: 10.1038/s41592-018-0235-4.

[61] Mark Jenkinson, Peter Bannister, Michael Brady, and Stephen Smith. Improved optimization for the robust and accurate linear registration and motion correction of brain images. NeuroImage, 17(2):825–841, 2002. doi: 10.1006/nimg.2002.1132.

[62] Jonathan D. Power, Anish Mitra, Timothy O. Laumann, Abraham Z. Snyder, Bradley L. Schlaggar, and Steven E. Petersen. Methods to detect, characterize, and remove motion artifact in resting state fmri. NeuroImage, 84:320–341, 2014. doi: 10.1016/j.neuroimage.2013.08.048.

[63] Alexander Schaefer, Ru Kong, Evan M. Gordon, Timothy O. Laumann, Xi-Nian Zuo, Avram J. Holmes, et al. Local-global parcellation of the human cerebral cortex from intrinsic functional connectivity mri. Cerebral Cortex, 28(9):3095–3114, 2018. doi: 10.1093/cercor/bhx179.

[64] Rastko Ciric, William H. Thompson, Romy Lorenz, Mathias Goncalves, Eilidh E. MacNicol, Christopher J. Markiewicz, et al. TemplateFlow: FAIR-sharing of multi-scale, multi-species brain models. Nature Methods, 19(12):1568–1571, 2022. doi: 10.1038/s41592-022-01681-2.

[65] B. T. Thomas Yeo, Fenna M. Krienen, Jorge Sepulcre, Mert R. Sabuncu, Danial Lashkari, Marisa Hollinshead, et al. The organization of the human cerebral cortex estimated by intrinsic functional connectivity. Journal of Neurophysiology, 106(3):1125–1165, 2011. doi: 10.1152/jn.00338.2011.

[66] Eric-Jan Wagenmakers. A practical solution to the pervasive problems of p values. Psychonomic Bulletin & Review, 14(5):779–804, 2007. doi: 10.3758/BF03194105.

[67] Franz Faul, Edgar Erdfelder, Albert-Georg Lang, and Axel Buchner. G*Power 3: A flexible statistical power analysis program for the social, behavioral, and biomedical sciences. Behavior Research Methods, 39(2):175–191, 2007. doi: 10.3758/BF03193146.

[68] Jeyoon Choi, Tomas Berjaga-Buisan, Elvira Garcia-Guzman, Motoaki Sugiura, and Gustavo Deco. Code and derived data for “Infection-related images reduce cortical edge metastability” (v2.1.0). GitHub repository, release v2.1.0, 2026. URL https://github.com/choijeyoon/bis-edge-metastability/releases/tag/v2.1.0.

