## Supplementary Information for "Infection-related images reduce cortical edge metastability: a specification-curve analysis of a block-design fMRI study"

[1]Institute of Development, Aging and Cancer, Tohoku University, Sendai, Japan. [2]Center for Brain and Cognition, Computational Neuroscience Group, Department of Engineering, Universitat Pompeu Fabra, Barcelona, Spain. [3]Vall d'Hebron Institut de Recerca (VHIR), Barcelona, Spain. [4]Center for Brain and Cognition, Computational Neuroscience Group, Faculty of Medicine and Life Sciences, Universitat Pompeu Fabra, 08005 Barcelona, Spain. [5]Institució Catalana de Recerca i Estudis Avançats (ICREA), 08010 Barcelona, Spain. [6]International Centre for Flourishing, Universities of Oxford (UK), Aarhus (Denmark) and Pompeu Fabra (Spain), Oxford OX3 9BX, UK. [7]Computational and Theoretical Neuroscience Lab, Institut de Recerca Sant Joan de Déu (IRSJD), Santa Rosa 39–57, 08950 Esplugues de Llobregat, Spain.

### S1 Supplementary Methods

#### S1.1 Stimulus selection and validation

Stimuli comprised images from two categories: infection-related threats and neutral scenes. A pool of 120 candidate infection-related pictures was initially compiled from publicly available online sources and evaluated by an independent sample of 180 Japanese raters. Each image was rated on three dimensions: disgust, fear, and perceived infection risk. Ratings were averaged across raters for each picture. To identify stimuli specifically associated with disease threat, as distinct from a broader negative-valence response, we computed a composite score by averaging the disgust and perceived infection risk ratings and subtracting the fear rating. This score favors items with a disgust / contamination profile over items with a primarily fear-based profile. The 20 highest-scoring images were retained for use in the fMRI experiment. Retained pictures depicted morphological or behavioral cues linked to infectious diseases, such as coughing, skin lesions, and disease-transmitting insects.

For the neutral condition, we selected 20 images from the Disgust-Related-Images (DIRTI) database[1], which provides affectively validated stimuli spanning a range of disgust levels. We specifically chose items rated low in disgust. Low-level visual properties of the two sets were not matched (Supplementary Table S1).

Low-level visual features of the two stimulus sets (luminance, contrast, spatial frequency content, edge density, hue, and entropy) were quantified post-hoc; results are reported in Section S2.1.

#### S1.2 Behavioral measures

Social orientation was measured with the 16-item horizontal–vertical individualism–collectivism scale of Triandis and Gelfand[2], translated into Japanese for this study, with four items each for horizontal individualism (HI), vertical individualism (VI), horizontal collectivism (HC) and vertical collectivism (VC). In this study items were rated from 0 to 6. Totals for individualism (IND, 8 items) and collectivism (COL, 8 items) and the four subscale sums were computed from the item-level Google Forms exports by participant identifier. No item is reverse-keyed. The scale was administered once before and once after the scanning visit. Internal consistencies and pre–post correlations are given in Table S9. Perceived vulnerability to disease[3] was administered at the same two times and is not analyzed in this manuscript.

#### S1.3 Procedure details

Upon arrival, participants received instructions about the experimental procedures and completed the individualism–collectivism and perceived-vulnerability questionnaires. Sex and age were taken from the participant registration record. During scanning, participants performed a visual rating task presented via PsychoPy[4], responding with an MRI-compatible response box. Each participant completed two consecutive functional runs within a single scanning visit, each lasting 720 s (360 volumes, TR = 2 s). These are referred to as run 1 and run 2; the data release labels them 1 and 2 in its session column and S1 and S2 in file names. Both runs used identical stimuli and task parameters, with run 2 serving as a repeated administration to assess whether the effects observed in run 1 persisted upon immediate re-exposure. For participant 1, the stored run 2 image file duplicated run 1 (identical image data and acquisition time), so the run 2 imaging data were unavailable. That run was excluded, and all run 2 and across-run analyses use $n = 43$.

Each block lasted 18 s and was composed of a 9 s stimulus presentation period followed by a 9 s fixation cross. During the stimulus presentation period one image from the condition of the block was shown for 9 s, and participants rated how strongly it evoked (1) disgust, (2) fear, and (3) perceived infection risk on a 4-point Likert scale (1 = "not related at all," 4 = "very highly related"). The rating prompts appeared one at a time below the image, 3 s each, and participants responded by pressing the corresponding button on the response box. The image remained on screen for the full 9 s whether or not a response had been given. The experimental sequence began with three initial neutral blocks, followed by three infection-related blocks, and then alternating neutral and infection-related blocks. Participants were instructed to maintain fixation on the central cross, attend to the images, and provide ratings as accurately as possible. After MRI scanning, participants again completed the individualism–collectivism questionnaire.

#### S1.4 MRI acquisition parameters

Neuroimaging data were collected on a 3.0 T Philips Achieva dStream scanner (Philips Healthcare, Best, The Netherlands) at Tohoku University. High-resolution structural images were acquired using a T1-weighted 3D MPRAGE sequence with the following parameters: TR $\approx$ 11.3 ms; TE $\approx$ 5.2 ms; flip angle = 8°; slice thickness = 0.7 mm; no gap; number of slices = 368; FoV = 256 mm; acquisition matrix = 368 × 368; voxel resolution = 0.7 × 0.7 × 0.7 mm$^3$. SENSE parallel imaging was applied in both in-plane and through-plane directions. These data were used for anatomical reference and spatial normalization.

Functional images were acquired using a T2*-weighted echo-planar imaging (EPI) sequence with the following parameters: TR = 2,000 ms; TE = 30 ms; flip angle = 80°; slice thickness = 4 mm, no gap; number of slices = 32; FoV = 240 mm; acquisition matrix = 64 × 64; voxel resolution = 3.75 × 3.75 × 4 mm$^3$; and ascending slice order. SENSE parallel imaging was applied. Each of the two consecutive functional runs (runs 1 and 2) consisted of 360 volumes, corresponding to a duration of 720 s.

#### S1.5 fMRI preprocessing

The following description was generated by fMRIPrep and is reproduced unchanged except for one [*sic*], as its license intends.

Results included in this manuscript come from preprocessing performed using *fMRIPrep* 24.1.1 (Esteban et al.[5]; Esteban et al.[6]; RRID:SCR_016216), which is based on *Nipype* 1.8.6 (Gorgolewski et al.[7]; Gorgolewski et al.[8]; RRID:SCR_002502).

**Anatomical data preprocessing** A total of 1 T1-weighted (T1w) images were found within the input BIDS dataset. The T1w image was corrected for intensity non-uniformity (INU) with `N4BiasFieldCorrection`[9], distributed with ANTs 2.5.3[10] RRID:SCR_004757, and used as T1w-reference throughout the workflow. The T1w-reference was then skull-stripped with a *Nipype* implementation of the `antsBrainExtraction.sh` workflow (from ANTs), using OASIS30ANTs as target template. Brain tissue segmentation of cerebrospinal fluid (CSF), white-matter (WM) and gray-matter (GM) was performed on the brain-extracted T1w using `fast` FSL (version unknown), RRID:SCR_002823,[11]. Volume-based spatial normalization to one standard space (MNI152NLin2009cAsym) was performed through nonlinear registration with `antsRegistration` (ANTs 2.5.3), using brain-extracted versions of both T1w reference and the T1w template. The following template was were [*sic*] selected for spatial normalization and accessed with *TemplateFlow* 24.2.0,[12]: *ICBM 152 Nonlinear Asymmetrical template version 2009c* [Fonov et al.[13], RRID:SCR_008796; TemplateFlow ID: MNI152NLin2009cAsym].

**Functional data preprocessing** For each of the 2 BOLD runs found per subject (across all tasks and sessions), the following preprocessing was performed. First, a reference volume was generated, using a custom methodology of *fMRIPrep*, for use in head motion correction. Head-motion parameters with respect to the BOLD reference (transformation matrices, and six corresponding rotation and translation parameters) are estimated before any spatiotemporal filtering using `mcflirt` FSL ,[14]. The BOLD reference was then co-registered to the T1w reference using `mri_coreg` (FreeSurfer) followed by `flirt` FSL ,[15] with the boundary-based registration[16] cost-function. Co-registration was configured with six degrees of freedom. Several confounding time-series were calculated based on the *preprocessed BOLD*: framewise displacement (FD), DVARS and three region-wise global signals. FD was computed using two formulations following Power (absolute sum of relative motions, Power et al.[17]) and Jenkinson (relative root mean square displacement between affines, Jenkinson et al.[14]). FD and DVARS are calculated for each functional run, both using their implementations in *Nipype* following the definitions by[17]. The three global signals are extracted within the CSF, the WM, and the whole-brain masks. Additionally, a set of physiological regressors were extracted to allow for component-based noise correction *CompCor*,[18]. Principal components are estimated after high-pass filtering the *preprocessed BOLD* time-series (using a discrete cosine filter with 128s cut-off) for the two *CompCor* variants: temporal (tCompCor) and anatomical (aCompCor). tCompCor components are then calculated from the top 2% variable voxels within the brain mask. For aCompCor, three probabilistic masks (CSF, WM and combined CSF+WM) are generated in anatomical space. The implementation differs from that of Behzadi et al. in that instead of eroding the masks by 2 pixels on BOLD space, a mask of pixels that likely contain a volume fraction of GM is subtracted from the aCompCor masks. This mask is obtained by thresholding the corresponding partial volume map at 0.05, and it ensures components are not extracted from voxels containing a minimal fraction of GM. Finally, these masks are resampled into BOLD space and binarized by thresholding at 0.99 (as in the original implementation). Components are also calculated separately within the WM and CSF masks. For each CompCor decomposition, the $k$ components with the largest singular values are retained, such that the retained components' time series are sufficient to explain 50 percent of variance across the nuisance mask (CSF, WM, combined, or temporal). The remaining components are dropped from consideration. The head-motion estimates calculated in the correction step were also placed within the corresponding confounds file. The confound time series derived from head motion estimates and global signals were expanded with the inclusion of temporal derivatives and quadratic terms for each[19]. Frames that exceeded a threshold of 0.5 mm FD or 1.5 standardized DVARS were annotated as motion outliers. Additional nuisance timeseries are calculated by means of principal components analysis of the signal found within a thin band (*crown*) of voxels around the edge of the brain, as proposed by[20]. All resamplings can be performed with *a single interpolation step* by composing all the pertinent transformations (i.e. head-motion transform matrices, susceptibility distortion correction when available, and co-registrations to anatomical and output spaces). Gridded (volumetric) resamplings were performed using `nitransforms`, configured with cubic B-spline interpolation.

Many internal operations of *fMRIPrep* use *Nilearn* 0.10.4[21] RRID:SCR_001362, mostly within the functional processing workflow. For more details of the pipeline, see the section corresponding to workflows in *fMRIPrep*'s documentation.

The first four volumes of each run were discarded before any analysis. No participant was excluded for head motion. Global signal regression was not part of preprocessing; where a specification applies it, the fMRIPrep whole-brain global signal is regressed from the parcel time series at the analysis stage (Edge metastability definitions below).

#### S1.6 Parcellation and network assignment

Parcel time series were extracted in the space of the fMRIPrep output (MNI152NLin2009cAsym, 2-mm isotropic, $97 \times 115 \times 97$ voxels) with the Schaefer atlases[22] at 100, 200, 400, 600, 800 and 1000 parcels (the 100- and 200-parcel versions were released by the authors after the original publication), obtained from TemplateFlow in that space[12]; each parcel value at each volume is the mean over the voxels carrying that label. The 400-parcel atlas is the primary parcellation; the others enter the specification curve, and the 200-parcel atlas also enters the motion and individual-difference checks (Tables S4 and S8). Each parcel was assigned to one of the seven resting-state networks of Yeo and colleagues[23] (visual, somatomotor, dorsal attention, ventral attention, limbic, control, default mode) by the labels distributed with the atlas.

#### S1.7 Block-wise time series extraction

Block onsets were read from the PsychoPy event files. For each block a window of nine TRs (18 s) was extracted starting at the onset shifted by 0, 1, 2 or 3 TR according to the specification. The first block of each run began at 0 s and its window overlapped the four discarded volumes, so it was excluded, leaving 20 infection and 19 neutral blocks. In the common block set the final block was also excluded so that every shift used 19 blocks per condition; at shifts of one TR or more the final block falls outside the run and the two block sets coincide. Normalization was applied either to the whole post-dummy run before extraction ("within run") or to the concatenated blocks of each condition after extraction ("within condition"). The primary analysis uses normalization within condition.

##### S1.8 Edge metastability: implementation details

The definitions are given in the main Methods. Writing $R_b(t)$ for the values of $R(t)$ within window $b$, the within-block component is the mean over windows of $\mathrm{SD}_t[R_b(t)]$ and the between-block component is the standard deviation over windows of the window means $\overline{R_b}$. By the law of total variance, the squared synchrony-based value equals the mean within-window variance of $R(t)$ plus the variance of the window means, up to degrees-of-freedom factors. For the within-block component the identity is only approximate, because that component is the mean of the within-window standard deviations, whereas the identity requires the square root of the mean within-window variance. The edge-wise definition averages $\mathrm{SD}_t[z_i(t)z_j(t)]$ over all $N(N-1)/2$ pairs. The global signal, when regressed, was the fMRIPrep brain-mask global signal, removed from each parcel time series within run by least squares before normalization.

##### S1.9 Statistical analysis plan

The analysis plan (primary test, secondary tests, specification-curve dimensions, exploratory families and their corrections) was written after the primary test had been run and before the specification curve was computed. It is summarized here and archived with the code. The primary contrast had been fixed before data inspection. Joint inference over the specification curve was adapted from Simonsohn et al.[24]: 10,000 sign-flip draws were applied to the participant-level ΔEM values identically across all specifications, and the observed median $d$ and number of significant negative specifications (two of the statistics they proposed), together with the sum of significant $d$ (an additional statistic), were compared with the null distributions so generated. Within each draw, and for the observed count, a specification counted as significant when its parametric paired $t$ test gave $p < .05$. The plan as first written listed the 400- and 200-parcel atlases. Later additions were the four extra parcellation resolutions (100, 600, 800 and 1000 parcels), added after the first curve had been computed and marked as additions in the archived plan, and the composition, surrogate, control and replication analyses (Tables S16 to S21), all computed after the specification curve. Individual specifications were tested with two-tailed sign-flip permutation on $d$. The trait family used max-statistic permutation: for each of 20,000 permutations of ΔEM the maximum absolute correlation across the 12 predictors was recorded, and the family-wise p of each observed correlation is the proportion of maxima at or above it. For HI, the dimension of prior interest, Table S11 gives the Spearman correlation, a bootstrap interval (10,000 resamples), the range of leave-one-out correlations, partial correlations controlling for mean FD and for sex, and the values under other specifications. The in-scanner ratings (disgust, fear, perceived infection risk and their mean) entered block-level mixed models with condition and block order as fixed effects and random intercepts and condition slopes per participant (maximum-likelihood fit), after centering each rating within participant and condition so that its term tests within-condition variation. Bootstrap intervals in Figure 3 used 2,000 resamples, and those in Figures 4 and S3, 5,000.

##### S1.10 Covariance-matched surrogates and composition checks

For each participant and condition, the concatenated windows (Schaefer 400, window at onset, normalization within condition, run 1) were replaced by 30 Gaussian surrogate series with the same $400 \times 400$ covariance matrix (Cholesky factor applied to independent standard normal samples of the same length, then z-scored) and no temporal structure. Synchrony-based and edge-wise edge metastability were computed on each surrogate and averaged. The surrogate contrast is the infection minus neutral difference of these averages. The observed minus surrogate contrast is the part of the condition difference that the static covariance does not reproduce. The same script (`composition_checks.py`) computes the mean pairwise correlation, $N\,\mathrm{Var}_t[m(t)]$, $\mathrm{Var}_t[q(t)]$ and $\mathrm{SD}_t[q(t)]$, the variance of the run-normalized parcel signals over the concatenated windows, the contrast on the strictly alternating segment (blocks 7 to 40) and the contrast with the 61 visual-network parcels removed, each tested with the 10,000-draw sign-flip permutation (Table S16). The within-block amplitude check (`recompute_schaefer400.py`, `amplitude_check.csv`) z-scores each parcel signal over the post-dummy run, extracts the same windows (onset, all blocks), and takes for each participant and condition the mean over parcels and windows of the within-window variance of these run-normalized signals; the infection minus neutral difference was tested with the paired $t$ test, without and with GSR.

##### S1.11 Control for evoked responses and nuisance signals

The script `control_analyses.py` re-estimates the synchrony-based and edge-wise contrasts with and without GSR (Schaefer 400, normalization within condition, all blocks, run 1) at shifts of 0 and 2 TR. Evoked-response removal subtracts, for each parcel and condition, the mean over that condition's 9-TR windows at each of the nine time points before the windows are concatenated and z-scored. This removes the stimulus-locked component of each condition. Nuisance regression is fitted within run on the 356 post-dummy volumes: linear and quadratic trends; 24 motion parameters (the six rigid-body parameters, their temporal derivatives and the squares of both, from the fMRIPrep confound file, with the undefined first derivative set to zero); and the mean white-matter and CSF signals. When GSR is applied, the global signal enters the same regression. Network-level values (Yeo seven networks, with the somatomotor and visual networks also split by hemisphere) are computed from the same z-scored windows with $R(t)$ restricted to the parcels of the network. Responses per block are the number of the three rating prompts answered, from the in-scanner rating logs. The across-run analysis expresses $\mathrm{cov}(\Delta\mathrm{EM}_1, \Delta\mathrm{EM}_2)$ as $\mathrm{cov}(I_1, I_2) + \mathrm{cov}(N_1, N_2) -$

$\mathrm{cov}(I_1, N_2) - \mathrm{cov}(N_1, I_2)$, where $I$ and $N$ are the infection and neutral levels and the subscript is the run.

To test the form of the effect that survived these steps, all variants were repeated in run 2 (main-text Figure 4), and the base and nuisance-regressed variants were repeated with the global signal fitted with separate coefficients for samples in infection windows, in neutral windows and elsewhere, and on the strictly alternating blocks 7 to 40; the contrast averaged over the two runs was tested in the 43 participants with both. In the condition-specific fit the mean-centered global signal is split into three regressors, each equal to it in one set of samples (the infection windows at the given shift, the neutral windows, or all remaining samples) and zero elsewhere; they replace the single global-signal regressor, alone or in the same regression as the nuisance signals, so that a condition difference in global-signal variance cannot be transferred to the residuals through a single coefficient. Table S21 gives the base and nuisance-regressed variants for run 1, run 2 and the pooled contrast, each tested with the 10,000-draw sign-flip permutation; participant-level values for all variants and both runs are in `control_analyses_subject.csv`.

### S2 Supplementary Results

#### S2.1 Stimulus low-level features

The two image sets were quantified post hoc on ten low-level features (Table S1). Six differed after FDR correction: the infection images had lower RMS contrast, entropy, mean hue, mid-spatial-frequency power and edge density, and more low-spatial-frequency power, than the neutral images. These differences cannot be removed from the present data. Under the pre-specified analysis the visual network showed a reduction of intermediate size ($d = -0.36$, $q = .07$; Table S14); after global signal regression its reduction was the largest of the seven networks ($d = -0.67$). Excluding the 61 visual parcels left the GSR contrasts almost unchanged (Table S16), so the reduction does not depend on visual cortex. A contribution of image statistics to the change within visual cortex itself is not excluded, and a replication with matched image statistics is needed.

Table S1: Low-level visual properties of the 20 infection and 20 neutral images (independent-samples t, BH-FDR across the ten features). sf: relative power in low, mid and high spatial-frequency bands.

| feature | infection M | SD | neutral M | SD | t | d | p (FDR) |
|---|---|---|---|---|---|---|---|
| luminance | 114.34 | 47.93 | 112.35 | 25.10 | 0.17 | 0.05 | 0.866 |
| rms contrast | 44.07 | 11.70 | 57.98 | 11.55 | -4.14 | -1.20 | 0.001 |
| hue mean | 44.78 | 34.09 | 68.16 | 26.98 | -2.58 | -0.76 | 0.029 |
| saturation mean | 100.95 | 34.55 | 98.37 | 45.85 | 0.23 | 0.06 | 0.866 |
| value mean | 145.53 | 59.27 | 135.13 | 21.26 | 0.75 | 0.23 | 0.574 |
| sf low | 0.92 | 0.04 | 0.88 | 0.06 | 2.57 | 0.71 | 0.029 |
| sf mid | 0.05 | 0.02 | 0.09 | 0.04 | -3.53 | -0.96 | 0.003 |
| sf high | 0.02 | 0.02 | 0.03 | 0.02 | -0.78 | -0.23 | 0.574 |
| edge density | 0.05 | 0.07 | 0.10 | 0.09 | -2.41 | -0.68 | 0.033 |
| entropy | 6.54 | 0.96 | 7.53 | 0.35 | -4.36 | -1.35 | 0.001 |

#### S2.2 Specification curve

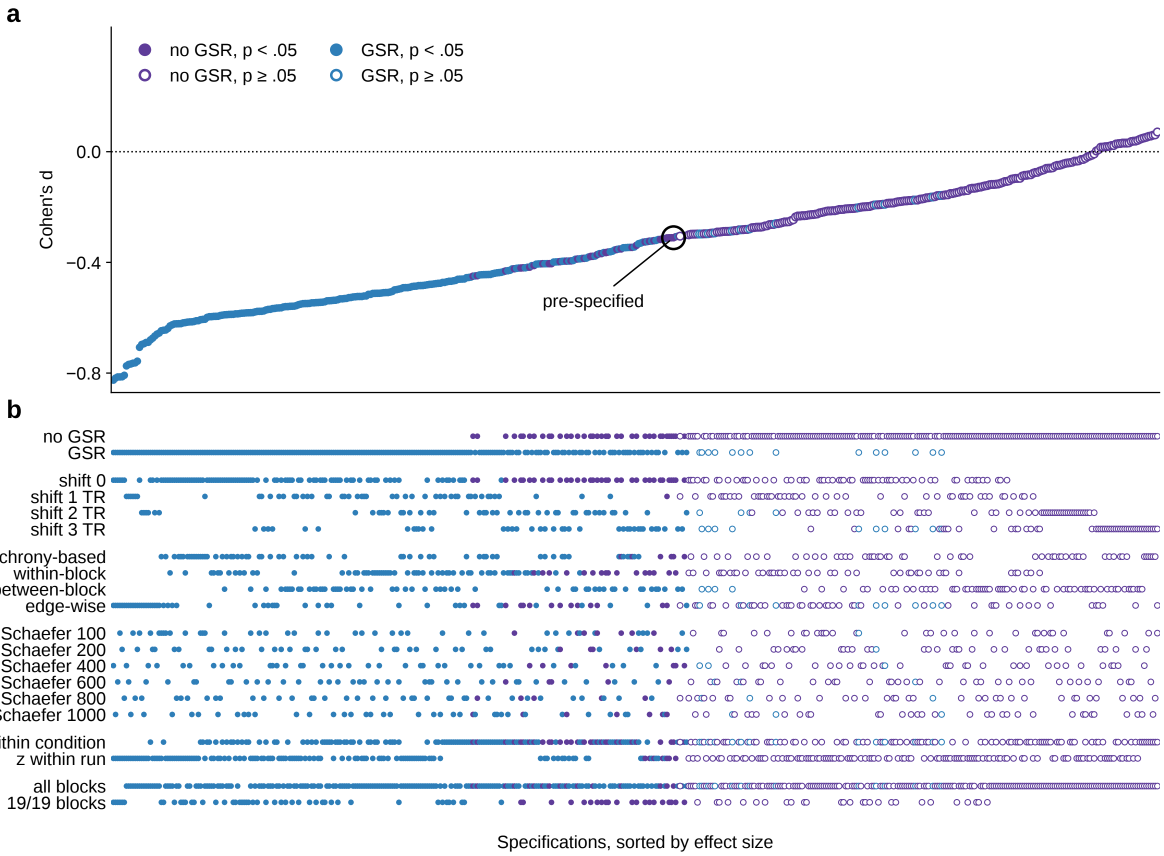

Figure S1: Full specification curve with indicator matrix. (a) Cohen's $d$ for the 480 specifications sorted by effect size; blue, GSR; purple, no GSR; filled, two-tailed permutation $p < .05$; the black ring marks the pre-specified analysis. (b) Indicator matrix: each column is one specification and the marked rows give its level on each dimension.

Table S2: All 480 specifications of the run 1 infection minus neutral contrast, sorted by Cohen's *d*. shift: window shift after onset in TR; z: normalization within condition (cond) or within run (run); parcels: Schaefer resolution; p from two-tailed sign-flip permutation.

| shift | definition | z | parcels | GSR | blocks | d | p (perm.) | n with reduction |
|---|---|---|---|---|---|---|---|---|
| 0 | edge-wise | run | 400 | yes | 19/19 | -0.825 | < .001 | 36 |
| 0 | edge-wise | run | 1000 | yes | 19/19 | -0.818 | < .001 | 37 |
| 0 | edge-wise | run | 600 | yes | 19/19 | -0.814 | < .001 | 36 |
| 0 | edge-wise | run | 100 | yes | 19/19 | -0.814 | < .001 | 34 |
| 0 | edge-wise | run | 200 | yes | 19/19 | -0.813 | < .001 | 36 |
| 0 | edge-wise | run | 800 | yes | 19/19 | -0.808 | < .001 | 36 |
| 1 | edge-wise | run | 400 | yes | all | -0.774 | < .001 | 37 |
| 1 | edge-wise | run | 600 | yes | all | -0.769 | < .001 | 37 |
| 1 | edge-wise | run | 1000 | yes | all | -0.767 | < .001 | 37 |
| 1 | edge-wise | run | 100 | yes | all | -0.764 | < .001 | 34 |
| 1 | edge-wise | run | 800 | yes | all | -0.763 | < .001 | 37 |
| 1 | edge-wise | run | 200 | yes | all | -0.757 | < .001 | 36 |
| 0 | edge-wise | run | 100 | yes | all | -0.707 | < .001 | 32 |
| 2 | edge-wise | run | 800 | yes | all | -0.697 | < .001 | 37 |
| 2 | edge-wise | run | 1000 | yes | all | -0.694 | < .001 | 36 |
| 2 | edge-wise | run | 600 | yes | all | -0.689 | < .001 | 37 |
| 2 | edge-wise | run | 400 | yes | all | -0.689 | < .001 | 37 |
| 0 | edge-wise | cond | 100 | yes | all | -0.680 | < .001 | 34 |
| 0 | edge-wise | run | 200 | yes | all | -0.674 | < .001 | 32 |
| 2 | edge-wise | run | 200 | yes | all | -0.666 | < .001 | 35 |
| 0 | edge-wise | run | 400 | yes | all | -0.659 | < .001 | 32 |
| 2 | edge-wise | run | 100 | yes | all | -0.655 | < .001 | 35 |
| 0 | synchrony-based | run | 100 | yes | 19/19 | -0.647 | < .001 | 32 |
| 0 | edge-wise | cond | 100 | yes | 19/19 | -0.645 | < .001 | 33 |
| 0 | synchrony-based | run | 100 | yes | all | -0.644 | < .001 | 32 |
| 0 | edge-wise | run | 600 | yes | all | -0.639 | < .001 | 32 |
| 0 | within-block | run | 100 | yes | all | -0.630 | < .001 | 33 |
| 0 | edge-wise | run | 1000 | yes | all | -0.626 | < .001 | 33 |
| 0 | synchrony-based | run | 200 | yes | 19/19 | -0.622 | < .001 | 31 |
| 0 | edge-wise | run | 800 | yes | all | -0.622 | < .001 | 33 |
| 0 | synchrony-based | run | 200 | yes | all | -0.622 | < .001 | 30 |
| 0 | synchrony-based | run | 800 | yes | 19/19 | -0.621 | < .001 | 31 |
| 0 | synchrony-based | run | 400 | yes | 19/19 | -0.619 | < .001 | 31 |
| 0 | within-block | run | 100 | yes | 19/19 | -0.618 | < .001 | 33 |
| 0 | synchrony-based | run | 800 | yes | all | -0.616 | < .001 | 30 |
| 0 | synchrony-based | run | 400 | yes | all | -0.615 | < .001 | 30 |
| 0 | synchrony-based | run | 1000 | yes | 19/19 | -0.615 | < .001 | 30 |
| 0 | synchrony-based | run | 600 | yes | 19/19 | -0.614 | < .001 | 31 |
| 0 | synchrony-based | run | 600 | yes | all | -0.611 | < .001 | 30 |

| shift | definition | z | parcels | GSR | blocks | d | p (perm.) | n with reduction |
|---|---|---|---|---|---|---|---|---|
| 0 | synchrony-based | run | 1000 | yes | all | -0.611 | < .001 | 30 |
| 0 | synchrony-based | cond | 100 | yes | all | -0.607 | < .001 | 32 |
| 0 | synchrony-based | cond | 100 | yes | 19/19 | -0.606 | < .001 | 32 |
| 1 | synchrony-based | run | 100 | yes | all | -0.605 | < .001 | 30 |
| 0 | synchrony-based | cond | 800 | yes | all | -0.599 | < .001 | 33 |
| 0 | edge-wise | cond | 200 | yes | all | -0.596 | < .001 | 32 |
| 0 | within-block | run | 200 | yes | all | -0.596 | < .001 | 33 |
| 0 | within-block | run | 400 | yes | all | -0.595 | < .001 | 32 |
| 0 | synchrony-based | cond | 800 | yes | 19/19 | -0.595 | < .001 | 31 |
| 0 | within-block | run | 1000 | yes | all | -0.595 | < .001 | 32 |
| 0 | within-block | run | 800 | yes | all | -0.592 | < .001 | 32 |
| 0 | synchrony-based | cond | 400 | yes | all | -0.591 | < .001 | 31 |
| 0 | between-block | run | 100 | yes | 19/19 | -0.589 | < .001 | 33 |
| 0 | synchrony-based | cond | 200 | yes | all | -0.589 | < .001 | 31 |
| 0 | within-block | run | 600 | yes | all | -0.588 | < .001 | 33 |
| 0 | synchrony-based | cond | 600 | yes | all | -0.588 | < .001 | 31 |
| 0 | synchrony-based | cond | 400 | yes | 19/19 | -0.588 | < .001 | 29 |
| 0 | within-block | run | 200 | yes | 19/19 | -0.586 | < .001 | 32 |
| 0 | synchrony-based | cond | 1000 | yes | all | -0.586 | < .001 | 32 |
| 0 | within-block | run | 400 | yes | 19/19 | -0.585 | < .001 | 31 |
| 0 | synchrony-based | cond | 200 | yes | 19/19 | -0.584 | < .001 | 30 |
| 0 | within-block | run | 1000 | yes | 19/19 | -0.583 | < .001 | 30 |
| 0 | synchrony-based | cond | 600 | yes | 19/19 | -0.582 | < .001 | 30 |
| 0 | synchrony-based | cond | 1000 | yes | 19/19 | -0.582 | < .001 | 30 |
| 0 | between-block | run | 100 | yes | all | -0.582 | < .001 | 32 |
| 0 | within-block | run | 800 | yes | 19/19 | -0.581 | < .001 | 32 |
| 3 | edge-wise | run | 1000 | yes | all | -0.579 | < .001 | 36 |
| 0 | within-block | run | 600 | yes | 19/19 | -0.577 | < .001 | 32 |
| 1 | synchrony-based | cond | 100 | yes | all | -0.577 | < .001 | 30 |
| 1 | synchrony-based | run | 200 | yes | all | -0.577 | < .001 | 27 |
| 3 | edge-wise | run | 800 | yes | all | -0.575 | < .001 | 36 |
| 0 | between-block | cond | 100 | yes | 19/19 | -0.572 | < .001 | 31 |
| 3 | edge-wise | run | 600 | yes | all | -0.570 | < .001 | 36 |
| 1 | synchrony-based | run | 400 | yes | all | -0.570 | < .001 | 29 |
| 3 | edge-wise | run | 400 | yes | all | -0.567 | < .001 | 35 |
| 0 | edge-wise | cond | 200 | yes | 19/19 | -0.566 | < .001 | 30 |
| 0 | edge-wise | cond | 400 | yes | all | -0.565 | < .001 | 33 |
| 1 | synchrony-based | run | 800 | yes | all | -0.565 | < .001 | 29 |
| 0 | between-block | run | 200 | yes | 19/19 | -0.564 | < .001 | 32 |
| 1 | synchrony-based | run | 600 | yes | all | -0.561 | < .001 | 28 |
| 0 | between-block | run | 400 | yes | 19/19 | -0.560 | < .001 | 30 |
| 0 | between-block | cond | 100 | yes | all | -0.560 | < .001 | 32 |
| 0 | between-block | run | 200 | yes | all | -0.559 | < .001 | 32 |

| shift | definition | z | parcels | GSR | blocks | d | p (perm.) | n with reduction |
|---|---|---|---|---|---|---|---|---|
| 0 | between-block | run | 800 | yes | 19/19 | -0.558 | < .001 | 33 |
| 1 | within-block | run | 100 | yes | all | -0.558 | < .001 | 27 |
| 1 | synchrony-based | run | 1000 | yes | all | -0.556 | < .001 | 29 |
| 0 | between-block | run | 600 | yes | 19/19 | -0.553 | < .001 | 31 |
| 0 | between-block | run | 400 | yes | all | -0.551 | < .001 | 31 |
| 1 | synchrony-based | cond | 400 | yes | all | -0.549 | < .001 | 28 |
| 3 | edge-wise | run | 200 | yes | all | -0.549 | < .001 | 35 |
| 1 | synchrony-based | cond | 200 | yes | all | -0.549 | < .001 | 29 |
| 0 | between-block | run | 1000 | yes | 19/19 | -0.548 | < .001 | 32 |
| 1 | synchrony-based | cond | 800 | yes | all | -0.546 | < .001 | 29 |
| 0 | between-block | run | 800 | yes | all | -0.546 | < .001 | 32 |
| 0 | between-block | cond | 200 | yes | 19/19 | -0.545 | < .001 | 33 |
| 3 | edge-wise | run | 100 | yes | all | -0.545 | < .001 | 35 |
| 0 | between-block | run | 600 | yes | all | -0.544 | < .001 | 31 |
| 0 | between-block | cond | 400 | yes | 19/19 | -0.544 | < .001 | 33 |
| 0 | between-block | cond | 800 | yes | 19/19 | -0.542 | < .001 | 33 |
| 1 | edge-wise | cond | 100 | yes | all | -0.539 | 0.001 | 31 |
| 1 | synchrony-based | cond | 600 | yes | all | -0.539 | < .001 | 27 |
| 0 | edge-wise | cond | 400 | yes | 19/19 | -0.538 | < .001 | 31 |
| 0 | between-block | run | 1000 | yes | all | -0.537 | < .001 | 32 |
| 0 | between-block | cond | 200 | yes | all | -0.537 | < .001 | 33 |
| 0 | between-block | cond | 600 | yes | 19/19 | -0.535 | < .001 | 33 |
| 0 | between-block | cond | 400 | yes | all | -0.532 | < .001 | 32 |
| 1 | within-block | run | 200 | yes | all | -0.532 | < .001 | 27 |
| 1 | synchrony-based | cond | 1000 | yes | all | -0.531 | < .001 | 27 |
| 0 | between-block | cond | 800 | yes | all | -0.530 | < .001 | 32 |
| 1 | within-block | run | 400 | yes | all | -0.527 | < .001 | 25 |
| 0 | between-block | cond | 1000 | yes | 19/19 | -0.527 | < .001 | 32 |
| 0 | between-block | cond | 600 | yes | all | -0.525 | < .001 | 32 |
| 2 | within-block | run | 100 | yes | all | -0.524 | 0.001 | 27 |
| 1 | within-block | run | 800 | yes | all | -0.523 | < .001 | 25 |
| 0 | edge-wise | cond | 600 | yes | all | -0.523 | < .001 | 31 |
| 1 | between-block | run | 100 | yes | all | -0.523 | < .001 | 31 |
| 1 | within-block | run | 600 | yes | all | -0.521 | < .001 | 25 |
| 1 | within-block | run | 1000 | yes | all | -0.521 | < .001 | 26 |
| 0 | within-block | cond | 400 | yes | all | -0.516 | < .001 | 29 |
| 0 | between-block | cond | 1000 | yes | all | -0.516 | < .001 | 31 |
| 2 | within-block | run | 800 | yes | all | -0.512 | 0.001 | 26 |
| 0 | within-block | cond | 100 | yes | all | -0.512 | < .001 | 30 |
| 0 | within-block | cond | 600 | yes | all | -0.512 | 0.001 | 30 |
| 2 | within-block | run | 400 | yes | all | -0.512 | 0.001 | 27 |
| 2 | within-block | run | 200 | yes | all | -0.511 | 0.001 | 28 |
| 2 | within-block | run | 1000 | yes | all | -0.509 | 0.001 | 26 |

| shift | definition | z | parcels | GSR | blocks | d | p (perm.) | n with reduction |
|---|---|---|---|---|---|---|---|---|
| 0 | within-block | cond | 800 | yes | all | -0.509 | < .001 | 30 |
| 2 | within-block | run | 600 | yes | all | -0.509 | 0.001 | 26 |
| 0 | within-block | cond | 1000 | yes | all | -0.507 | < .001 | 29 |
| 1 | between-block | cond | 100 | yes | all | -0.505 | < .001 | 31 |
| 0 | within-block | cond | 200 | yes | all | -0.499 | 0.001 | 30 |
| 1 | between-block | run | 200 | yes | all | -0.498 | 0.001 | 31 |
| 0 | edge-wise | cond | 600 | yes | 19/19 | -0.497 | 0.001 | 31 |
| 2 | synchrony-based | run | 100 | yes | all | -0.495 | 0.001 | 25 |
| 2 | synchrony-based | run | 800 | yes | all | -0.492 | 0.001 | 27 |
| 1 | between-block | run | 400 | yes | all | -0.491 | 0.002 | 31 |
| 3 | within-block | run | 100 | yes | all | -0.491 | 0.002 | 28 |
| 2 | synchrony-based | run | 200 | yes | all | -0.490 | 0.001 | 24 |
| 1 | between-block | run | 800 | yes | all | -0.487 | 0.002 | 30 |
| 3 | within-block | run | 800 | yes | all | -0.486 | 0.002 | 31 |
| 3 | within-block | run | 200 | yes | all | -0.486 | 0.002 | 27 |
| 3 | within-block | run | 1000 | yes | all | -0.484 | 0.002 | 31 |
| 2 | synchrony-based | run | 400 | yes | all | -0.484 | 0.002 | 25 |
| 3 | within-block | run | 400 | yes | all | -0.484 | 0.002 | 29 |
| 1 | between-block | run | 600 | yes | all | -0.482 | 0.002 | 30 |
| 0 | edge-wise | cond | 800 | yes | all | -0.481 | 0.002 | 30 |
| 2 | synchrony-based | run | 600 | yes | all | -0.480 | 0.002 | 26 |
| 3 | within-block | run | 600 | yes | all | -0.479 | 0.002 | 29 |
| 2 | synchrony-based | run | 1000 | yes | all | -0.478 | 0.002 | 26 |
| 1 | between-block | cond | 200 | yes | all | -0.477 | 0.002 | 32 |
| 0 | within-block | cond | 400 | yes | 19/19 | -0.476 | 0.002 | 29 |
| 1 | between-block | run | 1000 | yes | all | -0.475 | 0.002 | 29 |
| 0 | within-block | cond | 100 | yes | 19/19 | -0.472 | 0.003 | 28 |
| 1 | between-block | cond | 400 | yes | all | -0.472 | 0.002 | 30 |
| 0 | within-block | cond | 600 | yes | 19/19 | -0.470 | 0.002 | 29 |
| 0 | within-block | cond | 800 | yes | 19/19 | -0.468 | 0.003 | 29 |
| 1 | between-block | cond | 800 | yes | all | -0.468 | 0.002 | 30 |
| 0 | within-block | cond | 1000 | yes | 19/19 | -0.467 | 0.003 | 29 |
| 1 | edge-wise | cond | 200 | yes | all | -0.465 | 0.003 | 29 |
| 1 | between-block | cond | 600 | yes | all | -0.461 | 0.002 | 31 |
| 0 | edge-wise | cond | 1000 | yes | all | -0.461 | 0.003 | 30 |
| 0 | within-block | cond | 200 | yes | 19/19 | -0.460 | 0.003 | 28 |
| 0 | edge-wise | cond | 800 | yes | 19/19 | -0.460 | 0.003 | 31 |
| 2 | synchrony-based | cond | 800 | yes | all | -0.455 | 0.003 | 26 |
| 1 | within-block | cond | 100 | yes | all | -0.455 | 0.003 | 26 |
| 1 | within-block | cond | 400 | yes | all | -0.453 | 0.003 | 27 |
| 0 | edge-wise | cond | 1000 | no | all | -0.450 | 0.004 | 34 |
| 1 | between-block | cond | 1000 | yes | all | -0.449 | 0.003 | 30 |
| 0 | edge-wise | cond | 800 | no | all | -0.448 | 0.004 | 34 |

| shift | definition | z | parcels | GSR | blocks | d | p (perm.) | n with reduction |
|---|---|---|---|---|---|---|---|---|
| 2 | synchrony-based | cond | 400 | yes | all | -0.445 | 0.003 | 23 |
| 1 | within-block | cond | 600 | yes | all | -0.445 | 0.003 | 25 |
| 2 | synchrony-based | cond | 100 | yes | all | -0.444 | 0.003 | 26 |
| 2 | synchrony-based | cond | 200 | yes | all | -0.444 | 0.003 | 25 |
| 1 | within-block | cond | 800 | yes | all | -0.444 | 0.003 | 27 |
| 1 | within-block | cond | 200 | yes | all | -0.444 | 0.004 | 28 |
| 2 | synchrony-based | cond | 600 | yes | all | -0.441 | 0.003 | 24 |
| 1 | within-block | cond | 1000 | yes | all | -0.439 | 0.004 | 27 |
| 2 | synchrony-based | cond | 1000 | yes | all | -0.438 | 0.003 | 25 |
| 1 | edge-wise | cond | 400 | yes | all | -0.437 | 0.006 | 31 |
| 0 | edge-wise | cond | 1000 | yes | 19/19 | -0.436 | 0.006 | 31 |
| 3 | within-block | cond | 400 | yes | all | -0.433 | 0.007 | 28 |
| 0 | edge-wise | cond | 600 | no | all | -0.431 | 0.006 | 34 |
| 3 | within-block | cond | 1000 | yes | all | -0.430 | 0.007 | 27 |
| 2 | within-block | cond | 400 | yes | all | -0.428 | 0.007 | 27 |
| 3 | within-block | cond | 800 | yes | all | -0.424 | 0.007 | 27 |
| 0 | within-block | cond | 100 | no | all | -0.423 | 0.005 | 28 |
| 3 | within-block | cond | 600 | yes | all | -0.421 | 0.008 | 26 |
| 2 | within-block | cond | 600 | yes | all | -0.420 | 0.009 | 27 |
| 0 | edge-wise | cond | 800 | no | 19/19 | -0.420 | 0.008 | 32 |
| 0 | edge-wise | cond | 1000 | no | 19/19 | -0.420 | 0.008 | 33 |
| 2 | within-block | cond | 1000 | yes | all | -0.420 | 0.009 | 27 |
| 2 | within-block | cond | 800 | yes | all | -0.418 | 0.009 | 26 |
| 0 | edge-wise | cond | 400 | no | all | -0.417 | 0.007 | 32 |
| 3 | within-block | cond | 200 | yes | all | -0.412 | 0.010 | 26 |
| 0 | within-block | cond | 800 | no | all | -0.411 | 0.007 | 29 |
| 1 | edge-wise | cond | 600 | yes | all | -0.406 | 0.010 | 30 |
| 2 | within-block | cond | 200 | yes | all | -0.406 | 0.012 | 25 |
| 3 | synchrony-based | run | 800 | yes | all | -0.405 | 0.007 | 27 |
| 0 | within-block | cond | 400 | no | all | -0.405 | 0.007 | 29 |
| 3 | synchrony-based | run | 200 | yes | all | -0.405 | 0.007 | 25 |
| 2 | between-block | run | 100 | yes | all | -0.405 | 0.009 | 30 |
| 0 | within-block | cond | 600 | no | all | -0.405 | 0.008 | 29 |
| 0 | edge-wise | cond | 600 | no | 19/19 | -0.405 | 0.010 | 31 |
| 3 | synchrony-based | run | 400 | yes | all | -0.399 | 0.009 | 26 |
| 2 | within-block | cond | 100 | yes | all | -0.398 | 0.013 | 25 |
| 2 | between-block | run | 200 | yes | all | -0.398 | 0.009 | 28 |
| 0 | edge-wise | cond | 200 | no | all | -0.397 | 0.010 | 31 |
| 3 | synchrony-based | run | 1000 | yes | all | -0.396 | 0.008 | 27 |
| 3 | synchrony-based | run | 600 | yes | all | -0.395 | 0.009 | 26 |
| 0 | within-block | cond | 1000 | no | all | -0.395 | 0.010 | 28 |
| 3 | synchrony-based | run | 100 | yes | all | -0.395 | 0.009 | 25 |
| 0 | edge-wise | cond | 400 | no | 19/19 | -0.395 | 0.012 | 31 |

| shift | definition | z | parcels | GSR | blocks | d | p (perm.) | n with reduction |
|---|---|---|---|---|---|---|---|---|
| 2 | between-block | run | 800 | yes | all | -0.393 | 0.010 | 29 |
| 2 | between-block | run | 400 | yes | all | -0.389 | 0.011 | 27 |
| 0 | edge-wise | cond | 100 | no | all | -0.388 | 0.011 | 32 |
| 3 | within-block | cond | 100 | yes | all | -0.387 | 0.014 | 26 |
| 1 | edge-wise | cond | 800 | yes | all | -0.386 | 0.014 | 30 |
| 0 | within-block | cond | 100 | no | 19/19 | -0.385 | 0.009 | 29 |
| 2 | between-block | run | 600 | yes | all | -0.385 | 0.012 | 29 |
| 2 | between-block | run | 1000 | yes | all | -0.381 | 0.012 | 29 |
| 0 | edge-wise | cond | 200 | no | 19/19 | -0.378 | 0.015 | 30 |
| 0 | within-block | cond | 200 | no | all | -0.378 | 0.011 | 27 |
| 2 | between-block | cond | 100 | yes | all | -0.377 | 0.014 | 31 |
| 0 | edge-wise | cond | 100 | no | 19/19 | -0.372 | 0.015 | 30 |
| 2 | between-block | cond | 200 | yes | all | -0.369 | 0.016 | 30 |
| 0 | within-block | cond | 800 | no | 19/19 | -0.368 | 0.017 | 28 |
| 2 | between-block | cond | 800 | yes | all | -0.365 | 0.017 | 31 |
| 0 | within-block | cond | 400 | no | 19/19 | -0.364 | 0.016 | 28 |
| 0 | within-block | cond | 600 | no | 19/19 | -0.363 | 0.018 | 29 |
| 1 | edge-wise | cond | 1000 | yes | all | -0.362 | 0.021 | 30 |
| 2 | between-block | cond | 400 | yes | all | -0.360 | 0.019 | 28 |
| 2 | between-block | cond | 600 | yes | all | -0.355 | 0.019 | 30 |
| 0 | within-block | cond | 1000 | no | 19/19 | -0.353 | 0.021 | 28 |
| 3 | synchrony-based | cond | 800 | yes | all | -0.352 | 0.020 | 24 |
| 0 | synchrony-based | cond | 100 | no | all | -0.351 | 0.021 | 27 |
| 3 | synchrony-based | cond | 400 | yes | all | -0.347 | 0.022 | 24 |
| 2 | between-block | cond | 1000 | yes | all | -0.347 | 0.022 | 29 |
| 3 | synchrony-based | cond | 1000 | yes | all | -0.346 | 0.022 | 23 |
| 3 | synchrony-based | cond | 200 | yes | all | -0.346 | 0.021 | 24 |
| 0 | synchrony-based | cond | 100 | no | 19/19 | -0.346 | 0.025 | 27 |
| 3 | synchrony-based | cond | 600 | yes | all | -0.344 | 0.023 | 24 |
| 0 | within-block | cond | 200 | no | 19/19 | -0.338 | 0.025 | 29 |
| 3 | synchrony-based | cond | 100 | yes | all | -0.332 | 0.028 | 25 |
| 3 | between-block | run | 200 | yes | all | -0.330 | 0.028 | 27 |
| 3 | between-block | run | 100 | yes | all | -0.327 | 0.031 | 27 |
| 0 | within-block | run | 800 | no | 19/19 | -0.326 | 0.037 | 28 |
| 2 | edge-wise | cond | 100 | yes | all | -0.325 | 0.032 | 30 |
| 0 | within-block | run | 1000 | no | 19/19 | -0.323 | 0.038 | 27 |
| 3 | between-block | run | 800 | yes | all | -0.323 | 0.032 | 27 |
| 0 | within-block | run | 100 | no | 19/19 | -0.321 | 0.040 | 28 |
| 3 | between-block | run | 400 | yes | all | -0.320 | 0.034 | 26 |
| 3 | between-block | run | 600 | yes | all | -0.317 | 0.036 | 26 |
| 0 | synchrony-based | cond | 200 | no | all | -0.317 | 0.038 | 27 |
| 0 | edge-wise | run | 1000 | no | 19/19 | -0.314 | 0.043 | 32 |
| 3 | between-block | run | 1000 | yes | all | -0.314 | 0.038 | 27 |

| shift | definition | z | parcels | GSR | blocks | d | p (perm.) | n with reduction |
|---|---|---|---|---|---|---|---|---|
| 1 | edge-wise | run | 1000 | no | all | -0.312 | 0.043 | 30 |
| 0 | within-block | run | 600 | no | 19/19 | -0.311 | 0.046 | 27 |
| 0 | synchrony-based | cond | 200 | no | 19/19 | -0.311 | 0.042 | 27 |
| 0 | synchrony-based | cond | 400 | no | all | -0.311 | 0.042 | 27 |
| 0 | within-block | run | 400 | no | 19/19 | -0.307 | 0.050 | 27 |
| 3 | between-block | cond | 200 | yes | all | -0.305 | 0.044 | 30 |
| 1 | edge-wise | cond | 800 | no | all | -0.305 | 0.051 | 33 |
| 3 | between-block | cond | 100 | yes | all | -0.304 | 0.044 | 29 |
| 0 | synchrony-based | cond | 400 | no | 19/19 | -0.304 | 0.049 | 27 |
| 2 | edge-wise | cond | 200 | yes | all | -0.302 | 0.046 | 30 |
| 0 | within-block | run | 800 | no | all | -0.300 | 0.051 | 28 |
| 0 | synchrony-based | cond | 600 | no | all | -0.298 | 0.051 | 28 |
| 0 | within-block | run | 100 | no | all | -0.298 | 0.053 | 28 |
| 1 | edge-wise | cond | 1000 | no | all | -0.298 | 0.057 | 33 |
| 0 | edge-wise | run | 800 | no | 19/19 | -0.297 | 0.057 | 32 |
| 2 | edge-wise | cond | 400 | yes | all | -0.297 | 0.050 | 30 |
| 3 | between-block | cond | 800 | yes | all | -0.297 | 0.050 | 28 |
| 0 | synchrony-based | cond | 800 | no | all | -0.297 | 0.053 | 27 |
| 0 | within-block | run | 1000 | no | all | -0.296 | 0.055 | 27 |
| 3 | between-block | cond | 400 | yes | all | -0.296 | 0.050 | 29 |
| 1 | edge-wise | cond | 600 | no | all | -0.295 | 0.059 | 31 |
| 1 | edge-wise | run | 800 | no | all | -0.294 | 0.060 | 31 |
| 3 | between-block | cond | 600 | yes | all | -0.293 | 0.052 | 30 |
| 0 | synchrony-based | cond | 600 | no | 19/19 | -0.291 | 0.059 | 28 |
| 0 | within-block | run | 200 | no | 19/19 | -0.290 | 0.064 | 28 |
| 1 | within-block | run | 100 | no | all | -0.289 | 0.060 | 29 |
| 1 | within-block | cond | 100 | no | all | -0.288 | 0.055 | 27 |
| 1 | edge-wise | cond | 400 | no | all | -0.288 | 0.064 | 30 |
| 0 | synchrony-based | cond | 800 | no | 19/19 | -0.288 | 0.062 | 28 |
| 1 | within-block | run | 800 | no | all | -0.288 | 0.062 | 28 |
| 3 | between-block | cond | 1000 | yes | all | -0.286 | 0.058 | 28 |
| 1 | within-block | run | 1000 | no | all | -0.285 | 0.063 | 28 |
| 0 | within-block | run | 600 | no | all | -0.285 | 0.064 | 27 |
| 1 | edge-wise | cond | 200 | no | all | -0.282 | 0.067 | 27 |
| 2 | edge-wise | cond | 600 | yes | all | -0.282 | 0.063 | 29 |
| 0 | edge-wise | run | 600 | no | 19/19 | -0.282 | 0.073 | 31 |
| 0 | within-block | run | 400 | no | all | -0.281 | 0.069 | 28 |
| 0 | synchrony-based | cond | 1000 | no | all | -0.281 | 0.068 | 27 |
| 2 | edge-wise | cond | 800 | yes | all | -0.279 | 0.065 | 29 |
| 2 | edge-wise | run | 1000 | no | all | -0.275 | 0.072 | 28 |
| 1 | edge-wise | cond | 100 | no | all | -0.274 | 0.077 | 27 |
| 0 | synchrony-based | cond | 1000 | no | 19/19 | -0.273 | 0.077 | 28 |
| 1 | within-block | run | 600 | no | all | -0.273 | 0.073 | 28 |

| shift | definition | z | parcels | GSR | blocks | d | p (perm.) | n with reduction |
|---|---|---|---|---|---|---|---|---|
| 1 | edge-wise | run | 600 | no | all | -0.273 | 0.082 | 31 |
| 1 | within-block | run | 400 | no | all | -0.271 | 0.074 | 28 |
| 0 | edge-wise | run | 400 | no | 19/19 | -0.268 | 0.087 | 31 |
| 1 | synchrony-based | cond | 100 | no | all | -0.268 | 0.083 | 26 |
| 1 | within-block | cond | 800 | no | all | -0.264 | 0.081 | 26 |
| 1 | within-block | cond | 400 | no | all | -0.263 | 0.081 | 27 |
| 0 | within-block | run | 200 | no | all | -0.263 | 0.090 | 28 |
| 2 | edge-wise | cond | 1000 | yes | all | -0.260 | 0.087 | 29 |
| 1 | within-block | run | 200 | no | all | -0.260 | 0.086 | 29 |
| 1 | within-block | cond | 600 | no | all | -0.258 | 0.088 | 27 |
| 2 | edge-wise | run | 800 | no | all | -0.256 | 0.097 | 28 |
| 1 | within-block | cond | 1000 | no | all | -0.254 | 0.093 | 27 |
| 0 | edge-wise | run | 200 | no | 19/19 | -0.254 | 0.107 | 31 |
| 1 | edge-wise | run | 400 | no | all | -0.253 | 0.105 | 31 |
| 0 | edge-wise | run | 100 | no | 19/19 | -0.249 | 0.115 | 30 |
| 1 | within-block | cond | 200 | no | all | -0.246 | 0.105 | 27 |
| 1 | synchrony-based | cond | 200 | no | all | -0.236 | 0.127 | 26 |
| 2 | within-block | run | 800 | no | all | -0.232 | 0.124 | 29 |
| 0 | edge-wise | run | 1000 | no | all | -0.232 | 0.141 | 31 |
| 1 | edge-wise | run | 200 | no | all | -0.231 | 0.141 | 28 |
| 0 | between-block | cond | 100 | no | 19/19 | -0.231 | 0.139 | 27 |
| 0 | synchrony-based | run | 100 | no | 19/19 | -0.231 | 0.143 | 27 |
| 2 | within-block | run | 1000 | no | all | -0.229 | 0.131 | 28 |
| 3 | edge-wise | run | 1000 | no | all | -0.228 | 0.139 | 32 |
| 2 | edge-wise | run | 600 | no | all | -0.227 | 0.141 | 28 |
| 1 | synchrony-based | cond | 400 | no | all | -0.226 | 0.142 | 26 |
| 2 | within-block | run | 100 | no | all | -0.225 | 0.135 | 28 |
| 0 | edge-wise | run | 800 | no | all | -0.221 | 0.161 | 31 |
| 0 | between-block | cond | 100 | no | all | -0.220 | 0.159 | 27 |
| 0 | synchrony-based | run | 100 | no | all | -0.218 | 0.163 | 27 |
| 1 | edge-wise | run | 100 | no | all | -0.216 | 0.169 | 27 |
| 0 | edge-wise | run | 600 | no | all | -0.214 | 0.175 | 31 |
| 2 | within-block | run | 400 | no | all | -0.214 | 0.156 | 28 |
| 0 | edge-wise | run | 100 | no | all | -0.213 | 0.177 | 29 |
| 2 | within-block | run | 600 | no | all | -0.213 | 0.158 | 28 |
| 1 | synchrony-based | cond | 600 | no | all | -0.211 | 0.170 | 25 |
| 0 | edge-wise | run | 400 | no | all | -0.209 | 0.186 | 31 |
| 0 | synchrony-based | run | 200 | no | 19/19 | -0.209 | 0.182 | 27 |
| 0 | between-block | cond | 200 | no | 19/19 | -0.208 | 0.182 | 27 |
| 1 | synchrony-based | cond | 800 | no | all | -0.206 | 0.180 | 26 |
| 2 | within-block | run | 200 | no | all | -0.206 | 0.172 | 27 |
| 0 | synchrony-based | run | 400 | no | 19/19 | -0.206 | 0.187 | 26 |
| 0 | edge-wise | run | 200 | no | all | -0.205 | 0.194 | 30 |

| shift | definition | z | parcels | GSR | blocks | d | p (perm.) | n with reduction |
|---|---|---|---|---|---|---|---|---|
| 3 | edge-wise | run | 800 | no | all | -0.205 | 0.189 | 31 |
| 2 | within-block | cond | 100 | no | all | -0.204 | 0.186 | 27 |
| 3 | edge-wise | cond | 100 | yes | all | -0.202 | 0.188 | 28 |
| 2 | edge-wise | run | 400 | no | all | -0.201 | 0.192 | 28 |
| 0 | between-block | cond | 400 | no | 19/19 | -0.199 | 0.203 | 27 |
| 0 | synchrony-based | run | 800 | no | 19/19 | -0.199 | 0.203 | 26 |
| 0 | between-block | cond | 200 | no | all | -0.198 | 0.201 | 27 |
| 0 | synchrony-based | run | 600 | no | 19/19 | -0.198 | 0.204 | 26 |
| 0 | synchrony-based | run | 200 | no | all | -0.195 | 0.213 | 27 |
| 0 | synchrony-based | run | 400 | no | all | -0.193 | 0.216 | 27 |
| 3 | edge-wise | cond | 200 | yes | all | -0.192 | 0.209 | 29 |
| 0 | synchrony-based | run | 1000 | no | 19/19 | -0.192 | 0.219 | 26 |
| 1 | synchrony-based | cond | 1000 | no | all | -0.191 | 0.210 | 25 |
| 0 | between-block | cond | 400 | no | all | -0.190 | 0.221 | 27 |
| 3 | edge-wise | cond | 400 | yes | all | -0.190 | 0.216 | 28 |
| 0 | synchrony-based | run | 800 | no | all | -0.187 | 0.234 | 28 |
| 0 | synchrony-based | run | 600 | no | all | -0.186 | 0.236 | 27 |
| 0 | between-block | cond | 600 | no | 19/19 | -0.186 | 0.234 | 27 |
| 2 | within-block | cond | 800 | no | all | -0.185 | 0.231 | 28 |
| 0 | between-block | cond | 800 | no | 19/19 | -0.183 | 0.241 | 27 |
| 3 | edge-wise | run | 600 | no | all | -0.181 | 0.241 | 30 |
| 2 | within-block | cond | 400 | no | all | -0.180 | 0.246 | 27 |
| 0 | synchrony-based | run | 1000 | no | all | -0.179 | 0.256 | 27 |
| 1 | synchrony-based | run | 100 | no | all | -0.178 | 0.251 | 25 |
| 0 | between-block | cond | 600 | no | all | -0.177 | 0.254 | 27 |
| 3 | within-block | run | 800 | no | all | -0.177 | 0.249 | 24 |
| 3 | within-block | run | 1000 | no | all | -0.177 | 0.248 | 24 |
| 0 | between-block | cond | 800 | no | all | -0.175 | 0.261 | 27 |
| 3 | edge-wise | cond | 600 | yes | all | -0.175 | 0.261 | 29 |
| 2 | within-block | cond | 1000 | no | all | -0.175 | 0.257 | 27 |
| 2 | within-block | cond | 600 | no | all | -0.172 | 0.266 | 26 |
| 0 | between-block | cond | 1000 | no | 19/19 | -0.171 | 0.277 | 27 |
| 2 | edge-wise | run | 200 | no | all | -0.169 | 0.273 | 26 |
| 1 | between-block | cond | 100 | no | all | -0.167 | 0.284 | 26 |
| 2 | within-block | cond | 200 | no | all | -0.166 | 0.284 | 25 |
| 0 | between-block | run | 100 | no | 19/19 | -0.164 | 0.300 | 27 |
| 3 | edge-wise | cond | 800 | yes | all | -0.164 | 0.297 | 30 |
| 0 | between-block | cond | 1000 | no | all | -0.162 | 0.303 | 27 |
| 1 | synchrony-based | run | 200 | no | all | -0.159 | 0.303 | 26 |
| 3 | within-block | run | 600 | no | all | -0.158 | 0.302 | 24 |
| 3 | edge-wise | cond | 1000 | yes | all | -0.158 | 0.318 | 31 |
| 3 | within-block | run | 100 | no | all | -0.157 | 0.305 | 24 |
| 3 | within-block | run | 400 | no | all | -0.156 | 0.310 | 24 |

| shift | definition | z | parcels | GSR | blocks | d | p (perm.) | n with reduction |
|---|---|---|---|---|---|---|---|---|
| 3 | edge-wise | run | 400 | no | all | -0.156 | 0.315 | 27 |
| 1 | synchrony-based | run | 400 | no | all | -0.151 | 0.329 | 26 |
| 1 | between-block | cond | 200 | no | all | -0.151 | 0.328 | 27 |
| 0 | between-block | run | 100 | no | all | -0.149 | 0.346 | 28 |
| 0 | between-block | run | 200 | no | 19/19 | -0.147 | 0.358 | 27 |
| 3 | within-block | run | 200 | no | all | -0.146 | 0.345 | 24 |
| 1 | synchrony-based | run | 800 | no | all | -0.142 | 0.359 | 25 |
| 1 | synchrony-based | run | 600 | no | all | -0.142 | 0.360 | 26 |
| 1 | between-block | cond | 400 | no | all | -0.141 | 0.361 | 26 |
| 0 | between-block | run | 400 | no | 19/19 | -0.140 | 0.387 | 28 |
| 1 | synchrony-based | run | 1000 | no | all | -0.134 | 0.389 | 26 |
| 0 | between-block | run | 200 | no | all | -0.132 | 0.414 | 28 |
| 2 | edge-wise | run | 100 | no | all | -0.132 | 0.401 | 24 |
| 0 | between-block | run | 600 | no | 19/19 | -0.131 | 0.425 | 27 |
| 0 | between-block | run | 800 | no | 19/19 | -0.129 | 0.432 | 26 |
| 1 | between-block | cond | 600 | no | all | -0.128 | 0.410 | 26 |
| 0 | between-block | run | 400 | no | all | -0.125 | 0.443 | 28 |
| 1 | between-block | cond | 800 | no | all | -0.123 | 0.424 | 26 |
| 0 | between-block | run | 1000 | no | 19/19 | -0.122 | 0.457 | 27 |
| 1 | between-block | run | 100 | no | all | -0.119 | 0.455 | 26 |
| 2 | edge-wise | cond | 1000 | no | all | -0.118 | 0.445 | 30 |
| 3 | edge-wise | run | 200 | no | all | -0.118 | 0.449 | 25 |
| 2 | edge-wise | cond | 800 | no | all | -0.118 | 0.448 | 31 |
| 0 | between-block | run | 600 | no | all | -0.117 | 0.480 | 28 |
| 0 | between-block | run | 800 | no | all | -0.114 | 0.492 | 28 |
| 1 | between-block | cond | 1000 | no | all | -0.111 | 0.473 | 26 |
| 1 | between-block | run | 200 | no | all | -0.108 | 0.504 | 26 |
| 0 | between-block | run | 1000 | no | all | -0.107 | 0.525 | 28 |
| 2 | edge-wise | cond | 600 | no | all | -0.105 | 0.498 | 30 |
| 3 | within-block | cond | 800 | no | all | -0.100 | 0.522 | 23 |
| 1 | between-block | run | 400 | no | all | -0.098 | 0.545 | 26 |
| 3 | within-block | cond | 100 | no | all | -0.097 | 0.529 | 25 |
| 3 | within-block | cond | 1000 | no | all | -0.096 | 0.534 | 21 |
| 2 | edge-wise | cond | 400 | no | all | -0.096 | 0.535 | 30 |
| 1 | between-block | run | 600 | no | all | -0.088 | 0.586 | 25 |
| 1 | between-block | run | 800 | no | all | -0.086 | 0.598 | 25 |
| 3 | within-block | cond | 400 | no | all | -0.085 | 0.579 | 22 |
| 2 | edge-wise | cond | 200 | no | all | -0.084 | 0.588 | 29 |
| 3 | within-block | cond | 600 | no | all | -0.083 | 0.590 | 23 |
| 1 | between-block | run | 1000 | no | all | -0.078 | 0.635 | 25 |
| 2 | synchrony-based | cond | 100 | no | all | -0.075 | 0.623 | 22 |
| 3 | edge-wise | run | 100 | no | all | -0.075 | 0.638 | 24 |
| 3 | within-block | cond | 200 | no | all | -0.070 | 0.646 | 23 |

| shift | definition | z | parcels | GSR | blocks | d | p (perm.) | n with reduction |
|---|---|---|---|---|---|---|---|---|
| 2 | synchrony-based | cond | 200 | no | all | -0.068 | 0.659 | 23 |
| 2 | between-block | cond | 100 | no | all | -0.065 | 0.674 | 22 |
| 2 | between-block | cond | 200 | no | all | -0.060 | 0.696 | 22 |
| 2 | synchrony-based | cond | 400 | no | all | -0.060 | 0.698 | 24 |
| 2 | edge-wise | cond | 100 | no | all | -0.059 | 0.700 | 25 |
| 2 | synchrony-based | run | 100 | no | all | -0.058 | 0.707 | 23 |
| 2 | synchrony-based | run | 200 | no | all | -0.053 | 0.736 | 23 |
| 2 | between-block | cond | 400 | no | all | -0.050 | 0.750 | 23 |
| 2 | synchrony-based | cond | 600 | no | all | -0.049 | 0.748 | 22 |
| 2 | synchrony-based | cond | 800 | no | all | -0.047 | 0.763 | 24 |
| 2 | between-block | run | 100 | no | all | -0.044 | 0.781 | 22 |
| 2 | synchrony-based | run | 400 | no | all | -0.042 | 0.790 | 24 |
| 2 | between-block | run | 200 | no | all | -0.040 | 0.797 | 21 |
| 2 | between-block | cond | 600 | no | all | -0.039 | 0.800 | 22 |
| 2 | synchrony-based | cond | 1000 | no | all | -0.038 | 0.807 | 23 |
| 2 | between-block | cond | 800 | no | all | -0.034 | 0.826 | 24 |
| 2 | synchrony-based | run | 800 | no | all | -0.034 | 0.832 | 24 |
| 2 | synchrony-based | run | 600 | no | all | -0.033 | 0.836 | 24 |
| 2 | between-block | run | 400 | no | all | -0.028 | 0.858 | 23 |
| 2 | synchrony-based | run | 1000 | no | all | -0.028 | 0.859 | 24 |
| 2 | between-block | cond | 1000 | no | all | -0.024 | 0.880 | 22 |
| 2 | between-block | run | 600 | no | all | -0.018 | 0.907 | 23 |
| 2 | between-block | run | 800 | no | all | -0.016 | 0.923 | 23 |
| 3 | edge-wise | cond | 1000 | no | all | -0.012 | 0.935 | 28 |
| 2 | between-block | run | 1000 | no | all | -0.008 | 0.956 | 23 |
| 3 | edge-wise | cond | 800 | no | all | 0.003 | 0.986 | 28 |
| 3 | edge-wise | cond | 600 | no | all | 0.007 | 0.964 | 29 |
| 3 | between-block | run | 200 | no | all | 0.016 | 0.920 | 21 |
| 3 | edge-wise | cond | 400 | no | all | 0.016 | 0.922 | 27 |
| 3 | synchrony-based | run | 200 | no | all | 0.018 | 0.909 | 22 |
| 3 | between-block | run | 100 | no | all | 0.018 | 0.905 | 22 |
| 3 | synchrony-based | run | 100 | no | all | 0.020 | 0.900 | 23 |
| 3 | between-block | run | 400 | no | all | 0.022 | 0.886 | 21 |
| 3 | synchrony-based | run | 400 | no | all | 0.022 | 0.887 | 22 |
| 3 | between-block | cond | 200 | no | all | 0.028 | 0.854 | 21 |
| 3 | between-block | cond | 400 | no | all | 0.029 | 0.849 | 21 |
| 3 | synchrony-based | run | 600 | no | all | 0.030 | 0.855 | 24 |
| 3 | synchrony-based | run | 800 | no | all | 0.031 | 0.849 | 22 |
| 3 | between-block | cond | 100 | no | all | 0.031 | 0.840 | 22 |
| 3 | synchrony-based | run | 1000 | no | all | 0.031 | 0.847 | 24 |
| 3 | between-block | run | 600 | no | all | 0.032 | 0.836 | 21 |
| 3 | between-block | cond | 600 | no | all | 0.037 | 0.809 | 22 |
| 3 | between-block | run | 800 | no | all | 0.038 | 0.807 | 22 |

| shift | definition | z | parcels | GSR | blocks | d | p (perm.) | n with reduction |
|---|---|---|---|---|---|---|---|---|
| 3 | edge-wise | cond | 200 | no | all | 0.038 | 0.818 | 25 |
| 3 | between-block | run | 1000 | no | all | 0.041 | 0.793 | 21 |
| 3 | between-block | cond | 800 | no | all | 0.045 | 0.770 | 21 |
| 3 | between-block | cond | 1000 | no | all | 0.048 | 0.758 | 22 |
| 3 | synchrony-based | cond | 400 | no | all | 0.050 | 0.746 | 19 |
| 3 | synchrony-based | cond | 200 | no | all | 0.052 | 0.738 | 19 |
| 3 | synchrony-based | cond | 100 | no | all | 0.055 | 0.724 | 20 |
| 3 | synchrony-based | cond | 600 | no | all | 0.057 | 0.716 | 20 |
| 3 | synchrony-based | cond | 1000 | no | all | 0.060 | 0.704 | 21 |
| 3 | synchrony-based | cond | 800 | no | all | 0.061 | 0.697 | 20 |
| 3 | edge-wise | cond | 100 | no | all | 0.071 | 0.683 | 23 |

Table S3: Joint inference over the specification curve in each run (480 specifications, 10,000 sign-flip draws shared across specifications; run 1, n = 44; run 2, n = 43). Within each draw and for the observed counts, a specification counted as significant when its paired $t$ test gave $p < .05$.

| run | statistic | observed | p |
|---|---|---|---|
| 1 | median d | -0.341 | $< .001$ |
| 1 | number significant | 263 | $< .001$ |
| 1 | number significant and negative | 263 | $< .001$ |
| 1 | sum of significant d | -131.4 | $< .001$ |
| 2 | median d | -0.174 | 0.019 |
| 2 | number significant | 38 | 0.210 |
| 2 | number significant and negative | 38 | 0.102 |
| 2 | sum of significant d | -12.8 | 0.097 |

##### S2.3 Individual differences across specifications and runs

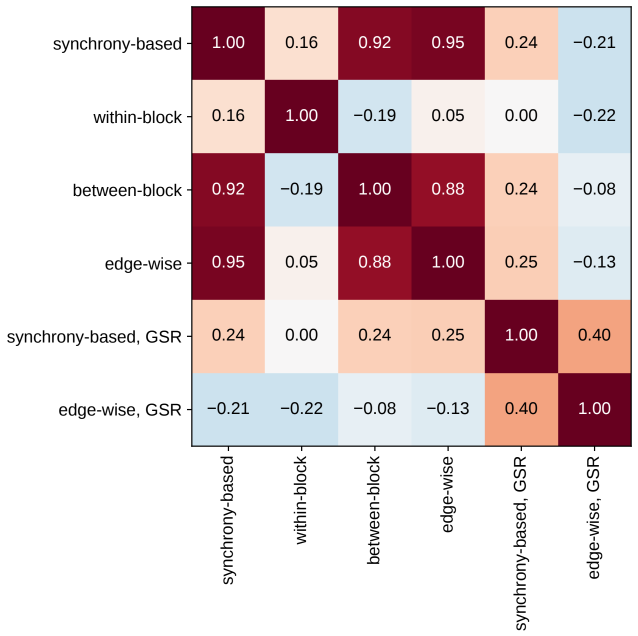

Figure S2: Between-subject Pearson correlation of $\Delta$EM (run 1, window at onset, Schaefer 400) across six specifications. Rankings of participants under specifications with and without global signal regression are almost unrelated to each other.

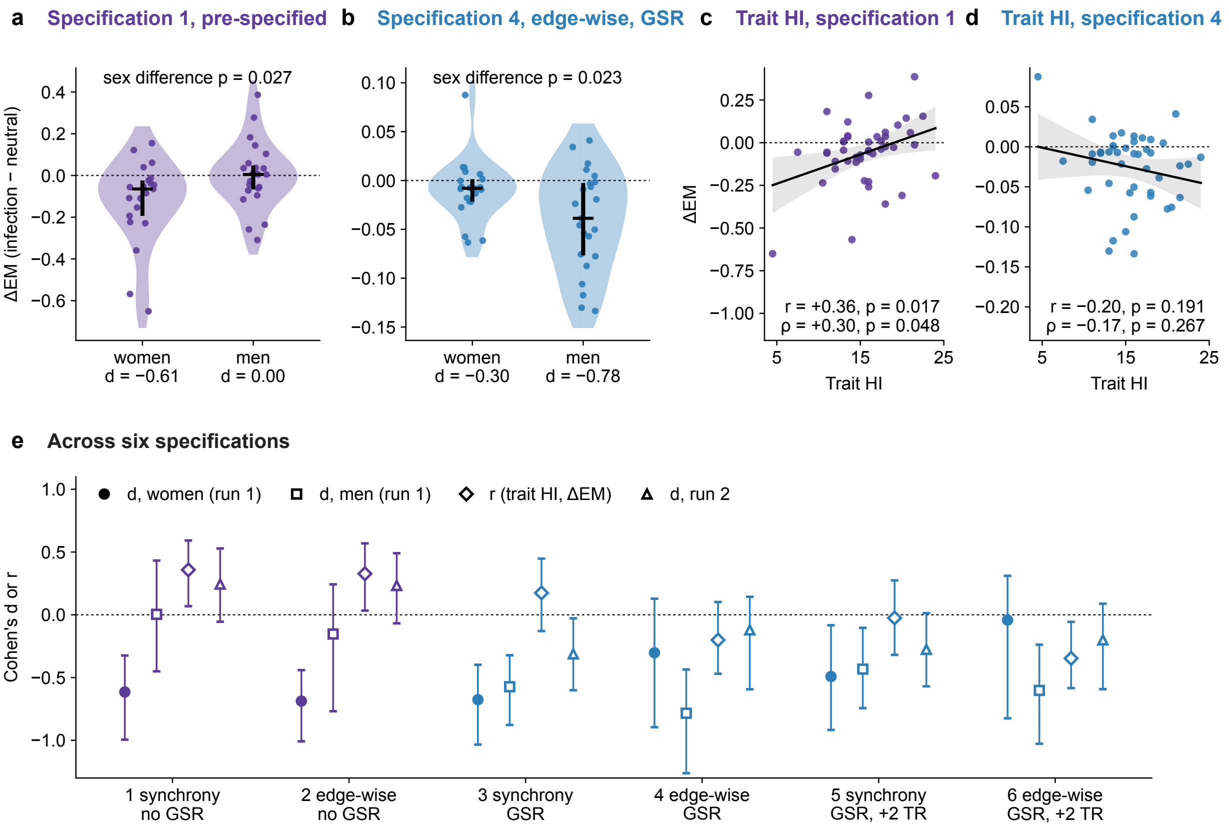

Figure S3: Individual differences depend on the specification. (a, b) ΔEM by sex under the pre-specified analysis (a) and the edge-wise GSR specification (b); note the different scales. (c, d) Trait horizontal individualism (mean of pre and post) against ΔEM under the same two specifications, with the least-squares line and its 95% band. (e) Sex-stratified effect sizes, the trait HI correlation, and the run 2 effect ($n = 43$) across six specifications (Schaefer 400; windows at onset unless a shift is stated), with bootstrap or Fisher 95% intervals. Specifications 1 to 6 in the panel titles are numbered as in panel e: specification 1 is the pre-specified analysis and specification 4 the edge-wise GSR specification.

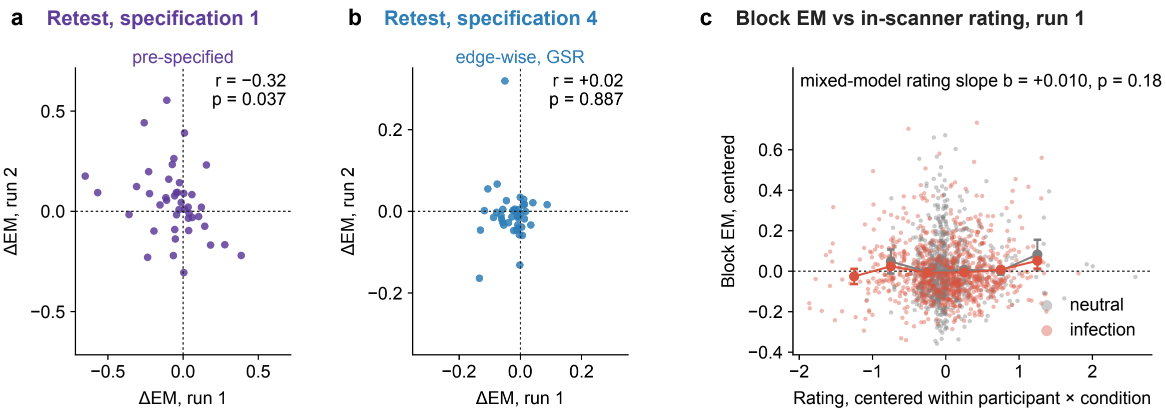

Figure S4: The condition contrast does not replicate across the two runs. (a, b) ΔEM in run 2 against ΔEM in run 1 under the pre-specified analysis (a; specification 1 of Figure S3e) and the edge-wise GSR specification (b; specification 4), with Pearson r ($n = 43$). (c) Block-level edge metastability (window-wise standard deviation of $R(t)$, run 1, centered within participant and condition) against the block's mean in-scanner rating, centered the same way; small points are blocks, large points and bars are bin means with standard errors, and the mixed-model slope is given.

#### S2.4 Motion

Table S4: Motion checks for the GSR specifications (run 1). r(ΔFD): correlation between the condition difference in framewise displacement and ΔEM; d, low-motion: effect after excluding the highest-motion quartile (n = 33); last column: p for the effect re-estimated at zero motion difference by regression.

| parcels | definition | shift | d | p | r(ΔFD) | p (r) | d, low-motion | p (low) | p at ΔFD = 0 |
|---|---|---|---|---|---|---|---|---|---|
| 200 | synchrony-based | 0 | -0.589 | < .001 | 0.099 | 0.524 | -0.606 | < .001 | 0.002 |
| 200 | synchrony-based | 2 | -0.444 | 0.003 | 0.199 | 0.195 | -0.428 | 0.016 | 0.026 |
| 200 | edge-wise | 0 | -0.596 | < .001 | 0.307 | 0.042 | -0.623 | < .001 | 0.004 |
| 200 | edge-wise | 2 | -0.302 | 0.046 | 0.332 | 0.028 | -0.264 | 0.131 | 0.259 |
| 400 | synchrony-based | 0 | -0.591 | < .001 | 0.107 | 0.488 | -0.618 | < .001 | 0.002 |
| 400 | synchrony-based | 2 | -0.445 | 0.003 | 0.216 | 0.160 | -0.438 | 0.014 | 0.028 |
| 400 | edge-wise | 0 | -0.565 | < .001 | 0.328 | 0.030 | -0.585 | 0.001 | 0.007 |
| 400 | edge-wise | 2 | -0.297 | 0.050 | 0.332 | 0.027 | -0.270 | 0.122 | 0.271 |

Table S5: Block-level linear mixed models (run 1, 1,716 windows, random condition slopes per participant) and motion checks. The dependent variable is the metastability of each 9-TR window on its own (standard deviation of $R(t)$ after z-scoring within the window), which is related to the within-block component of the pre-specified measure but is not the same quantity. Order: block order in units of ten blocks; FD: block-wise framewise displacement; both centered.

| model | term | beta | z | p |
|---|---|---|---|---|
| condition + order | condition | -0.021 | -3.015 | 0.003 |
| condition + order | order | 0.013 | 4.275 | $<$ .001 |
| condition + order + FD | condition | -0.019 | -2.650 | 0.008 |
| condition + order + FD | order | 0.010 | 3.336 | $<$ .001 |
| condition + order + FD | FD | 0.377 | 7.092 | $<$ .001 |
| condition × sex + order + FD | condition | -0.021 | -2.177 | 0.030 |
| condition × sex + order + FD | female | -0.034 | -1.741 | 0.082 |
| condition × sex + order + FD | condition × female | 0.005 | 0.381 | 0.703 |
| condition × sex + order + FD | order | 0.010 | 3.342 | $<$ .001 |
| condition × sex + order + FD | FD | 0.375 | 7.058 | $<$ .001 |
| motion checks | FD, infection minus neutral (paired t) | -0.007 | -2.507 | 0.016 |
| motion checks | r(ΔFD, ΔEM) | -0.342 | | 0.023 |
| motion checks | r(mean FD, ΔEM) | 0.210 | | 0.171 |
| motion checks | primary test without the highest-FD quartile (n = 33) | -0.429 | | 0.016 |

#### S2.5 Decomposition and sex

Table S6: Decomposition of the pre-specified measure (Schaefer 400, normalization within condition, no GSR) by sex at window shifts of 0 and 2 TR.

| shift | definition | d (all) | p (perm.) | d (women) | d (men) | t (sex) | p (sex) |
|---|---|---|---|---|---|---|---|
| 0 | synchrony-based | -0.311 | 0.042 | -0.615 | 0.004 | -2.285 | 0.027 |
| 0 | within-block | -0.405 | 0.007 | -0.210 | -0.576 | 1.308 | 0.198 |
| 0 | between-block | -0.190 | 0.221 | -0.721 | 0.155 | -2.610 | 0.012 |
| 0 | edge-wise | -0.417 | 0.007 | -0.687 | -0.153 | -2.043 | 0.047 |
| 2 | synchrony-based | -0.060 | 0.698 | -0.341 | 0.208 | -1.835 | 0.074 |
| 2 | within-block | -0.180 | 0.246 | 0.014 | -0.393 | 1.246 | 0.220 |
| 2 | between-block | -0.050 | 0.750 | -0.618 | 0.312 | -2.847 | 0.007 |
| 2 | edge-wise | -0.096 | 0.535 | -0.238 | 0.067 | -1.076 | 0.288 |

Table S7: Secondary tests as specified in the plan (linear mixed models, Holm-corrected), with BIC-approximated Bayes factors, and descriptive sex-stratified and run 2 effects. For the sex term the main text reports the equivalent Student's $t$ comparison of ΔEM between sexes ($t(42) = -2.28$, $p = .027$, Holm $p = .055$), because the Wald $z$ of the mixed model ignores the loss of degrees of freedom.

| test | beta | z | p | BF01 (BIC) | p (Holm) | d | n |
|---|---|---|---|---|---|---|---|
| condition × sex (run 1) | -0.124 | -2.338 | 0.019 | 0.713 | 0.039 | | |
| condition × run | 0.102 | 1.838 | 0.066 | 2.491 | 0.066 | | |
| primary contrast, women (descriptive) | -0.124 | | 0.006 | | | -0.615 | 21 |
| primary contrast, men (descriptive) | 0.001 | | 0.986 | | | 0.004 | 23 |
| primary contrast, run 2 (descriptive) | 0.043 | | 0.121 | | | 0.245 | 43 |

#### S2.6 Run 2 and individual differences by specification

Table S8: Run 1 and run 2 effects, sex-stratified effects, and the trait horizontal individualism (HI) correlation under 16 specifications (normalization within condition, all blocks; run 1, n = 44; run 2, n = 43).

| run | parcels | GSR | shift | definition | d | p (perm.) | d (women) | d (men) | p (sex) | r (HI) | p (HI) | $\rho$ (HI) |
|---|---|---|---|---|---|---|---|---|---|---|---|---|
| 1 | 200 | yes | 0 | synchrony-based | -0.589 | < .001 | -0.698 | -0.566 | 0.464 | 0.176 | 0.254 | -0.012 |
| 1 | 200 | yes | 0 | edge-wise | -0.596 | < .001 | -0.397 | -0.781 | 0.023 | -0.146 | 0.344 | -0.155 |
| 1 | 200 | yes | 2 | synchrony-based | -0.444 | 0.003 | -0.502 | -0.432 | 0.566 | -0.027 | 0.862 | -0.149 |
| 1 | 200 | yes | 2 | edge-wise | -0.302 | 0.046 | -0.042 | -0.601 | 0.104 | -0.327 | 0.030 | -0.173 |
| 1 | 200 | no | 0 | synchrony-based | -0.317 | 0.038 | -0.624 | 0.015 | 0.020 | 0.349 | 0.020 | 0.299 |
| 1 | 200 | no | 0 | edge-wise | -0.397 | 0.010 | -0.662 | -0.116 | 0.033 | 0.335 | 0.026 | 0.266 |
| 1 | 200 | no | 2 | synchrony-based | -0.068 | 0.659 | -0.342 | 0.216 | 0.067 | -0.203 | 0.187 | -0.160 |
| 1 | 200 | no | 2 | edge-wise | -0.084 | 0.588 | -0.243 | 0.115 | 0.219 | -0.244 | 0.111 | -0.173 |
| 1 | 400 | yes | 0 | synchrony-based | -0.591 | < .001 | -0.676 | -0.573 | 0.463 | 0.174 | 0.258 | 0.020 |
| 1 | 400 | yes | 0 | edge-wise | -0.565 | < .001 | -0.302 | -0.784 | 0.023 | -0.201 | 0.191 | -0.171 |
| 1 | 400 | yes | 2 | synchrony-based | -0.445 | 0.003 | -0.492 | -0.432 | 0.611 | -0.025 | 0.873 | -0.118 |
| 1 | 400 | yes | 2 | edge-wise | -0.297 | 0.050 | -0.042 | -0.602 | 0.111 | -0.347 | 0.021 | -0.196 |
| 1 | 400 | no | 0 | synchrony-based | -0.311 | 0.042 | -0.615 | 0.004 | 0.027 | 0.358 | 0.017 | 0.300 |
| 1 | 400 | no | 0 | edge-wise | -0.417 | 0.007 | -0.687 | -0.153 | 0.047 | 0.327 | 0.030 | 0.253 |
| 1 | 400 | no | 2 | synchrony-based | -0.060 | 0.698 | -0.341 | 0.208 | 0.074 | -0.182 | 0.236 | -0.128 |
| 1 | 400 | no | 2 | edge-wise | -0.096 | 0.535 | -0.238 | 0.067 | 0.288 | -0.253 | 0.097 | -0.214 |
| 2 | 200 | yes | 0 | synchrony-based | -0.319 | 0.043 | -0.407 | -0.286 | 0.841 | -0.145 | 0.354 | -0.118 |
| 2 | 200 | yes | 0 | edge-wise | -0.108 | 0.541 | -0.554 | -0.031 | 0.648 | -0.028 | 0.857 | -0.133 |
| 2 | 200 | yes | 2 | synchrony-based | -0.281 | 0.075 | -0.295 | -0.283 | 0.666 | -0.159 | 0.309 | -0.170 |
| 2 | 200 | yes | 2 | edge-wise | -0.182 | 0.245 | -0.200 | -0.194 | 0.632 | -0.017 | 0.915 | -0.082 |
| 2 | 200 | no | 0 | synchrony-based | 0.249 | 0.113 | 0.445 | 0.108 | 0.357 | -0.074 | 0.639 | -0.043 |
| 2 | 200 | no | 0 | edge-wise | 0.247 | 0.120 | 0.407 | 0.146 | 0.581 | -0.148 | 0.344 | -0.115 |
| 2 | 200 | no | 2 | synchrony-based | -0.033 | 0.834 | 0.251 | -0.388 | 0.052 | 0.003 | 0.987 | 0.125 |
| 2 | 200 | no | 2 | edge-wise | -0.019 | 0.900 | 0.301 | -0.328 | 0.046 | -0.056 | 0.724 | 0.122 |
| 2 | 400 | yes | 0 | synchrony-based | -0.312 | 0.048 | -0.392 | -0.281 | 0.838 | -0.130 | 0.406 | -0.116 |
| 2 | 400 | yes | 0 | edge-wise | -0.120 | 0.474 | -0.532 | -0.039 | 0.638 | -0.029 | 0.854 | -0.078 |
| 2 | 400 | yes | 2 | synchrony-based | -0.276 | 0.081 | -0.296 | -0.275 | 0.691 | -0.149 | 0.340 | -0.155 |
| 2 | 400 | yes | 2 | edge-wise | -0.200 | 0.195 | -0.185 | -0.223 | 0.565 | -0.018 | 0.907 | -0.054 |
| 2 | 400 | no | 0 | synchrony-based | 0.245 | 0.121 | 0.436 | 0.106 | 0.369 | -0.052 | 0.742 | -0.033 |
| 2 | 400 | no | 0 | edge-wise | 0.233 | 0.138 | 0.394 | 0.140 | 0.626 | -0.133 | 0.396 | -0.090 |
| 2 | 400 | no | 2 | synchrony-based | -0.034 | 0.828 | 0.254 | -0.382 | 0.051 | 0.001 | 0.994 | 0.129 |

| run | parcels | GSR | shift | definition | d | p (perm.) | d (women) | d (men) | p (sex) | r (HI) | p (HI) | $\rho$ (HI) |
|---|---|---|---|---|---|---|---|---|---|---|---|---|
| 2 | 400 | no | 2 | edge-wise | -0.029 | 0.861 | 0.314 | -0.331 | 0.041 | -0.067 | 0.672 | 0.117 |

#### S2.7 Change-score reliability

Table S9: Internal consistency, pre–post correlation and change-score reliability of the individualism–collectivism scales (n = 44). Reliability of change = $(\alpha_{pre} + \alpha_{post} - 2r)/(2 - 2r)$. IND, COL: 8-item totals; HI, VI, HC, VC: 4-item horizontal and vertical subscales. The formula gives a negative value whenever the pre–post correlation exceeds the mean internal consistency; such estimates are shown as $\leq 0$.

| scale | items | $\alpha$ pre | $\alpha$ post | r pre–post | reliability of change | SD change | r (change, ΔEM) | p |
|---|---|---|---|---|---|---|---|---|
| IND | 8 | 0.509 | 0.595 | 0.826 | $\leq 0$ | 3.400 | -0.065 | 0.675 |
| HI | 4 | 0.595 | 0.684 | 0.894 | $\leq 0$ | 1.865 | -0.027 | 0.860 |
| VI | 4 | 0.623 | 0.731 | 0.888 | $\leq 0$ | 2.026 | -0.084 | 0.588 |
| COL | 8 | 0.614 | 0.720 | 0.923 | $\leq 0$ | 2.459 | -0.082 | 0.595 |
| HC | 4 | 0.635 | 0.714 | 0.904 | $\leq 0$ | 1.605 | -0.215 | 0.161 |
| VC | 4 | 0.638 | 0.668 | 0.906 | $\leq 0$ | 1.764 | 0.081 | 0.602 |

#### S2.8 Trait predictors

Table S10: Trait predictors of ΔEM (pre-specified analysis, run 1): Pearson r with max-statistic permutation family-wise p over the 12 predictors, and Spearman $\rho$.

| predictor | r | p (raw) | p (FWER) | $\rho$ | p ($\rho$) |
|---|---|---|---|---|---|
| IND_pre | 0.192 | 0.211 | 0.745 | 0.145 | 0.347 |
| IND_post | 0.145 | 0.346 | 0.905 | 0.078 | 0.615 |
| HI_pre | 0.350 | 0.020 | 0.131 | 0.292 | 0.054 |
| HI_post | 0.347 | 0.021 | 0.138 | 0.292 | 0.054 |
| VI_pre | -0.093 | 0.547 | 0.988 | -0.042 | 0.785 |
| VI_post | -0.121 | 0.435 | 0.958 | -0.082 | 0.596 |
| COL_pre | -0.063 | 0.684 | 0.999 | -0.053 | 0.732 |
| COL_post | -0.088 | 0.570 | 0.991 | -0.088 | 0.572 |
| HC_pre | -0.082 | 0.596 | 0.994 | -0.123 | 0.425 |
| HC_post | -0.172 | 0.265 | 0.821 | -0.188 | 0.221 |
| VC_pre | -0.016 | 0.918 | 1 | 0.065 | 0.675 |
| VC_post | 0.019 | 0.902 | 1 | 0.051 | 0.742 |

Table S11: Sensitivity of the horizontal individualism (HI) correlation to analytic choices. HI was the dimension of prior interest within the exploratory trait family and is reported for that reason; it did not survive the family-wise correction in Table S10. For the leave-one-out row, r and r (max) are the smallest and largest of the 44 correlations.

| analysis | r | p | CI low | CI high | r (max) | n |
|---|---|---|---|---|---|---|
| HI mean of pre and post vs ΔEM | 0.358 | 0.017 | | | | |
| HI pre vs ΔEM | 0.350 | 0.020 | | | | |
| HI post vs ΔEM | 0.347 | 0.021 | | | | |
| HI mean: Spearman | 0.300 | 0.048 | | | | |
| HI mean: bootstrap 95% CI | | | -0.054 | 0.651 | | |
| HI mean: leave-one-out range | 0.187 | | | | 0.420 | |
| HI mean: partial, controlling mean FD | 0.342 | 0.023 | | | | |
| HI mean: partial, controlling sex | 0.366 | 0.015 | | | | |
| HI mean, women | 0.413 | 0.062 | | | | 21 |
| HI mean, men | 0.295 | 0.172 | | | | 23 |
| HI mean vs ΔEM: within-block, shift 0 | 0.015 | 0.922 | | | | |
| HI mean vs ΔEM: between-block, shift 0 | 0.373 | 0.013 | | | | |
| HI mean vs ΔEM: edge-wise, shift 0 | 0.327 | 0.030 | | | | |
| HI mean vs ΔEM: synchrony-based, shift 2 | -0.182 | 0.236 | | | | |
| HI mean vs ΔEM: within-block, shift 2 | -0.404 | 0.006 | | | | |
| HI mean vs ΔEM: between-block, shift 2 | 0.083 | 0.593 | | | | |
| HI mean vs ΔEM: edge-wise, shift 2 | -0.253 | 0.097 | | | | |
| HI mean vs ΔEM, run 2 | -0.052 | 0.742 | | | | 43 |

#### S2.9 Stability across runs and in-scanner ratings

Table S12: Stability between the two consecutive runs (runs 1 and 2; n = 43, the participants with both runs; Schaefer 400, normalization within condition, all blocks) of the condition contrast ΔEM and of the two condition levels, under eight specifications.

| GSR | definition | shift | n | r (ΔEM) | p | $\rho$ (ΔEM) | r (EM, infection) | r (EM, neutral) |
|---|---|---|---|---|---|---|---|---|
| no | synchrony-based | 0 | 43 | -0.319 | 0.037 | -0.373 | 0.485 | 0.457 |
| no | synchrony-based | 2 | 43 | 0.030 | 0.849 | -0.047 | 0.504 | 0.570 |
| no | edge-wise | 0 | 43 | -0.432 | 0.004 | -0.391 | 0.365 | 0.329 |
| no | edge-wise | 2 | 43 | 0.088 | 0.574 | 0.041 | 0.398 | 0.536 |
| yes | synchrony-based | 0 | 43 | -0.107 | 0.496 | 0.227 | 0.433 | 0.646 |
| yes | synchrony-based | 2 | 43 | 0.000 | 1.000 | 0.131 | 0.454 | 0.664 |
| yes | edge-wise | 0 | 43 | 0.022 | 0.887 | 0.059 | 0.153 | 0.374 |
| yes | edge-wise | 2 | 43 | 0.066 | 0.676 | -0.027 | 0.197 | 0.350 |

Table S13: In-scanner ratings and block-level edge metastability (window-wise standard deviation of $R(t)$, no GSR). Mixed models of block EM on condition, block order and the block's rating centered within participant and condition (random intercepts and condition slopes); r (subject): correlation between a participant's infection minus neutral rating difference and $\Delta$EM. Run 1: n = 44; run 2: n = 42, because participant 1's run 2 was excluded and participant 5 has no run 2 rating log.

| run | rating | blocks | b (rating) | p (rating) | b (condition) | p (condition) | r (subject) | p (r) |
|---|---|---|---|---|---|---|---|---|
| 1 | disgust | 1660 | 0.006 | 0.326 | -0.021 | 0.003 | 0.136 | 0.380 |
| 1 | fear | 1708 | 0.008 | 0.244 | -0.022 | 0.002 | 0.239 | 0.118 |
| 1 | perceived infection risk | 1707 | 0.007 | 0.255 | -0.021 | 0.002 | 0.054 | 0.730 |
| 1 | mean of three | 1715 | 0.010 | 0.181 | -0.021 | 0.002 | 0.162 | 0.292 |
| 2 | disgust | 1566 | 0.003 | 0.652 | 0.004 | 0.598 | -0.040 | 0.800 |
| 2 | fear | 1606 | -0.007 | 0.343 | 0.003 | 0.742 | 0.019 | 0.905 |
| 2 | perceived infection risk | 1613 | 0.004 | 0.582 | 0.002 | 0.834 | 0.047 | 0.766 |
| 2 | mean of three | 1618 | 0.001 | 0.930 | 0.002 | 0.786 | 0.015 | 0.924 |

#### S2.10 Network-level and parcel-level statistics

Table S14 gives within-network synchrony-based edge metastability differences for the seven Yeo networks under the pre-specified analysis and after global signal regression, at window shifts of 0 and 2 TR, with BH-FDR across the seven networks. Parcel-wise statistics for the edge-wise definition (400 parcels, BH-FDR across parcels) are provided as a CSV file in the data release (`node_stats_edgewise_S1.csv`); the maps are shown in main Figure 2d,e.

Table S14: Within-network synchrony-based edge metastability change (run 1, n = 44) by specification. shift: window shift after onset in TR; q: BH-FDR across the seven networks.

| GSR | shift | network | d | t | p | $q$ (FDR) |
|---|---|---|---|---|---|---|
| no | 0 | visual | -0.363 | -2.405 | 0.021 | 0.072 |
| no | 0 | somatomotor | -0.511 | -3.390 | 0.002 | 0.011 |
| no | 0 | dorsal attention | -0.186 | -1.232 | 0.225 | 0.287 |
| no | 0 | ventral attention | -0.317 | -2.106 | 0.041 | 0.096 |
| no | 0 | limbic | -0.045 | -0.298 | 0.767 | 0.767 |
| no | 0 | control | -0.177 | -1.176 | 0.246 | 0.287 |
| no | 0 | default mode | -0.238 | -1.581 | 0.121 | 0.212 |
| no | 2 | visual | -0.036 | -0.238 | 0.813 | 0.909 |
| no | 2 | somatomotor | -0.250 | -1.658 | 0.105 | 0.732 |
| no | 2 | dorsal attention | 0.135 | 0.894 | 0.376 | 0.909 |
| no | 2 | ventral attention | -0.103 | -0.680 | 0.500 | 0.909 |
| no | 2 | limbic | -0.045 | -0.296 | 0.768 | 0.909 |
| no | 2 | control | 0.087 | 0.580 | 0.565 | 0.909 |
| no | 2 | default mode | -0.017 | -0.115 | 0.909 | 0.909 |
| yes | 0 | visual | -0.665 | -4.411 | $< .001$ | $< .001$ |
| yes | 0 | somatomotor | -0.353 | -2.340 | 0.024 | 0.028 |
| yes | 0 | dorsal attention | -0.388 | -2.572 | 0.014 | 0.019 |
| yes | 0 | ventral attention | -0.509 | -3.379 | 0.002 | 0.005 |
| yes | 0 | limbic | -0.050 | -0.332 | 0.741 | 0.741 |
| yes | 0 | control | -0.415 | -2.754 | 0.009 | 0.019 |
| yes | 0 | default mode | -0.394 | -2.615 | 0.012 | 0.019 |
| yes | 2 | visual | -0.530 | -3.518 | 0.001 | 0.007 |
| yes | 2 | somatomotor | -0.207 | -1.372 | 0.177 | 0.297 |
| yes | 2 | dorsal attention | -0.281 | -1.862 | 0.069 | 0.162 |
| yes | 2 | ventral attention | -0.474 | -3.146 | 0.003 | 0.011 |
| yes | 2 | limbic | 0.020 | 0.132 | 0.896 | 0.896 |
| yes | 2 | control | -0.191 | -1.267 | 0.212 | 0.297 |
| yes | 2 | default mode | -0.161 | -1.071 | 0.290 | 0.339 |

#### S2.11 Power

Table S15: Sensitivity power analysis (paired design, two-tailed $\alpha = .05$).

| quantity | value |
|---|---|
| minimum detectable d, n = 44, $\alpha = .05$ two-tailed, power = .80 | 0.43 |
| n required for d = 0.30, $\alpha = .05$ two-tailed, power = .80 | 90 |
| achieved power for d = 0.30 at n = 44 | 0.49 |

##### S2.12 Composition of the contrast

Table S16: Composition of the run 1 contrast (Schaefer 400, window at onset, normalization within condition, all blocks unless stated; n = 44). Alternating segment: blocks 7 to 40, 17 blocks per condition. Means are over participants; r rows give the correlation across participants between the observed and surrogate ΔEM; difference is infection minus neutral, tested with the 10,000-draw sign-flip permutation on Cohen's *d*. Surrogates: 30 covariance-matched Gaussian series per participant and condition. The within-block parcel variance row is the amplitude check defined in Section S1.10 (mean over parcels and windows of the within-window variance of the run-normalized signals); its p is from the paired *t* test.

| GSR | quantity | infection M | neutral M | difference or r | d | p (perm.) |
|---|---|---|---|---|---|---|
| no | synchrony-based EM | 0.471 | 0.530 | -0.059 | -0.311 | 0.042 |
| no | edge-wise EM | 1.085 | 1.124 | -0.039 | -0.417 | 0.007 |
| no | mean pairwise correlation | 0.299 | 0.305 | -0.005 | -0.082 | 0.602 |
| no | $N\,\mathrm{Var}_t[m(t)]$ | 120.473 | 122.597 | -2.124 | -0.082 | 0.602 |
| no | $\mathrm{Var}_t[q(t)]$ | 0.409 | 0.499 | -0.090 | -0.311 | 0.041 |
| no | $\mathrm{SD}_t[q(t)]$ | 0.589 | 0.665 | -0.076 | -0.460 | 0.003 |
| no | surrogate synchrony-based EM | 0.422 | 0.425 | -0.002 | -0.024 | 0.875 |
| no | surrogate edge-wise EM | 1.075 | 1.071 | 0.004 | 0.177 | 0.259 |
| no | parcel variance, concatenated windows (run-norm.) | 0.969 | 0.980 | -0.011 | -0.091 | 0.570 |
| no | SD over parcels of parcel variance | 0.106 | 0.105 | 0.001 | 0.069 | 0.647 |
| no | $\mathrm{SD}_t[q(t)]$, run-norm. signals | 0.586 | 0.665 | -0.078 | -0.284 | 0.070 |
| no | synchrony-based EM, alternating segment | 0.469 | 0.523 | -0.054 | -0.267 | 0.087 |
| no | between-block component, alternating segment | 0.295 | 0.321 | -0.026 | -0.125 | 0.423 |
| no | edge-wise EM, alternating segment | 1.086 | 1.118 | -0.032 | -0.312 | 0.042 |
| no | synchrony-based EM without visual parcels | 0.486 | 0.546 | -0.060 | -0.311 | 0.043 |
| no | edge-wise EM without visual parcels | 1.089 | 1.130 | -0.041 | -0.420 | 0.007 |
| no | observed minus surrogate, synchrony-based | | | -0.056 | -0.418 | 0.006 |
| no | r(observed, surrogate ΔEM), synchrony-based | | | 0.743 | | |
| no | observed minus surrogate, edge-wise | | | -0.043 | -0.527 | < .001 |
| no | r(observed, surrogate ΔEM), edge-wise | | | 0.564 | | |
| no | within-block parcel variance (run-norm.) | 0.554 | 0.557 | -0.003 | -0.044 | 0.771 |
| yes | synchrony-based EM | 0.156 | 0.210 | -0.054 | -0.591 | < .001 |
| yes | edge-wise EM | 1.032 | 1.058 | -0.026 | -0.565 | < .001 |
| yes | mean pairwise correlation | 0.101 | 0.120 | -0.018 | -0.544 | 0.001 |
| yes | $N\,\mathrm{Var}_t[m(t)]$ | 41.362 | 48.729 | -7.367 | -0.544 | 0.001 |
| yes | $\mathrm{Var}_t[q(t)]$ | 0.197 | 0.263 | -0.067 | -0.530 | < .001 |
| yes | $\mathrm{SD}_t[q(t)]$ | 0.425 | 0.492 | -0.067 | -0.596 | < .001 |
| yes | surrogate synchrony-based EM | 0.143 | 0.168 | -0.024 | -0.514 | 0.002 |
| yes | surrogate edge-wise EM | 1.038 | 1.039 | -0.001 | -0.173 | 0.260 |
| yes | parcel variance, concatenated windows (run-norm.) | 0.959 | 0.999 | -0.041 | -0.580 | < .001 |
| yes | SD over parcels of parcel variance | 0.108 | 0.103 | 0.005 | 0.405 | 0.010 |
| yes | $\mathrm{SD}_t[q(t)]$, run-norm. signals | 0.406 | 0.503 | -0.097 | -0.642 | < .001 |
| yes | synchrony-based EM, alternating segment | 0.145 | 0.169 | -0.023 | -0.297 | 0.049 |
| yes | between-block component, alternating segment | 0.097 | 0.112 | -0.015 | -0.227 | 0.147 |
| yes | edge-wise EM, alternating segment | 1.031 | 1.047 | -0.015 | -0.324 | 0.039 |
| yes | synchrony-based EM without visual parcels | 0.174 | 0.223 | -0.049 | -0.559 | < .001 |
| yes | edge-wise EM without visual parcels | 1.035 | 1.060 | -0.025 | -0.535 | < .001 |
| yes | observed minus surrogate, synchrony-based | | | -0.030 | -0.496 | < .001 |
| yes | r(observed, surrogate ΔEM), synchrony-based | | | 0.807 | | |
| yes | observed minus surrogate, edge-wise | | | -0.024 | -0.552 | < .001 |
| yes | r(observed, surrogate ΔEM), edge-wise | | | 0.258 | | |

| GSR | quantity | infection M | neutral M | difference or r | d | p (perm.) |
|---|---|---|---|---|---|---|
| yes | within-block parcel variance (run-norm.) | 0.616 | 0.613 | 0.003 | 0.049 | 0.748 |

#### S2.13 Control for evoked responses and nuisance signals

Table S17: Control for evoked responses and nuisance signals (run 1, Schaefer 400, normalization within condition, all blocks, $n = 44$). Cells give Cohen's $d$ of $\Delta$EM and the two-tailed sign-flip permutation $p$ (10,000 draws, uncorrected; the rows re-estimate the same contrast under alternative preprocessing). With GSR, the global signal entered the same regression as the nuisance signals. The synchrony-based GSR column survives Holm–Bonferroni correction across the six variants at each shift.

| | | no GSR | | GSR | |
|---|---|---|---|---|---|
| shift | removed | synchrony-based | edge-wise | synchrony-based | edge-wise |
| 0 | none (as in the specification curve) | −0.31 (0.042) | −0.42 (0.007) | −0.59 (< .001) | −0.56 (< .001) |
| 0 | stimulus-locked response removed | −0.27 (0.083) | −0.38 (0.015) | −0.54 (< .001) | −0.50 (0.001) |
| 0 | linear and quadratic trends | −0.28 (0.067) | −0.27 (0.075) | −0.55 (< .001) | −0.19 (0.223) |
| 0 | 24 motion parameters | −0.33 (0.033) | −0.32 (0.035) | −0.42 (< .001) | −0.25 (0.107) |
| 0 | white matter and CSF | −0.10 (0.514) | −0.10 (0.520) | −0.35 (0.014) | −0.18 (0.250) |
| 0 | trends, motion, white matter and CSF | −0.11 (0.491) | 0.08 (0.608) | −0.47 (< .001) | −0.16 (0.300) |
| 0 | all nuisance steps and evoked response | 0.04 (0.818) | 0.23 (0.135) | −0.44 (0.002) | −0.04 (0.786) |
| 2 | none (as in the specification curve) | −0.06 (0.698) | −0.10 (0.535) | −0.45 (0.003) | −0.30 (0.050) |
| 2 | stimulus-locked response removed | −0.07 (0.672) | −0.06 (0.698) | −0.45 (0.003) | −0.20 (0.179) |
| 2 | linear and quadratic trends | 0.02 (0.911) | 0.10 (0.529) | −0.61 (< .001) | −0.09 (0.558) |
| 2 | 24 motion parameters | −0.10 (0.511) | −0.03 (0.871) | −0.45 (< .001) | −0.28 (0.061) |
| 2 | white matter and CSF | −0.11 (0.477) | −0.15 (0.322) | −0.41 (0.006) | −0.21 (0.164) |
| 2 | trends, motion, white matter and CSF | −0.13 (0.387) | 0.02 (0.912) | −0.57 (< .001) | −0.17 (0.270) |
| 2 | all nuisance steps and evoked response | −0.05 (0.763) | 0.19 (0.235) | −0.54 (< .001) | 0.00 (0.986) |

#### S2.14 Button responses and networks after nuisance regression

Table S18: Responses and ratings by condition from the in-scanner rating logs (paired $t$ tests; participant 5 has no run 2 rating log; the run 2 log of participant 1 is included, because only the image file of that run was affected). Participants responded with the right hand.

| run | quantity | n | infection | neutral | difference | t | p |
|---|---|---|---|---|---|---|---|
| 1 | responses per block (of 3) | 44 | 2.939 | 2.976 | -0.038 | -3.473 | 0.001 |
| 1 | mean rating (1–4) | 44 | 2.824 | 1.213 | 1.612 | 19.988 | < .001 |
| 2 | responses per block (of 3) | 43 | 2.916 | 2.934 | -0.017 | -1.582 | 0.121 |
| 2 | mean rating (1–4) | 43 | 2.854 | 1.181 | 1.674 | 19.994 | < .001 |

Table S19: Within-network synchrony-based ΔEM (run 1, onset) without and with the three nuisance steps (trends, 24 motion parameters, white matter and CSF), for the seven networks ($q$: BH-FDR across the seven applied to the paired $t$-test $p$, as in Table S14) and for the somatomotor and visual networks split by hemisphere.

| GSR | nuisance | network | d | p (perm.) | p ($t$) | $q$ (FDR) |
|---|---|---|---|---|---|---|
| no | no | visual | -0.363 | 0.017 | 0.021 | 0.072 |
| no | no | somatomotor | -0.511 | 0.001 | 0.002 | 0.011 |
| no | no | dorsal attention | -0.186 | 0.227 | 0.225 | 0.287 |
| no | no | ventral attention | -0.317 | 0.041 | 0.041 | 0.096 |
| no | no | limbic | -0.045 | 0.781 | 0.767 | 0.767 |
| no | no | control | -0.177 | 0.247 | 0.246 | 0.287 |
| no | no | default mode | -0.238 | 0.125 | 0.121 | 0.212 |
| no | no | somatomotor, left | -0.435 | 0.007 | 0.006 | |
| no | no | somatomotor, right | -0.507 | 0.001 | 0.002 | |
| no | no | somatomotor, left minus right | 0.084 | 0.587 | 0.580 | |
| no | no | visual, left | -0.408 | 0.006 | 0.010 | |
| no | no | visual, right | -0.352 | 0.024 | 0.024 | |
| no | no | visual, left minus right | -0.051 | 0.737 | 0.738 | |
| no | yes | visual | -0.441 | 0.005 | 0.005 | 0.038 |
| no | yes | somatomotor | -0.073 | 0.638 | 0.631 | 0.631 |
| no | yes | dorsal attention | 0.141 | 0.359 | 0.354 | 0.631 |
| no | yes | ventral attention | -0.252 | 0.103 | 0.101 | 0.355 |
| no | yes | limbic | 0.085 | 0.580 | 0.576 | 0.631 |
| no | yes | control | 0.122 | 0.482 | 0.422 | 0.631 |
| no | yes | default mode | -0.107 | 0.496 | 0.480 | 0.631 |
| no | yes | somatomotor, left | -0.070 | 0.648 | 0.644 | |
| no | yes | somatomotor, right | 0.003 | 0.984 | 0.983 | |
| no | yes | somatomotor, left minus right | -0.093 | 0.543 | 0.542 | |
| no | yes | visual, left | -0.598 | $< .001$ | $< .001$ | |
| no | yes | visual, right | -0.311 | 0.044 | 0.046 | |
| no | yes | visual, left minus right | -0.455 | 0.004 | 0.004 | |
| yes | no | visual | -0.665 | $< .001$ | $< .001$ | $< .001$ |
| yes | no | somatomotor | -0.353 | 0.023 | 0.024 | 0.028 |
| yes | no | dorsal attention | -0.388 | 0.010 | 0.014 | 0.019 |
| yes | no | ventral attention | -0.509 | 0.001 | 0.002 | 0.005 |
| yes | no | limbic | -0.050 | 0.748 | 0.741 | 0.741 |
| yes | no | control | -0.415 | 0.006 | 0.009 | 0.019 |
| yes | no | default mode | -0.394 | 0.008 | 0.012 | 0.019 |
| yes | no | somatomotor, left | -0.469 | 0.002 | 0.003 | |
| yes | no | somatomotor, right | -0.224 | 0.145 | 0.145 | |
| yes | no | somatomotor, left minus right | -0.191 | 0.214 | 0.211 | |
| yes | no | visual, left | -0.753 | $< .001$ | $< .001$ | |
| yes | no | visual, right | -0.618 | $< .001$ | $< .001$ | |
| yes | no | visual, left minus right | -0.104 | 0.502 | 0.492 | |
| yes | yes | visual | -0.550 | $< .001$ | $< .001$ | 0.005 |
| yes | yes | somatomotor | -0.141 | 0.361 | 0.355 | 0.414 |
| yes | yes | dorsal attention | -0.414 | 0.008 | 0.009 | 0.020 |

| GSR | nuisance | network | d | p (perm.) | p ($t$) | $q$ (FDR) |
|---|---|---|---|---|---|---|
| yes | yes | ventral attention | -0.505 | 0.002 | 0.002 | 0.006 |
| yes | yes | limbic | 0.052 | 0.736 | 0.730 | 0.730 |
| yes | yes | control | -0.209 | 0.183 | 0.174 | 0.243 |
| yes | yes | default mode | -0.242 | 0.117 | 0.116 | 0.202 |
| yes | yes | somatomotor, left | -0.160 | 0.295 | 0.296 | |
| yes | yes | somatomotor, right | -0.055 | 0.720 | 0.718 | |
| yes | yes | somatomotor, left minus right | -0.116 | 0.445 | 0.447 | |
| yes | yes | visual, left | -0.650 | < .001 | < .001 | |
| yes | yes | visual, right | -0.461 | 0.002 | 0.004 | |
| yes | yes | visual, left minus right | -0.356 | 0.024 | 0.023 | |

#### S2.15 Across-run structure of the contrast

Table S20: Across-run structure of the condition contrast (onset, Schaefer 400, normalization within condition, $n = 43$, the participants with both runs). $I$ and $N$ are the infection and neutral edge metastability levels, 1 and 2 the run, $\Delta = I - N$. The covariance of $\Delta_1$ and $\Delta_2$ is $\text{cov}(I_1, I_2) + \text{cov}(N_1, N_2) - \text{cov}(I_1, N_2) - \text{cov}(N_1, I_2)$; covariances and variances are in `retest_covariance.csv` in the data release.

| GSR | definition | r(Δ1, Δ2) | r(I1, I2) | r(N1, N2) | r(I1, N2) | r(N1, I2) | r(I1, N1) | r(I2, N2) |
|---|---|---|---|---|---|---|---|---|
| no | synchrony-based | -0.32 | 0.48 | 0.46 | 0.65 | 0.43 | 0.74 | 0.85 |
| no | edge-wise | -0.43 | 0.37 | 0.33 | 0.61 | 0.33 | 0.74 | 0.75 |
| yes | synchrony-based | -0.11 | 0.43 | 0.65 | 0.55 | 0.61 | 0.79 | 0.72 |
| yes | edge-wise | 0.02 | 0.15 | 0.37 | 0.19 | 0.30 | 0.68 | 0.69 |

##### S2.16 Replication in run 2, global-signal fit and block order

Table S21: Replication of the control analyses in run 2, with a condition-specific global-signal fit and on the strictly alternating blocks (Schaefer 400, normalization within condition). Cells give Cohen's $d$ of $\Delta$EM and the two-tailed sign-flip permutation $p$ (10,000 draws, uncorrected). Run 1: $n = 44$; run 2: $n = 43$, because participant 1's run 2 was excluded (the file exported as run 2 was a copy of run 1); pooled: the mean of the two runs' contrasts in the 43 participants with both runs. Blocks: all blocks, or the strictly alternating blocks 7 to 40 (17 per condition; at a 2-TR shift the final infection block falls outside the run, leaving 16). GSR: single, the global signal regressed with one coefficient per parcel, as in Table S17; by cond., the global signal fitted with separate coefficients for samples in infection windows, in neutral windows and elsewhere. Nuisance: linear and quadratic trends, 24 motion parameters and the white-matter and CSF signals, in the same regression as the global signal. Shift: window shift after onset in TR. The run 1 rows for all blocks with no or single GSR repeat Table S17.

| | | | shift 0 | | | shift 2 | | |
|---|---|---|---|---|---|---|---|---|
| blocks | GSR | nuisance | run 1 | run 2 | pooled | run 1 | run 2 | pooled |
| *synchrony-based* | | | | | | | | |
| all | none | no | −0.31 (0.042) | 0.24 (0.121) | −0.07 (0.652) | −0.06 (0.698) | −0.03 (0.828) | −0.05 (0.747) |
| all | none | yes | −0.11 (0.491) | −0.12 (0.440) | −0.16 (0.315) | −0.13 (0.387) | −0.06 (0.737) | −0.13 (0.404) |
| all | single | no | −0.59 (< .001) | −0.31 (0.048) | −0.67 (< .001) | −0.45 (0.003) | −0.28 (0.081) | −0.50 (0.001) |
| all | single | yes | −0.47 (< .001) | −0.44 (0.005) | −0.65 (< .001) | −0.57 (< .001) | −0.29 (0.063) | −0.60 (< .001) |
| all | by cond. | no | −0.52 (< .001) | −0.32 (0.047) | −0.59 (< .001) | −0.40 (0.007) | −0.34 (0.035) | −0.48 (< .001) |
| all | by cond. | yes | −0.47 (< .001) | −0.42 (0.005) | −0.63 (< .001) | −0.56 (< .001) | −0.35 (0.028) | −0.64 (< .001) |
| 7–40 | none | no | −0.27 (0.087) | 0.21 (0.188) | −0.07 (0.652) | −0.10 (0.513) | −0.04 (0.811) | −0.08 (0.623) |
| 7–40 | none | yes | −0.12 (0.461) | −0.10 (0.561) | −0.13 (0.418) | −0.19 (0.227) | −0.04 (0.811) | −0.14 (0.391) |
| 7–40 | single | no | −0.30 (0.049) | −0.22 (0.157) | −0.36 (0.023) | −0.09 (0.541) | −0.25 (0.113) | −0.27 (0.086) |
| 7–40 | single | yes | −0.36 (0.021) | −0.26 (0.095) | −0.38 (0.018) | −0.27 (0.080) | −0.12 (0.446) | −0.22 (0.151) |
| 7–40 | by cond. | no | −0.30 (0.050) | −0.17 (0.281) | −0.33 (0.036) | −0.13 (0.392) | −0.21 (0.184) | −0.26 (0.097) |
| 7–40 | by cond. | yes | −0.30 (0.052) | −0.23 (0.140) | −0.33 (0.036) | −0.31 (0.047) | −0.12 (0.426) | −0.25 (0.107) |
| *edge-wise* | | | | | | | | |
| all | none | no | −0.42 (0.007) | 0.23 (0.138) | −0.06 (0.714) | −0.10 (0.535) | −0.03 (0.861) | −0.06 (0.698) |
| all | none | yes | 0.08 (0.608) | 0.06 (0.737) | 0.08 (0.642) | 0.02 (0.912) | −0.05 (0.752) | −0.02 (0.880) |
| all | single | no | −0.56 (< .001) | −0.12 (0.474) | −0.42 (0.007) | −0.30 (0.050) | −0.20 (0.195) | −0.35 (0.027) |
| all | single | yes | −0.16 (0.300) | −0.03 (0.872) | −0.12 (0.466) | −0.17 (0.270) | −0.12 (0.444) | −0.19 (0.235) |
| all | by cond. | no | −0.28 (0.072) | −0.02 (0.928) | −0.18 (0.243) | −0.15 (0.340) | −0.14 (0.370) | −0.18 (0.239) |
| all | by cond. | yes | −0.10 (0.511) | −0.01 (0.963) | −0.07 (0.701) | −0.13 (0.404) | −0.12 (0.430) | −0.17 (0.284) |
| 7–40 | none | no | −0.31 (0.042) | 0.21 (0.173) | −0.03 (0.846) | −0.08 (0.592) | −0.05 (0.777) | −0.07 (0.679) |
| 7–40 | none | yes | 0.03 (0.856) | 0.01 (0.938) | 0.03 (0.882) | −0.05 (0.744) | −0.13 (0.422) | −0.13 (0.437) |
| 7–40 | single | no | −0.32 (0.038) | −0.04 (0.836) | −0.22 (0.170) | −0.10 (0.537) | −0.15 (0.329) | −0.17 (0.269) |
| 7–40 | single | yes | 0.00 (0.993) | −0.02 (0.916) | −0.01 (0.971) | −0.03 (0.826) | −0.15 (0.345) | −0.12 (0.450) |
| 7–40 | by cond. | no | −0.17 (0.275) | 0.05 (0.784) | −0.06 (0.670) | 0.03 (0.853) | −0.11 (0.482) | −0.06 (0.713) |
| 7–40 | by cond. | yes | 0.04 (0.800) | 0.00 (0.996) | 0.03 (0.876) | 0.00 (0.989) | −0.15 (0.325) | −0.11 (0.512) |

### References

[1] Anke Haberkamp, Julia A. Glombiewski, Frank Schmidt, and Antonia Barke. The DIsgust-RelaTed-Images (DIRTI) database: Validation of a novel standardized set of disgust pictures. *Behaviour Research and Therapy*, 89:86–94, 2017. doi: 10.1016/j.brat.2016.11.010.

[2] Harry C Triandis and Michele J Gelfand. Converging measurement of horizontal and vertical individualism and collectivism. *Journal of Personality and Social Psychology*, 74(1):118–128, 1998. doi: 10.1037/0022-3514.74.1.118.

[3] Lesley A Duncan, Mark Schaller, and Justin H Park. Perceived vulnerability to disease: Development and validation of a 15-item self-report instrument. *Personality and Individual Differences*, 47(6):541–546, 2009. doi: 10.1016/j.paid.2009.05.001.

[4] Jonathan Peirce, Jeremy R. Gray, Sol Simpson, Michael MacAskill, Richard Höchenberger, Hiroyuki Sogo, et al. PsychoPy2: Experiments in behavior made easy. *Behavior Research Methods*, 51(1):195–203, 2019. doi: 10.3758/s13428-018-01193-y.

[5] Oscar Esteban, Christopher Markiewicz, Ross W Blair, Craig Moodie, Ayse Ilkay Isik, Asier Erramuzpe Aliaga, et al. fMRIPrep: a robust preprocessing pipeline for functional MRI. *Nature Methods*, 16:111–116, 2019. doi: 10.1038/s41592-018-0235-4.

[6] Oscar Esteban, Ross Blair, Christopher J. Markiewicz, Shoshana L. Berleant, Craig Moodie, Feilong Ma, et al. fMRIPrep 24.1.1, 2024. Software.

[7] K. Gorgolewski, C. D. Burns, C. Madison, D. Clark, Y. O. Halchenko, M. L. Waskom, et al. Nipype: a flexible, lightweight and extensible neuroimaging data processing framework in python. *Frontiers in Neuroinformatics*, 5:13, 2011. doi: 10.3389/fninf.2011.00013.

[8] Krzysztof J. Gorgolewski, Oscar Esteban, Christopher J. Markiewicz, Erik Ziegler, David Gage Ellis, Michael Philipp Notter, et al. Nipype, 2018. Software.

[9] N. J. Tustison, B. B. Avants, P. A. Cook, Y. Zheng, A. Egan, P. A. Yushkevich, et al. N4itk: Improved n3 bias correction. *IEEE Transactions on Medical Imaging*, 29(6):1310–1320, 2010. ISSN 0278-0062. doi: 10.1109/TMI.2010.2046908.

[10] B.B. Avants, C.L. Epstein, M. Grossman, and J.C. Gee. Symmetric diffeomorphic image registration with cross-correlation: Evaluating automated labeling of elderly and neurodegenerative brain. *Medical Image Analysis*, 12(1):26–41, 2008. ISSN 1361-8415. doi: 10.1016/j.media.2007.06.004. URL `https://www.sciencedirect.com/science/article/pii/S1361841507000606`.

[11] Y. Zhang, M. Brady, and S. Smith. Segmentation of brain MR images through a hidden markov random field model and the expectation-maximization algorithm. *IEEE Transactions on Medical Imaging*, 20(1):45–57, 2001. ISSN 0278-0062. doi: 10.1109/42.906424.

[12] R. Ciric, William H. Thompson, R. Lorenz, M. Goncalves, E. MacNicol, C. J. Markiewicz, et al. TemplateFlow: FAIR-sharing of multi-scale, multi-species brain models. *Nature Methods*, 19: 1568–1571, 2022. doi: 10.1038/s41592-022-01681-2.

[13] VS Fonov, AC Evans, RC McKinstry, CR Almli, and DL Collins. Unbiased nonlinear average age-appropriate brain templates from birth to adulthood. *NeuroImage*, 47, Supplement 1:S102, 2009. doi: 10.1016/S1053-8119(09)70884-5.

[14] Mark Jenkinson, Peter Bannister, Michael Brady, and Stephen Smith. Improved optimization for the robust and accurate linear registration and motion correction of brain images. *NeuroImage*, 17(2):825–841, 2002. ISSN 1053-8119. doi: 10.1006/nimg.2002.1132. URL https://www.sciencedirect.com/science/article/pii/S1053811902911328.

[15] Mark Jenkinson and Stephen Smith. A global optimisation method for robust affine registration of brain images. *Medical Image Analysis*, 5(2):143–156, 2001. ISSN 1361-8415. doi: 10.1016/S1361-8415(01)00036-6. URL https://www.sciencedirect.com/science/article/pii/S1361841501000366.

[16] Douglas N Greve and Bruce Fischl. Accurate and robust brain image alignment using boundary-based registration. *NeuroImage*, 48(1):63–72, 2009. ISSN 1095-9572. doi: 10.1016/j.neuroimage.2009.06.060.

[17] Jonathan D. Power, Anish Mitra, Timothy O. Laumann, Abraham Z. Snyder, Bradley L. Schlaggar, and Steven E. Petersen. Methods to detect, characterize, and remove motion artifact in resting state fmri. *NeuroImage*, 84(Supplement C):320–341, 2014. ISSN 1053-8119. doi: 10.1016/j.neuroimage.2013.08.048. URL https://www.sciencedirect.com/science/article/pii/S1053811913009117.

[18] Yashar Behzadi, Khaled Restom, Joy Liau, and Thomas T. Liu. A component based noise correction method (CompCor) for BOLD and perfusion based fmri. *NeuroImage*, 37(1):90–101, 2007. ISSN 1053-8119. doi: 10.1016/j.neuroimage.2007.04.042. URL https://www.sciencedirect.com/science/article/pii/S1053811907003837.

[19] Theodore D. Satterthwaite, Mark A. Elliott, Raphael T. Gerraty, Kosha Ruparel, James Loughead, Monica E. Calkins, et al. An improved framework for confound regression and filtering for control of motion artifact in the preprocessing of resting-state functional connectivity data. *NeuroImage*, 64(1):240–256, 2013. ISSN 10538119. doi: 10.1016/j.neuroimage.2012.08.052. URL https://www.sciencedirect.com/science/article/pii/S1053811912008609.

[20] Rémi Patriat, Richard C. Reynolds, and Rasmus M. Birn. An improved model of motion-related signal changes in fMRI. *NeuroImage*, 144, Part A:74–82, January 2017. ISSN 1053-8119. doi: 10.1016/j.neuroimage.2016.08.051. URL https://www.sciencedirect.com/science/article/pii/S1053811916304360.

[21] Alexandre Abraham, Fabian Pedregosa, Michael Eickenberg, Philippe Gervais, Andreas Mueller, Jean Kossaifi, et al. Machine learning for neuroimaging with scikit-learn. *Frontiers in Neuroinformatics*, 8, 2014. ISSN 1662-5196. doi: 10.3389/fninf.2014.00014. URL https://www.frontiersin.org/articles/10.3389/fninf.2014.00014/full.

[22] Alexander Schaefer, Ru Kong, Evan M. Gordon, Timothy O. Laumann, Xi-Nian Zuo, Avram J. Holmes, et al. Local-global parcellation of the human cerebral cortex from intrinsic functional connectivity mri. *Cerebral Cortex*, 28(9):3095–3114, 2018. doi: 10.1093/cercor/bhx179.

[23] B. T. Thomas Yeo, Fenna M. Krienen, Jorge Sepulcre, Mert R. Sabuncu, Danial Lashkari, Marisa Hollinshead, et al. The organization of the human cerebral cortex estimated by intrinsic functional connectivity. *Journal of Neurophysiology*, 106(3):1125–1165, 2011. doi: 10.1152/jn.00338.2011.

[24] Uri Simonsohn, Joseph P Simmons, and Leif D Nelson. Specification curve analysis. *Nature Human Behaviour*, 4(11):1208–1214, 2020. doi: 10.1038/s41562-020-0912-z.
